# MOFF2: A Transferable Coarse-Grained Protein Force Field for Predictive Condensate Simulations

**DOI:** 10.64898/2026.06.10.731384

**Authors:** Shuming Liu, Yumeng Zhang, Ivan Riveros, Cong Wang, Bin Zhang

**Author notes:** Contributed equally to this work (Shuming Liu, Yumeng Zhang).

## Abstract

Coarse-grained protein force fields enable simulations of biomolecular systems at length and time scales that are difficult to access with atomistic models, but achieving transferability across folded, intrinsically disordered, and multidomain proteins remains challenging. A central difficulty is that one-bead-per-residue models must represent chemically specific residue interactions while also absorbing solvent-mediated and many-body effects into a simplified energy function. Here, we present MOFF2, a transferable coarse-grained protein force field that combines residue-pair-specific interactions with a density-dependent many-body potential. MOFF2 is optimized using a two-stage strategy: bottom-up parameter learning from heterogeneous reference ensembles followed by refinement against experimental conformational observables. The resulting model provides balanced performance across ordered proteins, intrinsically disordered proteins, and multidomain proteins, and predicts condensate saturation-concentration trends for A1-LCD variant systems. Analysis of the learned parameters reveals chemically interpretable interaction patterns and density-dependent effects that explain the model’s improved transferability. These results demonstrate that combining a generalized coarse-grained energy function with data-driven optimization can produce a practical and interpretable force field for protein conformational and condensate simulations.

## Introduction

Biomolecular condensates have emerged as a fundamental mechanism for organizing cellular biochemistry.^1–5^ These dynamic, membraneless assemblies regulate diverse processes, including transcription, signaling, and genome organization. Their ability to concentrate specific biomolecules within distinct chemical environments enables precise spatial and temporal control of cellular reactions.^1,4,6,7^ Dysregulation of condensate formation has been linked to neurodegenerative diseases and cancer, motivating intensive efforts to understand how amino acid sequences encode condensate stability, internal organization, and material properties, often described as the molecular grammar of phase separation.^4,8–39^ Decoding this molecular grammar represents a central challenge in connecting sequence-level interactions to emergent mesoscale behaviors.

Molecular simulations offer a powerful means to address this challenge.^11,22,29,40–46^ They provide atomistic to mesoscopic resolution of biomolecular assemblies and enable controlled perturbations of sequence and interaction strength to reveal the determinants of condensate formation. Coarse-grained (CG) models, which reduce the system representation while retaining residue-level chemical specificity, are particularly well suited for exploring condensate organization and thermodynamics at biologically relevant scales.^11,23,25–28,34,39,42,43,47–54^ Among these, one-bead-per-amino-acid (1BPA) models have yielded key mechanistic insights into molecular interactions that drive phase separation.^23–27,47,48,50,51^

CG models, however, often compromise on accuracy. They typically adopt pairwise additive interactions, analogous to those used in explicit-solvent force fields, to describe residueresidue interactions. Different parameterization strategies have been introduced, ranging from empirical mixing rules based on hydrophobicity scales to potentials of mean force derived from atomistic simulations or optimization against experimental observables.^55–62^ Despite these efforts, it is well established that coarse-graining introduces many-body effects^55,63–65^ that cannot be faithfully represented by pairwise interactions alone.

The compromise in accuracy of CG models is reflected in their lack of transferability. Several studies have shown that while 1BPA models can reproduce the dimensions of intrinsically disordered proteins (IDPs), often quantified by the radius of gyration (*R_g_*), they generally fail to stabilize folded proteins or to capture realistic conformations of multidomain systems.^22,30,56,62,66–68^ Although folded domains are frequently constrained in simulations, an accurate description of their underlying energetics remains essential for maintaining the correct thermodynamic balance between folded, disordered, and associated states. Misrepresenting these interactions can distort the coupling between ordered and disordered regions, alter interdomain packing, and bias condensate thermodynamics. To address such deficiencies, some models employ different representations or interaction potentials for folded and disordered domains.^51,69^ Although these artificial treatments can improve specific cases, they lack rigorous physical justification and may reduce model generality when applied to new systems. These limitations underscore the need for a physically grounded, transferable framework capable of describing folded, disordered, and hybrid proteins within a single potential.

Here we present **MOFF2**, a general protein CG force field designed to achieve consistent accuracy across folded, disordered, and multidomain systems. To address the limitations of pairwise additive models, MOFF2 incorporates a density-dependent many-body potential inspired by the AWSEM force field^55^ that captures local packing and solvent-mediated effects arising from coarse-graining. In parallel, we developed a data-driven optimization framework that enables efficient parameterization of complex energy functions using large and heterogeneous training datasets. The force field is trained against conformational ensembles and experimental observables spanning ordered proteins (OPs), IDPs, and multidomain proteins (MDPs), thereby ensuring balanced performance across diverse structural contexts.

We demonstrate that MOFF2 reproduces single-chain conformational statistics across protein classes with high fidelity. The model also quantitatively captures experimental trends in condensate saturation concentrations (*C*_sat_). By unifying the treatment of folded and disordered proteins within a single physically grounded framework, MOFF2 provides atransferable platform for simulating protein conformations and phase behavior.

## Results

### MOFF2: Model Overview

The limited transferability of existing 1BPA models remains a major obstacle to predictive simulations of biomolecular condensates. This limitation arises in part from the simplified interaction potentials commonly employed in these models, which often rely on pairwise additive interactions and therefore provide only a limited description of the physical factors governing protein association and conformational stability.^11,23,25–28,42,43,47–53,70^

To address these limitations, we developed MOFF2, a next-generation CG force field that extends conventional 1BPA models through a more expressive energy function. In addition to residue-specific pair interactions and screened electrostatics, MOFF2 incorporates both first-solvation-shell corrections and density-dependent many-body interactions (Figure 1A). Together, these terms enable the model to account for hydration effects, desolvation barriers, and environment-dependent packing contributions that are difficult to represent within conventional pairwise formulations.

**Figure 1:**
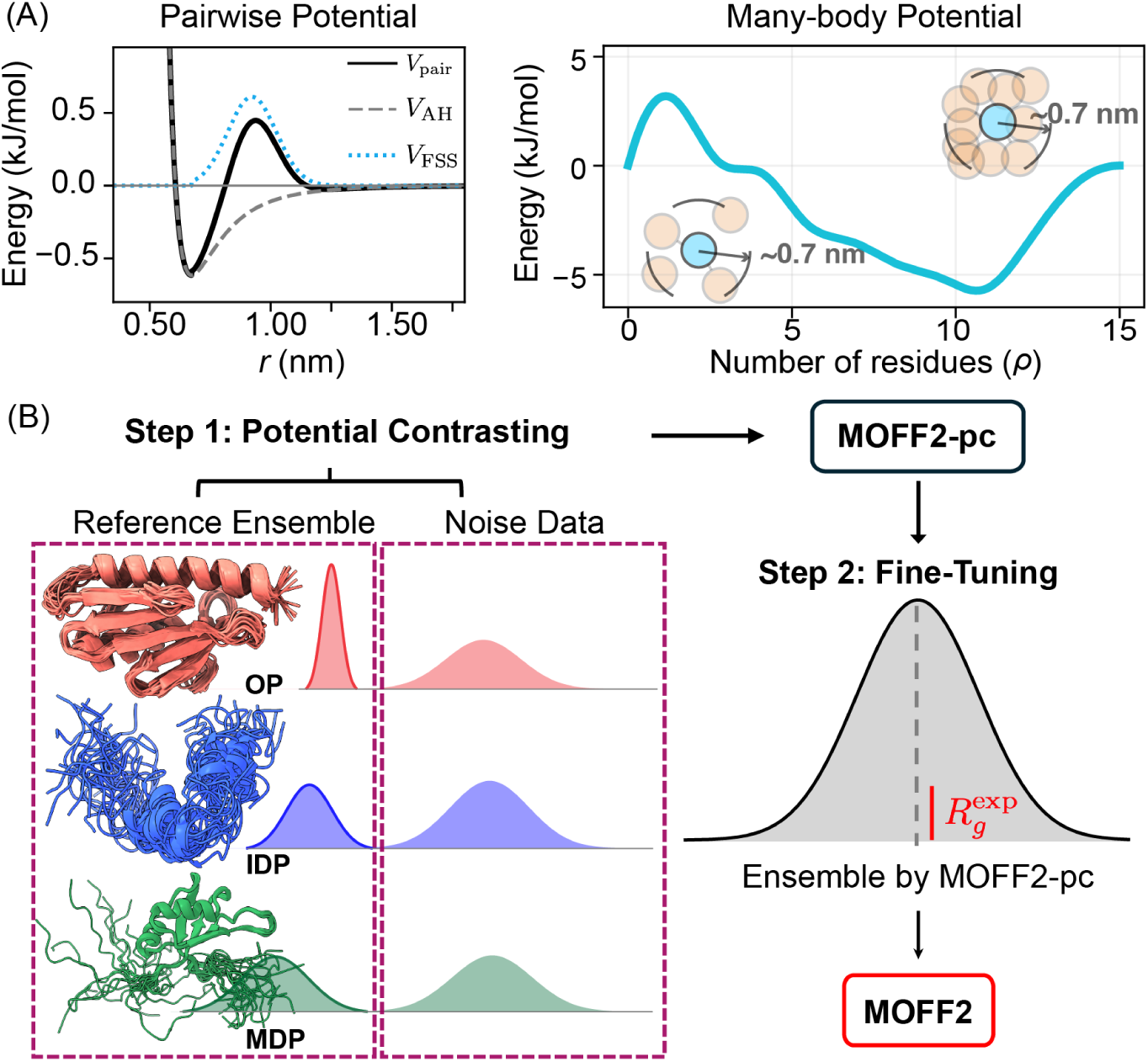
MOFF2 energy function and optimization workflow. (A) MOFF2 extends conventional 1BPA models through a flexible energy function that includes bonded and electrostatic interactions together with three nonbonded components: residue-specific pair interactions based on the Ashbaugh-Hatch (AH) potential (*V*_AH_),^71^ a Gaussian first-solvation-shell correction (*V*_FSS_) that accounts for desolvation barriers during contact formation, and a density-dependent many-body term that captures environment-dependent interactions. (B) Two-stage optimization framework combining bottom-up learning from conformational ensembles and top-down refinement against experimental measurements.

The pairwise interaction term in MOFF2 includes both direct amino-acid contacts^71^ and an explicit first-solvation-shell correction that captures the energetic barrier associated with displacing solvent molecules during contact formation. As a result, residue interactions are determined not only by their contact stability but also by the free-energy cost of removing intervening solvent. This additional level of physical realism improves the description of protein cores and intermolecular interfaces, where hydration effects play a critical role.^66,72,73^ To further account for collective effects arising from coarse-graining, MOFF2 includes a density-dependent many-body potential. Unlike pairwise interactions, the many-body term introduces an explicit dependence on the local packing density surrounding each amino acid. This allows the model to distinguish between physical environments that would otherwise appear identical in a pairwise description. For example, amino acids buried within a densely packed protein core experience different effective interactions than those in expanded or dilute conformations.^65,74–78^ By accounting for these differences, the many-body potential captures important effects such as hydrophobic burial, solvent-mediated stabilization, and local packing cooperativity, thereby enabling a unified description of OPs, IDPs, and MDPs.

### Transferability Across Protein Classes

The richer functional form of MOFF2 introduces a larger number of parameters and therefore requires a systematic optimization strategy. To address this challenge, we employ a two-stage framework that combines complementary bottom-up and top-down approaches (Figure 1B). First, the force field is trained against diverse conformational ensembles spanning OPs, IDPs, and MDPs. This stage uses potential contrasting optimization^79,80^ to reproduce the structural distributions of the reference ensembles. Second, the resulting model is refined against experimental observables using a thermodynamic reweighting procedure.^81,82^ Together, these steps enable efficient parameterization of a transferable force field while integrating information from both high-resolution simulations and experiments.

For the potential contrasting stage, the training set included 37 OPs, 41 IDPs, and 13 MDPs (Tables S1–S5), covering a broad range of chain lengths, amino-acid compositions, and structural organizations (Figure S1). The resulting model, MOFF2-pc, successfully reproduced the overall conformational trends observed across the training proteins. Strong correlations were obtained between predicted and reference ensemble properties (Figure 2A), indicating that the force field captures physical interactions that generalize across distinct structural classes.

**Figure 2:**
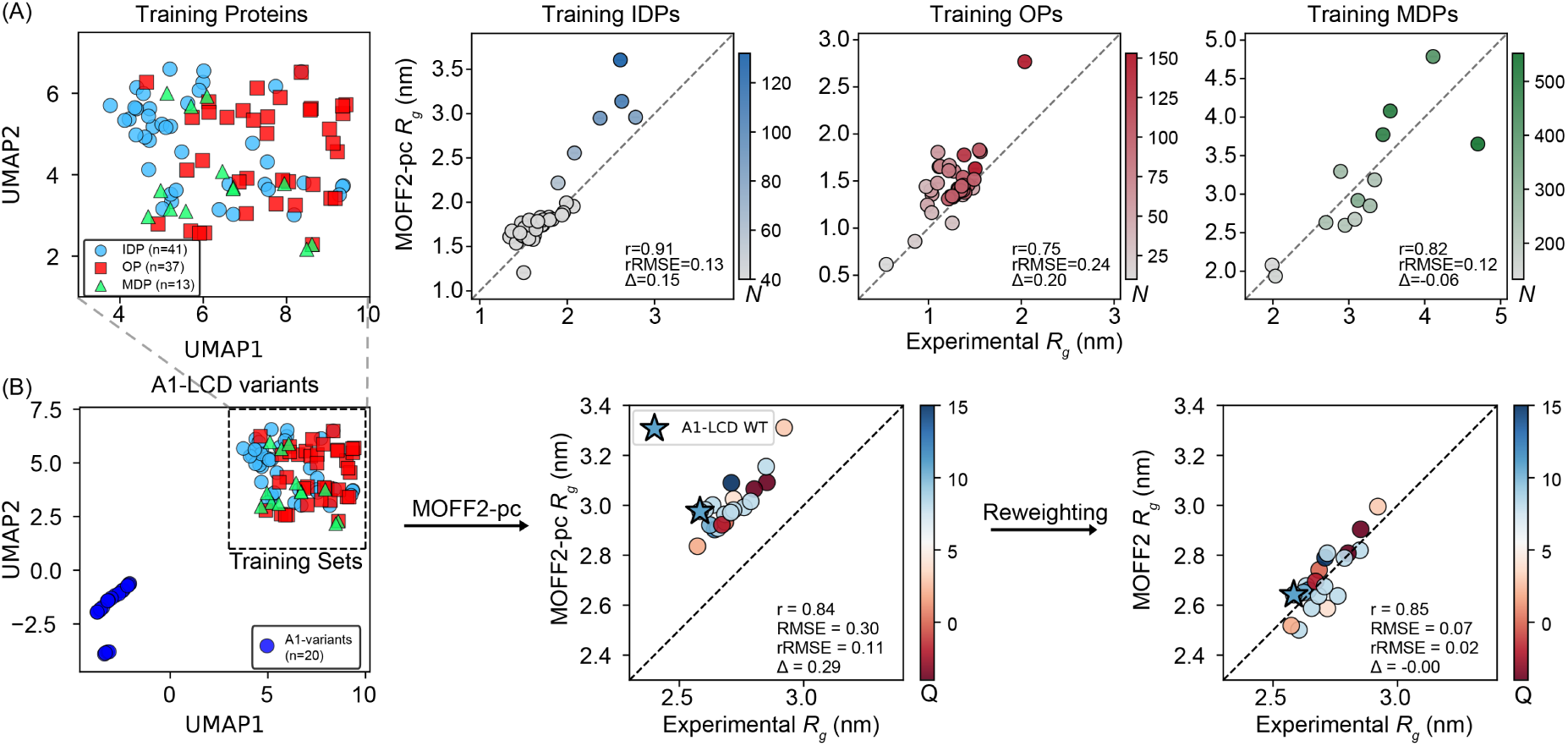
Two-stage optimization of MOFF2 establishes transferability across protein classes and sequence space. (A) Results of potential-contrasting optimization. The left panel shows a two-dimensional UMAP^83^ projection of sequence-composition features for proteins used for model optimization. The right panels compare reference and MOFF2pc predicted ⟨*R_g_*⟩ values for proteins in the training set. Proteins are colored by chain length, and dashed lines indicate perfect alignment. Legends report the Pearson correlation coefficient (*r*), the relative root-mean-square error (rRMSE), and the mean signed deviation (Δ) (see *Methods* section for detailed definitions). (B) Top-down refinement of sequence regions that are underrepresented during potential contrasting. The left panel shows the location of A1-LCD variants in the same sequence-feature space, highlighting their limited overlap with the potential contrasting training set. The middle and right panels compare simulation results from MOFF2-pc and MOFF2 with experimental values.

Importantly, the various components of the MOFF2 energy function contributed synergistically to this performance. Models containing only pairwise interactions exhibited less balanced behavior between OPs and MDPs, whereas inclusion of the density-dependent many-body term improved transferability across protein classes (Figure S2). Introducing the first-solvation-shell correction further enhanced predictions for MDPs while maintaining accuracy for OPs and IDPs. Together, these results demonstrate that chemically specific pair interactions, local solvation effects, and many-body packing contributions are all required to achieve balanced performance.

To evaluate generalization beyond the training data, we next tested MOFF2-pc on an independent validation set consisting of 9 IDPs (Table S6) and 14 MDPs (Table S8) that were not included during optimization. The model retained strong predictive power for these proteins (Figure S3), indicating that the learned interactions provide a broadly transferable physical description. Nevertheless, a systematic deviation emerged for the A1-LCD variant family, where the model consistently overestimated chain expansion (Figure 2B). This behavior is consistent with the sequence-space distribution of the training set. A1-LCD variants are enriched in Gly and Ser residues and occupy a region of sequence space that is sparsely represented among the proteins used for potential contrasting (Figure 2B and Figure S1). These results suggest that while potential contrasting establishes a robust global model, additional refinement may be beneficial for sequence families that are underrepresented in the bottom-up training ensembles.

Rather than generating new atomistic reference ensembles for these proteins, which would be computationally demanding and may still suffer from incomplete convergence, we used MOFF2-pc as a physically meaningful starting point and refined it against experimental observables using a thermodynamic reweighting method.^81,82^

For this refinement stage, we incorporated experimental measurements (⟨*R_g_*⟩) for 20 A1-LCD variants together with 14 MDPs and 21 OPs (Tables S7–S9). Including multiple protein classes prevents overfitting toward a single disordered protein family while preserving the global balance established during potential contrasting. The refinement substantially improved predictions for the A1-LCD variants and corrected the systematic overexpansion observed in the MOFF2-pc model (Figure 2B). At the same time, the accuracy for OPs and MDPs was maintained (Figure S4), indicating that the update improved local performance without compromising overall transferability.

We refer to the resulting two-stage optimized force field as MOFF2. To further assess transferability, we evaluated MOFF2 on an independent test set of proteins that were not included in either the potential contrasting optimization or the reweighting refinement (Figure 3, Tables S10–S12). For IDPs, MOFF2 achieved strong agreement with experimental ⟨*R_g_*⟩ values. For OPs and MDPs, where comprehensive experimental conformational ensembles are generally unavailable, we used AlphaFold-predicted structures^84^ as structural references. These references are expected to be more reliable for OPs than for MDPs because interdomain orientations and linker-mediated conformations are often predicted with lower confidence. Consequently, the larger deviations observed for some MDPs likely reflect uncertainties in the reference structures rather than deficiencies of the force field itself. Despite these challenges, MOFF2 preserved the overall conformational trends across all protein classes with strong correlations, supporting its transferability across OPs, IDPs, and MDPs.

**Figure 3:**
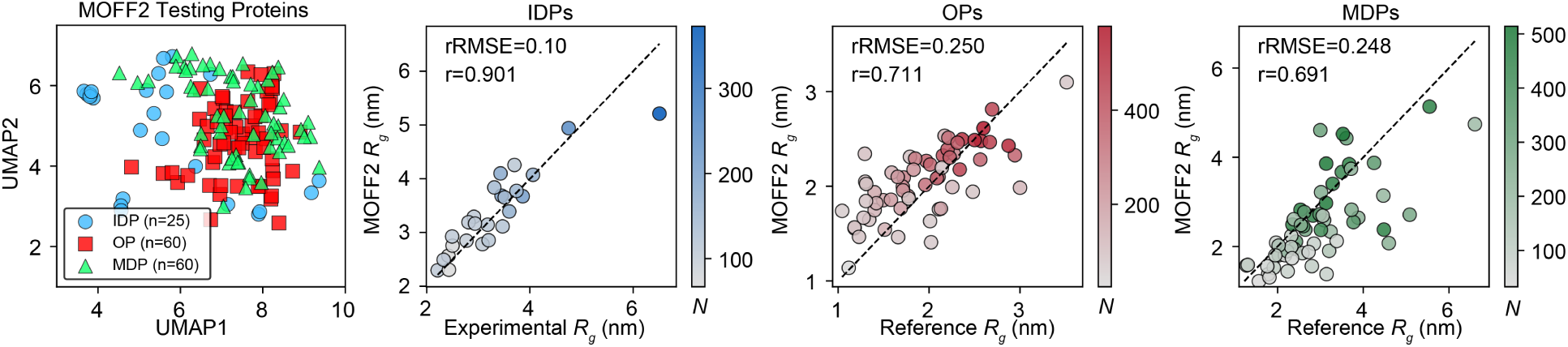
Evaluating MOFF2 on proteins unseen during training. The same plotting style as in Figure 2A is adopted here.

### MOFF2 accurately predicts phase behavior of A1-LCD variants

Having established the accuracy of MOFF2 for single-chain properties, we next evaluated its ability to predict sequence-dependent liquid–liquid phase separation. A1-LCD variants provide a particularly stringent benchmark because extensive experimental measurements^31,85–87^ are available for a diverse set of mutations that alter charge, aromatic composition, and sequence patterning, resulting in saturation concentrations that span a broad dynamic range. As shown in Figure 4A, MOFF2 reproduced the experimental saturation-concentration trends for most A1-LCD variants.^86,87^ In particular, MOFF2 captured the effect of the nuclear-localization-signal (NLS) region disruption, distinguishing weaker phase separation ability for A1-LCD^-NLS^ relative to wild-type A1-LCD^+NLS^ (P105G/Y106S mutants). The predicted phase diagrams showed good agreement with in vitro measurements for variants spanning broad classes of sequence perturbations, including aromatic substitutions (A1- LCD^+7F-7Y^, A1-LCD^-12F+12Y^, A1-LCD^-8F+4Y^, and A1-LCD^-9F+6Y^), charge-pattern variants (A1-LCD^-3R+3K^), and acidic-residue mutations (A1-LCD^+12E^).

**Figure 4:**
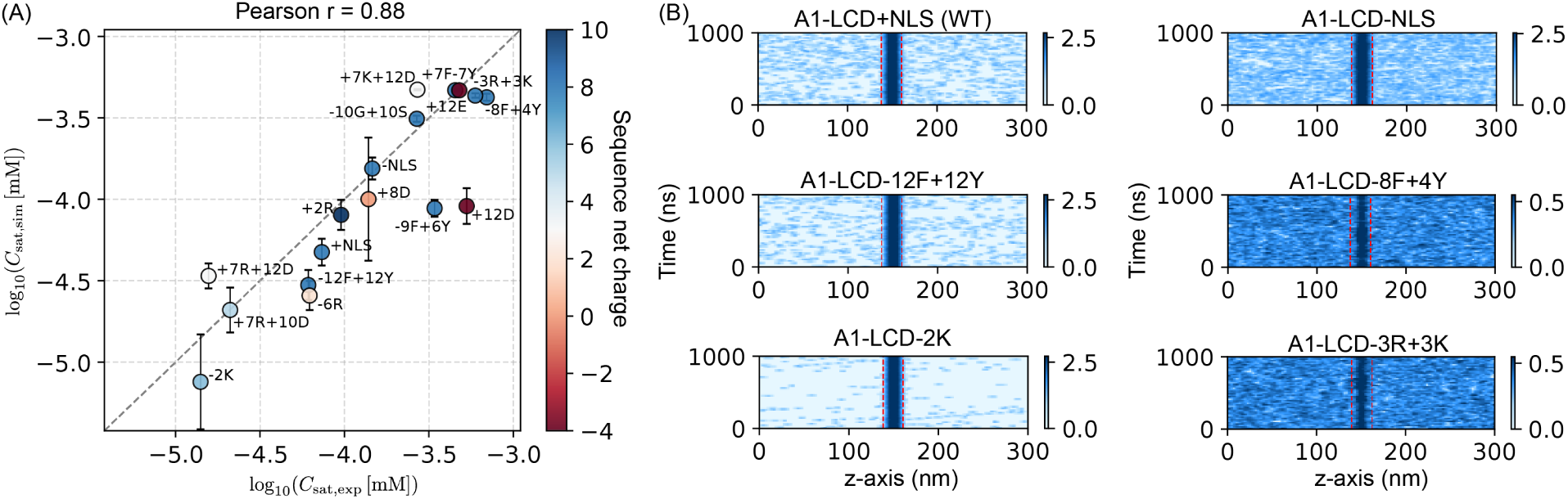
MOFF2 predicts sequence-dependent saturation concentrations of A1LCD variants. (A) Comparison between simulated and experimental saturation concentrations, *C*_sat_, for A1-LCD variants. Both axes show log_10_(*C*_sat_) in mM. Proteins are colored by sequence net charge, and error bars indicate the standard error estimated from five-block averaging of the simulated *C*_sat_. (B) Representative slab-density profiles used to estimate *C*_sat_ from coexistence simulations (see *Methods* for details). Each heat map shows the protein concentration profile along the *z*-axis during the final 1 *µ*s of the trajectory that was used for analysis. Red dashed lines mark the separation between condensed and dilute phase used to estimate the *C*_sat_.

MOFF2 performed particularly well for several challenging charge-engineered variants. For example, the saturation concentrations of A1-LCD^+7R+12D^ and A1-LCD^-6R^ were predicted in close agreement with experiment. Likewise, MOFF2 substantially improved the description of highly charged variants, including A1-LCD^+8D^, A1-LCD^+12E^, and A1-LCD^+12D^, for which Mpipi-Recharged, among the most accurate 1BPA models for LLPS prediction, still underestimates the saturation concentration by approximately two orders of magnitude.^88^

The largest discrepancies were observed for Trp-containing variants, including A1-LCD^allW^, A1-LCD^FtoW^, A1-LCD^YtoW^, and A1-LCD^W^(Table S13). This limitation is consistent with the relatively sparse representation of Trp-rich IDP sequence environments in the current training datasets (Figure S1). Consequently, these variants highlight a specific region of sequence space where the present parameterization remains weakly constrained, rather than indicating a general failure to predict A1-LCD phase behavior.

To place these results in context, we compared MOFF2 directly with Mpipi-Recharged on the same A1-LCD benchmark set (Table S13). The overall deviation was calculated as the mean absolute difference between simulated and experimental *C*_sat_ values in units of M, 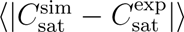. Despite the larger errors associated with Trp-containing variants, MOFF2 achieved an overall deviation of 0.048, comparable to the value of 0.044 reported for Mpipi-Recharged.^88^ These results demonstrate that MOFF2 achieves state-of-the-art accuracy for predicting sequence-dependent phase separation while retaining the broader transferability afforded by its physics-based parameterization.

### Learned pairwise and many-body interactions explain MOFF2 transferability

Finally, we analyzed the learned interaction parameters to understand how MOFF2 achieves balanced performance across protein classes and condensate-forming systems. Because MOFF2 decomposes residue interactions into direct pairwise contacts, first-solvation-shell corrections, and density-dependent many-body interactions, the model provides an opportunity to examine how distinct physical contributions emerge during optimization. We therefore asked whether these learned terms recover chemically interpretable features of protein biophysics and how they contribute to transferability.

We first examined the learned pairwise interaction strength *λ*_AH_, which describes the stability of direct residue–residue contacts. As shown in Figure 5A and Figures S5–S6, the learned interaction matrix captures several well-established sequence determinants of protein folding and phase separation. Most notably, MOFF2 naturally learns favorable cation–*π* interactions involving aromatic residues and positively charged side chains. Arg– aromatic contacts are substantially more favorable than the corresponding Lys–aromatic interactions, with Arg–Tyr among the strongest interactions in the matrix. This distinction is particularly important for condensate-forming proteins, where Arg-to-Lys substitutions frequently weaken phase separation despite preserving net charge.^11,47,86,89,90^ MOFF2 also favors aromatic–aromatic interactions, including Tyr-containing contacts that have been repeatedly implicated in driving condensation of low-complexity domains.^11,31,86,90–93^ Importantly, these chemically specific interaction patterns emerge directly from the optimization procedure and were not imposed through explicit interaction rules.

**Figure 5:**
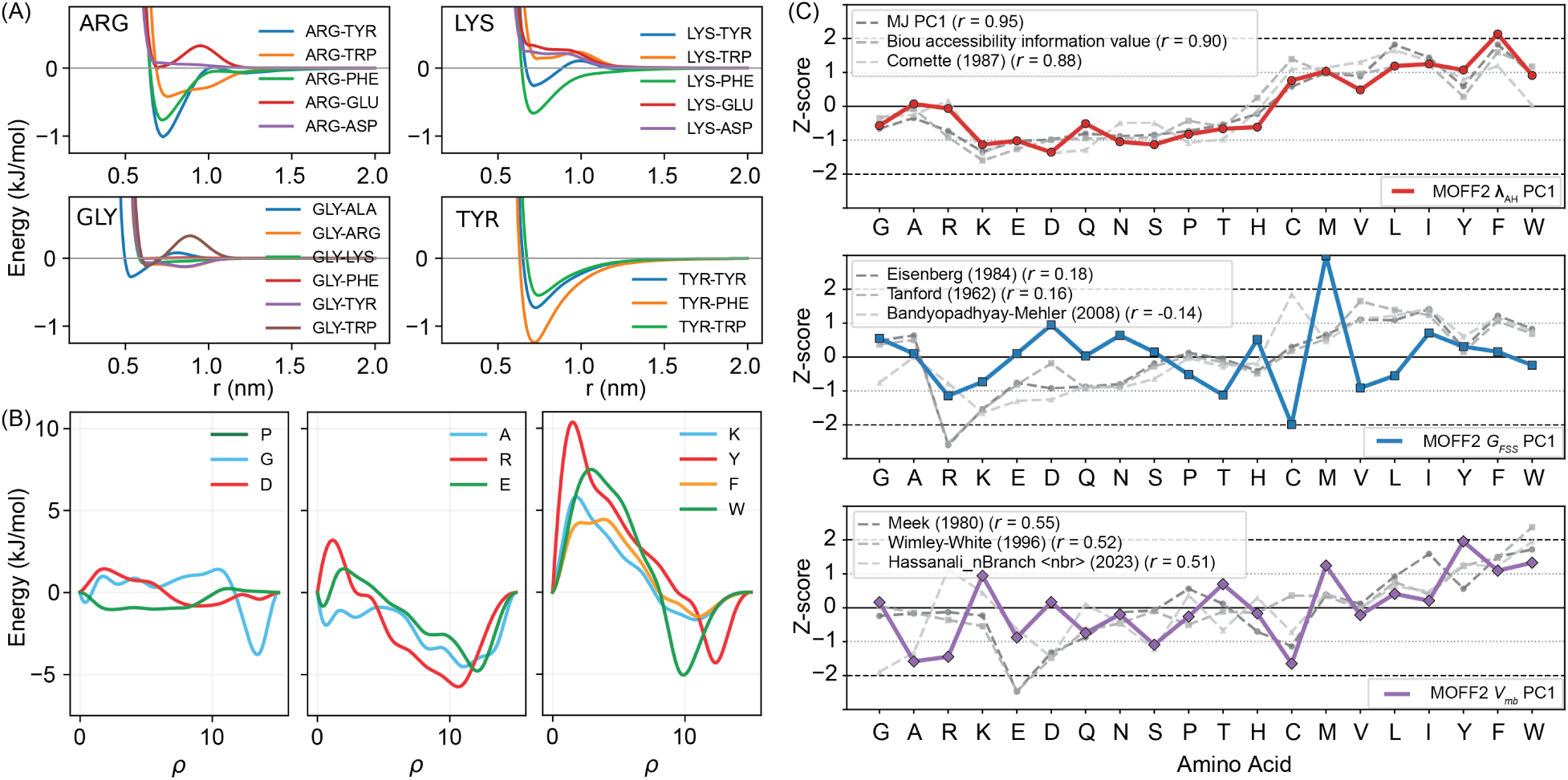
Interpretable chemical information from MOFF2 learned profiles. (A) Representative residue-pair interaction potentials learned by MOFF2. (B) Representative amino-acid-specific density-dependent many-body profiles. (C) Comparison between the first principal components (PC1) of the learned MOFF2 parameter sets: *λ*_AH_ (top), *G*_FSS_ (middle), and *V*_mb_ (bottom), and representative HPS references. Colored lines show the standardized MOFF2 PC1 profiles, gray lines show the three most correlated reference scales, and Pearson correlation coefficients (*r*) are reported in the legends.

The first-solvation-shell correction *G*_FSS_ provides a complementary description of residue interactions. Whereas the pairwise term quantifies how favorable a contact is once formed, the solvation-shell term captures the energetic cost of transitioning between solvent-separated and direct-contact states. Several physically meaningful patterns emerge from the learned barriers. For example, Arg–Glu exhibits a pronounced barrier, reflecting the energetic cost of removing hydration waters before a salt bridge can form. ^94,95^ Interestingly, Gly–Trp also displays a substantial first-solvation-shell correction despite relatively weak direct-contact attraction. This feature may reflect the unique CH–*π* interaction between the glycine backbone and the Trp aromatic ring, which has recently been implicated in regulating phase separation of low-complexity domains.^42,96–98^

The density-dependent many-body interactions provide a third and distinct layer of physical information. Rather than describing specific pairwise contacts, these terms encode how amino-acid preferences change with local packing density. As shown in Figure 5B and Figure S6C, aromatic and hydrophobic residues such as Trp, Tyr, and Phe become increasingly favorable in densely packed environments, consistent with burial-driven stabilization and the hydrophobic effect. In contrast, many polar and charged residues display weaker density dependence and remain comparatively favorable in dilute or partially packed environments.^99–101^ The distinction between Arg and Lys is particularly notable. Arg exhibits favorable contributions across a broader range of densities, whereas Lys shows a stronger preference for highly packed environments. These observations suggest that the many-body term captures environmental preferences that cannot be represented through pairwise interactions alone and likely contributes to the transferability of MOFF2 across proteins spanning heterogeneous conformational space and interaction environments.

To place the learned interactions in the context of existing amino-acid descriptors, we compared the principal components of each MOFF2 interaction class (Figure S7) with established hydrophobicity and residue-interaction scales (Figures 5C and Figure S8). The direct pairwise contact term correlates strongly with the Miyazawa–Jernigan (MJ)^102^ potential. This result is not unexpected because the pairwise contact parameters were initialized from the MJ matrix. Nevertheless, the high correlation indicates that the optimization procedure largely preserves the residue-contact preferences encoded in MJ while refining other components of the force field.

In contrast, the density-dependent many-body term shows strong correlations with interfacial hydrophobicity scales such as Wimley–White^103^ (Pearson *r* = 0.52). This result is particularly notable because no such information was imposed during training, indicating that MOFF2 independently learned environmental preferences associated with the hydrophobic effect. The first-solvation-shell correction exhibits weaker correlations with existing scales, likely reflecting both its smaller magnitude and the absence of closely related descriptors in the chosen literature. Nevertheless, its strongest correspondence is with the Eisenberg hydrophobicity scale,^104^ which measures residue interactions with water and is therefore conceptually aligned with the hydration-mediated processes captured by this term.

Together, the emergence of these chemically interpretable features provides a mechanistic explanation for the ability of MOFF2 to accurately describe various systems within a unified framework.

## Conclusions and Discussion

Here, we present MOFF2, a protein CG force field designed to capture residue-specific interactions, local solvation effects, and environment-dependent packing contributions within a unified framework. By coupling this physically motivated energy function with a two-stage optimization strategy that integrates potential contrasting and thermodynamic reweighting, MOFF2 establishes a transferable framework for protein modeling. The resulting model accurately reproduces conformational properties across OPs, IDPs, and MDPs while capturing sequence-dependent trends in biomolecular phase separation.

Despite these advances, the present study also highlights important opportunities for further improvement. Although the training set spans a broad range of proteins, the transferability of MOFF2 remains fundamentally constrained by the diversity of the available training data. Certain sequence environments, particularly aromatic-rich and low-complexity sequences, remain underrepresented. As illustrated by the residual deviations observed for several A1-LCD variants, interactions in these regions of sequence space are less tightly constrained than those represented more extensively in the training set. More generally, the current optimization strategy relies primarily on conformational ensembles and single-chain observables, whereas condensate thermodynamics are only indirectly incorporated through the underlying molecular interactions.

A promising direction for future development is therefore to extend the refinement framework directly to condensed-phase properties. The thermodynamic reweighting strategy introduced here is not limited to single-chain simulations and could, in principle, be generalized to condensate ensembles. Experimental observables such as saturation concentrations could then be incorporated directly into force-field optimization. Such an approach would enable systematic refinement of intermolecular interactions while preserving the physically meaningful baseline established through bottom-up learning.

More broadly, the combination of many-body interactions, atomistically informed training data, and statistically controlled optimization provides a general framework for devel- oping next-generation CG force fields. These ideas should be readily extendable to nucleic acids, protein–nucleic acid assemblies, and other multicomponent biomolecular systems in which environment-dependent interactions play a central role. We anticipate that MOFF2 will serve both as a practical platform for studying biomolecular condensates and as a foundation for future efforts toward predictive CG modeling of complex cellular assemblies.

## Methods

### MOFF2 energy function

MOFF2 employs a 1BPA representation in which each amino acid is mapped to its C*α* atom. The resulting coarse-grained configuration is described by the energy function

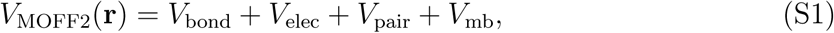

where *V*_bond_ maintains chain connectivity, *V*_elec_ describes screened electrostatic interactions, *V*_pair_ describes residue-specific pair interactions, and *V*_mb_ captures density-dependent manybody effects. Detailed mathematical forms are provided in the *Supporting Information: Explicit Energy Function*.

The pairwise nonbonded interaction accounts for both the direct amino acid contacts, described by the Ashbaugh–Hatch (AH) potential,^105^ and the desolvation barrier, described by a Gaussian potential, that amino acids must overcome as they approach one another to form contacts.

To capture collective effects that are not representable by pairwise additive interactions, MOFF2 includes a density-dependent many-body potential,

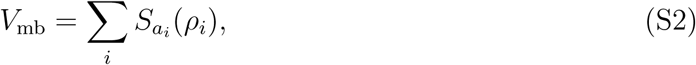

where *a_i_* is the amino-acid type, *ρ_i_*is the local residue density, and *S_ai_* is an amino-acid-specific cubic B-spline function.^106^

For proteins containing folded domains, an additional structure-based potential, *V*_native_, is used to maintain the reference fold. This term includes angle, dihedral, and native-contact restraints. During training and model evaluation of folded proteins, native contacts are restricted to contacts within the same continuous secondary-structure element, so long-range tertiary compaction is governed primarily by the transferable MOFF2 nonbonded terms.

### Model optimization

We employ a two-stage parameterization framework that integrates complementary bottomup and top-down approaches. A brief overview is provided below, while detailed descriptions of the training data and optimization procedures can be found in the *Supporting Information: Force Field Optimization*.

#### Potential contrasting optimization

In the first stage, we parameterized MOFF2, using potential contrasting,^79,80^ a variational framework that learns a coarse-grained energy function from reference configurational ensembles. Potential contrasting compares configurations sampled from the target data distribution with configurations sampled from a broader noise distribution. The optimized model assigns higher Boltzmann probability to data configurations than to noise configurations, thereby learning an energy landscape consistent with the reference ensemble without requiring force matching or repeated molecular dynamics simulations during optimization.

The optimized parameters include the 210 AH pair coefficients, the 210 Gaussian potential corrections, and the 240 coefficients of the density-dependent spline potential. Regularization terms were applied separately to the pairwise and density-dependent parameter classes to control parameter magnitude and improve transferability.

The training data include OPs, IDPs, and MDPs. For OPs and IDPs, configurations were collected from explicit-solvent atomistic molecular dynamics simulations. The OP training set includes long-timescale trajectories of 19 proteins reported by D. E. Shaw Research. ^107–109^ To increase sequence and structural diversity, we additionally generated trajectories for 18 ordered proteins using the a99SB-*disp* force field in OpenMM.^110^ The IDP training set includes seven trajectories from D. E. Shaw Research^108^ and 34 additional trajectories from Ref. 13.

For MDPs, converged explicit-solvent atomistic simulations are not practical because of their larger system sizes and slow interdomain rearrangements. We therefore generated coarse-grained conformational ensembles and reweighted them by a maximum-entropy procedure so that the reweighted ensembles reproduce experimental mean radii of gyration, ⟨*R_g_*⟩.

Noise ensembles were generated using HPS-Urry-based coarse-grained simulations.^111^ For IDPs, umbrella sampling was performed along *R_g_*. For OPs, the HPS-Urry potential was supplemented with *V*_native_, and umbrella sampling was performed along the root-mean-square deviation (RMSD) from a representative folded reference structure. Configurations from all umbrella windows were combined into a generalized ensemble using MBAR.^112^

#### Fine-tuning with thermodynamic reweighting

In the second stage, we further refine the force field against experimental measurements for a selected set of larger proteins (Table S7–S9), where constructing fully converged atomistic ensembles remains challenging. This step follows a top-down strategy, in which model parameters are adjusted to reproduce experimentally observed ensemble-averaged properties. Here, we use the *R_g_* as the target observable. The optimization is carried out using a thermodynamic reweighting framework,^81^ combined with regularization based on effective sample size^82^ to ensure stable and efficient parameter updates without additional simulations. This scheme allows estimation of how small perturbations in the energy function parameters affect an observable, i.e., *R_g_*, using reference ensembles that only need to be constructed once.

### Molecular dynamics simulation details

MOFF2 was implemented in OpenABC,^36^ a Python-based simulation package built on OpenMM for GPU-accelerated molecular dynamics simulations of coarse-grained biomolecular systems.

#### Single-chain simulations

Simulations of individual proteins were performed in the NVT ensemble using the Langevin middle integrator,^113^ with a friction coefficient of 1 ps*^−^*^1^ and a timestep of 10 fs. Each system was energy-minimized and equilibrated for 5 × 10^5^ steps before production. Production simulations were run for 2 × 10^8^ steps, corresponding to 2 *µ*s, in a cubic box with a side length of 1000 nm. Coordinates were saved every 5 × 10^3^ steps, yielding 4 × 10^4^ frames per trajectory. Simulation temperatures and ionic strengths were matched to the corresponding experimental conditions when available, as summarized in Table S6–S10. For OP and MDP MOFF2 testing simulations without experimental conditions, simulations were performed at 298 K and 150 mM ionic strength, as summarized in Tables S11–S12.

#### Condensate simulations

For A1-LCD phase-separation simulations, the initial multi-chain configurations were generated by randomly inserting 100 copies of each A1-LCD variant into a rectangular simulation box of size 25 × 25 × 300 nm^3^. Systems were first compressed under NPT conditions using a Monte Carlo barostat^114^ at 1 bar and the Langevin middle integrator with a 10 fs timestep. The NPT compression was performed for 8.0 × 10^4^ steps, and the resulting configuration was used to initialize the slab simulations. After compression, the barostat was removed and the simulation box was reset to 25 × 25 × 300 nm^3^. Slab simulations were performed at 150 mM ionic strength and at the temperature matched for experimental values (Table S13). Production simulations were performed for 2.0 *µ*s and configurations were saved every 1×10^5^ steps. The trajectories were recentered and aligned along the slab normal prior to calculating concentration profiles and saturation concentrations.

The saturation concentration (*C*_sat_) was estimated from slab simulations using the density of the dilute phase. For each A1-LCD variant, the first 1 *µ*s of the trajectory was discarded as equilibration. A one-dimensional *z*-density profile was computed from the retained frames and smoothed with a moving-average window. The bin with the maximum smoothed density was assigned as the dense-phase center. A dense-core region was then defined using neighboring bins around this maximum, with an additional 5 nm margin on both sides (red dashed line in Figure 4B). Protein chains outside this dense-core region were used to estimate the dilute-phase concentration.

### Analysis details

#### UMAP sequence analysis

UMAP^83^ was used to visualize protein sequence-composition space across training, fine-tuning, and test sets. Each protein was represented by a 20dimensional amino-acid composition vector containing the fractional abundance of each residue type. UMAP was then applied to the combined composition matrix using Euclidean distance, with *n*_neighbors_ = 20, min dist = 0.1.

#### PCA analysis of MOFF2 interactions

To interpret the residue-level chemical information encoded by MOFF2, we analyzed the learned interaction components using principal component analysis (PCA).^115,116^ PCA was applied separately to each MOFF2 component, including the AH interaction matrix *λ*_AH_, the first-solvation-shell Gaussian matrix *G*_FSS_, and the density-dependent many-body potential *V*_mb_ (Figures S6–S7).

For matrix-valued components, including *λ*_AH_, *G*_FSS_, each amino acid was represented by its interaction profile against all 20 amino acids, yielding a 20-dimensional vector per amino acid. For the density-dependent many-body term, each amino acid is represented by the energy values on a uniform grid of 500 local-density values between *ρ*_min_ = 0 and *ρ*_max_ = 15. Before PCA, each profile matrix was standardized feature-wise using *z*-score normalization. The resulting PC scores provide compact amino-acid-level representations of the dominant chemical trends encoded by each learned component. In particular, PC1 was used as a one-dimensional residue profile for comparison with published hydrophobicity, solvation, and residue-interaction scales.

For reference comparisons, we included the Miyazawa–Jernigan model,^102^ the HPS–Urry scale,^25^ and 21 additional published residue-level scales.^100,103,104,117–131^ Reference scales, except for Miyazawa–Jernigan model, were provided as one-dimensional amino-acid profiles and standardized using *z*-score normalization for comparisons with the MOFF2 PC1 profiles. The Miyazawa–Jernigan model was first analyzed using the same PCA procedure applied to the MOFF2 matrix-valued components. Its resulting PC1 amino-acid profiles were then used for comparison.

#### Relative root-mean-square error (rRMSE)

rRMSE was used to quantify the prediction error relative to the reference value for each protein,

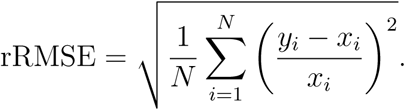

Here, *x_i_* and *y_i_* are the reference and predicted values for protein *i*, respectively, and *N* is the number of proteins in the corresponding evaluation set.

**Mean signed deviation (**Δ). Δ was used to quantify systematic bias,

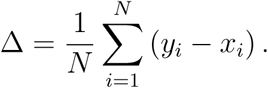

Here, *x_i_* and *y_i_* are the reference and predicted values for protein *i*, respectively, and *N* is the number of proteins in the corresponding evaluation set. Positive Δ indicates that the model overestimates the reference values on average, whereas negative Δ indicates underestimation.

## Acknowledgement

This work was supported by the National Institutes of Health (Grant R35GM133580) and the National Science Foundation (MCB-2042362). This work used Bridges-2 at PSC through allocation BIO240299 from the Advanced Cyberinfrastructure Coordination Ecosystem: Services & Support (ACCESS) program, which is supported by U.S. National Science Foundation grants #2138259, #2138286, #2138307, #2137603, and #2138296.

## Competing interests

The authors declare that they have no competing interests.

## Data and materials availability

MOFF2 is implemented in the open-source software package OpenABC. Workflow for model development is available on GitHub: MOFF2- development. Tutorials for setting up simulations are available on GitHub: MOFF2-simulations. All data and results used for this work are available on GitHub: paper-data.

## Notes

### Competing Interest Statement

The authors have declared no competing interest.

## References

(1) Banani, S. F.; Lee, H. O.; Hyman, A. A.; Rosen, M. K. Biomolecular Condensates: Organizers of Cellular Biochemistry. Nature Reviews Molecular Cell Biology 2017, 18, 285–298.

(2) Hirose, T.; Ninomiya, K.; Nakagawa, S.; Yamazaki, T. A Guide to Membraneless Organelles and Their Various Roles in Gene Regulation. Nature Reviews Molecular Cell Biology 2023, 24, 288–304.

(3) Choi, J.-M.; Holehouse, A. S.; Pappu, R. V. Physical Principles Underlying the Complex Biology of Intracellular Phase Transitions. Annual Review of Biophysics 2020, 49, 107–133.

(4) Pappu, R. V.; Cohen, S. R.; Dar, F.; Farag, M.; Kar, M. Phase Transitions of Associative Biomacromolecules. Chemical Reviews 2023, 123, 8945–8987.

(5) Ginell, G. M.; Holehouse, A. S. In Phase-Separated Biomolecular Condensates: Methods and Protocols; Zhou, H.-X., Spille, J.-H., Banerjee, P. R., Eds.; Springer US: New York, NY, 2023; pp 95–116.

(6) Hyman, A. A.; Weber, C. A.; Jülicher, F. Liquid-Liquid Phase Separation in Biology. Annual Review of Cell and Developmental Biology 2014, 30, 39–58.

(7) Feric, M.; Vaidya, N.; Harmon, T. S.; Mitrea, D. M.; Zhu, L.; Richardson, T. M.; Kriwacki, R. W.; Pappu, R. V.; Brangwynne, C. P. Coexisting Liquid Phases Underlie Nucleolar Subcompartments. Cell 2016, 165, 1686–1697.

(8) Alberti, S.; Dormann, D. Liquid–Liquid Phase Separation in Disease. Annual Review of Genetics 2019, 53, 171–194.

(9) Zbinden, A.; Pérez-Berlanga, M.; De Rossi, P.; Polymenidou, M. Phase Separation and Neurodegenerative Diseases: A Disturbance in the Force. Developmental Cell 2020, 55, 45–68.

(10) Rai, S. K.; Savastano, A.; Singh, P.; Mukhopadhyay, S.; Zweckstetter, M. LiquidLiquid Phase Separation of Tau: From Molecular Biophysics to Physiology and Disease. Protein Science: A Publication of the Protein Society 2021, 30, 1294–1314.

(11) Das, S.; Lin, Y.-H.; Vernon, R. M.; Forman-Kay, J. D.; Chan, H. S. Comparative Roles of Charge, *π*, and Hydrophobic Interactions in Sequence-Dependent Phase Separation of Intrinsically Disordered Proteins. Proceedings of the National Academy of Sciences 2020, 117, 28795–28805.

(12) Zhang, Y.; Zheng, J.; Zhang, B. Protein Language Model Identifies Disordered, Conserved Motifs Implicated in Phase Separation. eLife 2025, 14 .

(13) Wang, C.; Zhang, B. Sequence-Dependent Conformational Landscapes of Intrinsically Disordered Proteins Reveal Asymmetric Chain Compaction. Journal of Chemical Theory and Computation 2025, 21, 11282–11292.

(14) Schuette, G.; Lao, Z.; Zhang, B. ChromoGen: Diffusion Model Predicts Single-Cell Chromatin Conformations. Science Advances 2025, 11, eadr8265.

(15) Brangwynne, C. P.; Tompa, P.; Pappu, R. V. Polymer Physics of Intracellular Phase Transitions. Nature Physics 2015, 11, 899–904.

(16) Zeng, X.; Pappu, R. V. Developments in Describing Equilibrium Phase Transitions of Multivalent Associative Macromolecules. Current Opinion in Structural Biology 2023, 79, 102540.

(17) Schmit, J. D.; Bouchard, J. J.; Martin, E. W.; Mittag, T. Protein Network Structure Enables Switching between Liquid and Gel States. Journal of the American Chemical Society 2020, 142, 874–883.

(18) Mittag, T.; Pappu, R. V. A Conceptual Framework for Understanding Phase Separation and Addressing Open Questions and Challenges. Molecular Cell 2022, 82, 2201–2214.

(19) Dzuricky, M.; Roberts, S.; Chilkoti, A. Convergence of Artificial Protein Polymers and Intrinsically Disordered Proteins. Biochemistry 2018, 57, 2405–2414.

(20) Flory, P. J. Thermodynamics of High Polymer Solutions. The Journal of Chemical Physics 1942, 10, 51–61.

(21) Huggins, M. L. Some Properties of Solutions of Long-chain Compounds. The Journal of Physical Chemistry 1942, 46, 151–158.

(22) Dignon, G. L.; Zheng, W.; Kim, Y. C.; Best, R. B.; Mittal, J. Sequence Determinants of Protein Phase Behavior from a Coarse-Grained Model. PLOS Computational Biology 2018, 14, e1005941.

(23) Dignon, G. L.; Zheng, W.; Mittal, J. Simulation Methods for Liquid–Liquid Phase Separation of Disordered Proteins. Current Opinion in Chemical Engineering 2019, 23, 92–98.

(24) Latham, A. P.; Zhang, B. Maximum Entropy Optimized Force Field for Intrinsically Disordered Proteins. Journal of Chemical Theory and Computation 2020, 16, 773– 781.

(25) Regy, R. M.; Thompson, J.; Kim, Y. C.; Mittal, J. Improved Coarse-Grained Model for Studying Sequence Dependent Phase Separation of Disordered Proteins. Protein Science 2021, 30, 1371–1379.

(26) Latham, A. P.; Zhang, B. Consistent Force Field Captures Homologue-Resolved HP1 Phase Separation. Journal of Chemical Theory and Computation 2021, 17, 3134–3144.

(27) Tesei, G.; Schulze, T. K.; Crehuet, R.; Lindorff-Larsen, K. Accurate Model of Liquid– Liquid Phase Behavior of Intrinsically Disordered Proteins from Optimization of Single-Chain Properties. Proceedings of the National Academy of Sciences 2021, 118, e2111696118.

(28) Benayad, Z.; von Bülow, S.; Stelzl, L. S.; Hummer, G. Simulation of FUS Protein Condensates with an Adapted Coarse-Grained Model. Journal of Chemical Theory and Computation 2021, 17, 525–537.

(29) Latham, A. P.; Zhang, B. Molecular Determinants for the Layering and Coarsening of Biological Condensates. Aggregate 2022, 3, e306.

(30) Latham, A. P.; Zhang, B. Unified Protein Force Field for Simulations of Liquid-Liquid Phase Separation. Biophysical Journal 2022, 121, 356a–357a.

(31) Farag, M.; Borcherds, W. M.; Bremer, A.; Mittag, T.; Pappu, R. V. Phase Separation of Protein Mixtures Is Driven by the Interplay of Homotypic and Heterotypic Interactions. Nature Communications 2023, 14, 5527.

(32) Lotthammer, J. M.; Ginell, G. M.; Griffith, D.; Emenecker, R. J.; Holehouse, A. S. Direct Prediction of Intrinsically Disordered Protein Conformational Properties from Sequence. Nature Methods 2024,

(33) von Bülow, S.; Tesei, G.; Lindorff-Larsen, K. Prediction of Phase Separation Propensities of Disordered Proteins from Sequence. https://www.biorxiv.org/content/10.1101/2024.06.03.597109v1, 2024.

(34) Zhang, Y.; Li, S.; Gong, X.; Chen, J. Toward Accurate Simulation of Coupling between Protein Secondary Structure and Phase Separation. Journal of the American Chemical Society 2024, 146, 342–357.

(35) Kapoor, U.; Kim, Y. C.; Mittal, J. Coarse-Grained Models to Study Protein–DNA Interactions and Liquid–Liquid Phase Separation. Journal of Chemical Theory and Computation 2024, 20, 1717–1731.

(36) Liu, S.; Wang, C.; Latham, A. P.; Ding, X.; Zhang, B. OpenABC Enables Flexible, Simplified, and Efficient GPU Accelerated Simulations of Biomolecular Condensates. PLOS Computational Biology 2023, 19, e1011442.

(37) Zhou, L.; Zhu, L.; Wang, C.; Xu, T.; Wang, J.; Zhang, B.; Zhang, X.; Wang, H. Multiphasic Condensates Formed with Mono-Component of Tetrapeptides via Phase Separation. Nature Communications 2025, 16, 2706.

(38) Liu, S.; Athreya, A.; Lao, Z.; Zhang, B. From nucleosomes to compartments: physicochemical interactions underlying chromatin organization. Annual review of biophysics 2024, 53 .

(39) Qiu, Y.; Liu, S.; Lin, X.; Unarta, I. C.; Huang, X.; Zhang, B. Nucleosome condensate and linker DNA alter chromatin folding pathways and rates. Biophysical Journal 2026, 125, 282–293.

(40) Dignon, G. L.; Zheng, W.; Best, R. B.; Kim, Y. C.; Mittal, J. Relation between Single-Molecule Properties and Phase Behavior of Intrinsically Disordered Proteins. Proceedings of the National Academy of Sciences 2018, 115, 9929–9934.

(41) Dignon, G. L.; Zheng, W.; Kim, Y. C.; Mittal, J. Temperature-Controlled Liquid– Liquid Phase Separation of Disordered Proteins. ACS Central Science 2019, 5, 821– 830.

(42) Vernon, R. M.; Chong, P. A.; Tsang, B.; Kim, T. H.; Bah, A.; Farber, P.; Lin, H.; Forman-Kay, J. D. Pi-Pi Contacts Are an Overlooked Protein Feature Relevant to Phase Separation. eLife 2018, 7, e31486.

(43) Chou, H.-Y.; Aksimentiev, A. Single-Protein Collapse Determines Phase Equilibria of a Biological Condensate. The Journal of Physical Chemistry Letters 2020, 11, 4923– 4929.

(44) Latham, A. P.; Zhang, B. On the Stability and Layered Organization of Protein-DNA Condensates. Biophysical Journal 2022, 121, 1727–1737.

(45) Latham, A. P.; Zhu, L.; Sharon, D. A.; Ye, S.; Willard, A. P.; Zhang, X.; Zhang, B. Microphase Separation Produces Interfacial Environment within Diblock Biomolecular Condensates. eLife 2024, 12 .

(46) Jiang, H.; Wang, S.; Huang, Y.; He, X.; Cui, H.; Zhu, X.; Zheng, Y. Phase Transition of Spindle-Associated Protein Regulate Spindle Apparatus Assembly. Cell 2015, 163, 108–122.

(47) Joseph, J. A.; Reinhardt, A.; Aguirre, A.; Chew, P. Y.; Russell, K. O.; Espinosa, J. R.; Garaizar, A.; Collepardo-Guevara, R. Physics-Driven Coarse-Grained Model for Biomolecular Phase Separation with near-Quantitative Accuracy. Nature Computational Science 2021, 1, 732–743.

(48) Dannenhoffer-Lafage, T.; Best, R. B. A Data-Driven Hydrophobicity Scale for Predicting Liquid–Liquid Phase Separation of Proteins. The Journal of Physical Chemistry B 2021, 125, 4046–4056.

(49) Maristany, M. J.; Emelianova, A.; Chew, P. Y.; Aguirre, A.; Collepardo-Guevara, R.; Joseph, J. A. Modeling Biomolecular Condensates across Scales: Atomistic, CoarseGrained, and Data-Driven Approaches. Advances in Physics: X 2025, 10, 2592547.

(50) Chakravarti, A.; Joseph, J. A. Accurate Prediction of Thermoresponsive Phase Behavior of Disordered Proteins. Protein Science 2025, 34, e70284.

(51) Cao, F.; von Bülow, S.; Tesei, G.; Lindorff-Larsen, K. A Coarse-Grained Model for Disordered and Multi-Domain Proteins. Protein Science 2024, 33, e5172.

(52) Latham, A. P.; Zhu, L.; Sharon, D. A.; Ye, S.; Willard, A. P.; Zhang, X.; Zhang, B. Microphase Separation Produces Interfacial Environment within Diblock Biomolecular Condensates. eLife 2025, 12, RP90750.

(53) Wang, C.; Kilgore, H. R.; Latham, A. P.; Zhang, B. Nonspecific Yet Selective Interactions Contribute to Small Molecule Condensate Binding. Journal of Chemical Theory and Computation 2024, 20, 10247–10258.

(54) Liu, S.; Lin, X.; Zhang, B. Chromatin fiber breaks into clutches under tension and crowding. Nucleic Acids Research 2022, 50, 9738–9747.

(55) Davtyan, A.; Schafer, N. P.; Zheng, W.; Clementi, C.; Wolynes, P. G.; Papoian, G. A. AWSEM-MD: Protein Structure Prediction Using Coarse-Grained Physical Potentials and Bioinformatically Based Local Structure Biasing. The Journal of Physical Chemistry B 2012, 116, 8494–8503.

(56) Wu, H.; Wolynes, P. G.; Papoian, G. A. AWSEM-IDP: A Coarse-Grained Force Field for Intrinsically Disordered Proteins. The Journal of Physical Chemistry B 2018, 122, 11115–11125.

(57) Bereau, T.; Deserno, M. Generic Coarse-Grained Model for Protein Folding and Aggregation. The Journal of Chemical Physics 2009, 130, 235106.

(58) Sterpone, F.; Melchionna, S.; Tuffery, P.; Pasquali, S.; Mousseau, N.; Cragnolini, T.; Chebaro, Y.; St-Pierre, J.-F.; Kalimeri, M.; Barducci, A.; Laurin, Y.; Tek, A.; Baaden, M.; Nguyen, P. H.; Derreumaux, P. The OPEP Protein Model: From Single Molecules, Amyloid Formation, Crowding and Hydrodynamics to DNA/RNA Systems. Chemical Society Reviews 2014, 43, 4871–4893.

(59) Sterpone, F.; Derreumaux, P.; Melchionna, S. Protein Simulations in Fluids: Coupling the OPEP Coarse-Grained Force Field with Hydrodynamics. Journal of Chemical Theory and Computation 2015, 11, 1843–1853.

(60) Das, R.; Baker, D. Macromolecular Modeling with Rosetta. Annual Review of Biochemistry 2008, 77, 363–382.

(61) Liwo, A.; Lee, J.; Ripoll, D. R.; Pillardy, J.; Scheraga, H. A. Protein Structure Prediction by Global Optimization of a Potential Energy Function. Proceedings of the National Academy of Sciences 1999, 96, 5482–5485.

(62) Kmiecik, S.; Gront, D.; Kolinski, M.; Wieteska, L.; Dawid, A. E.; Kolinski, A. CoarseGrained Protein Models and Their Applications. Chemical Reviews 2016, 116, 7898– 7936.

(63) Sanyal, T.; Shell, M. S. Coarse-Grained Models Using Local-Density Potentials Optimized with the Relative Entropy: Application to Implicit Solvation. The Journal of Chemical Physics 2016, 145, 034109.

(64) Dama, J. F.; Jin, J.; Voth, G. A. The Theory of Ultra-Coarse-Graining. 3. CoarseGrained Sites with Rapid Local Equilibrium of Internal States. Journal of Chemical Theory and Computation 2017, 13, 1010–1022.

(65) Noid, W. G.; Chu, J.-W.; Ayton, G. S.; Krishna, V.; Izvekov, S.; Voth, G. A.; Das, A.; Andersen, H. C. The Multiscale Coarse-Graining Method. I. A Rigorous Bridge between Atomistic and Coarse-Grained Models. The Journal of Chemical Physics 2008, 128, 244114.

(66) Kar, P.; Feig, M. In Biomolecular Modelling and Simulations; Karabencheva-Christova, T., Ed.; Advances in Protein Chemistry and Structural Biology; Academic Press, 2014; Vol. 96; pp 143–180.

(67) Das, S.; Amin, A. N.; Lin, Y.-H.; Chan, H. S. Coarse-Grained Residue-Based Models of Disordered Protein Condensates: Utility and Limitations of Simple Charge Pattern Parameters. Physical Chemistry Chemical Physics 2018, 20, 28558–28574.

(68) Baul, U.; Chakraborty, D.; Mugnai, M. L.; Straub, J. E.; Thirumalai, D. Sequence Effects on Size, Shape, and Structural Heterogeneity in Intrinsically Disordered Proteins. The Journal of Physical Chemistry B 2019, 123, 3462–3474.

(69) Jussupow, A.; Bartley, D.; Lapidus, L. J.; Feig, M. COCOMO2: A Coarse-Grained Model for Interacting Folded and Disordered Proteins. Journal of Chemical Theory and Computation 2025, 21, 2095–2107.

(70) Cao, F.; von Bülow, S.; Tesei, G.; Lindorff-Larsen, K. A coarse-grained model for disordered and multi-domain proteins. Protein Science 2024, 33, e5172.

(71) Ashbaugh, H. S.; Hatch, H. W. Natively unfolded protein stability as a coil-to-globule transition in charge/hydropathy space. Journal of the American Chemical Society 2008, 130, 9536–9542.

(72) Kharche, S.; Yadav, M.; Hande, V.; Prakash, S.; Sengupta, D. Improved Protein Dynamics and Hydration in the Martini3 Coarse-Grain Model. Journal of Chemical Information and Modeling 2024, 64, 837–850.

(73) Renevey, A.; Riniker, S. Benchmarking Hybrid Atomistic/Coarse-Grained Schemes for Proteins with an Atomistic Water Layer. The Journal of Physical Chemistry B 2019, 123, 3033–3042.

(74) Rudzinski, J. F.; Noid, W. G. The Role of Many-Body Correlations in Determining Potentials for Coarse-Grained Models of Equilibrium Structure. The Journal of Physical Chemistry B 2012, 116, 8621–8635.

(75) Wang, B.; Zhang, L.; Dai, T.; Qin, Z.; Lu, H.; Zhang, L.; Zhou, F. Liquid–Liquid Phase Separation in Human Health and Diseases. Signal Transduction and Targeted Therapy 2021, 6, 1–16.

(76) Liu, S.; Wang, C.; Zhang, B. Toward Predictive Coarse-Grained Simulations of Biomolecular Condensates. Biochemistry 2025, 64, 1750–1761.

(77) Airas, J.; Zhang, B. Knowledge Distillation of a Protein Language Model Yields a Foundational Implicit Solvent Model. 2026.

(78) Noid, W. G. Perspective: Coarse-grained Models for Biomolecular Systems. The Journal of Chemical Physics 2013, 139, 090901.

(79) Ding, X. Optimizing Force Fields with Experimental Data Using Ensemble Reweighting and Potential Contrasting. The Journal of Physical Chemistry B 2024, 128, 6760– 6769.

(80) Ding, X.; Zhang, B. Contrastive Learning of Coarse-Grained Force Fields. Journal of Chemical Theory and Computation 2022, 18, 6334–6344.

(81) Thaler, S.; Zavadlav, J. Learning Neural Network Potentials from Experimental Data via Differentiable Trajectory Reweighting. Nature Communications 2021, 12, 6884.

(82) Riveros, I.; Zhang, B. NEAT-DNA: A Chemically Accurate, Sequence-Dependent Coarse-Grained Model for Large-Scale DNA Simulations. Journal of Chemical Theory and Computation 2026, 22, 3709–3719.

(83) McInnes, L.; Healy, J.; Saul, N.; Großberger, L. UMAP: Uniform Manifold Approximation and Projection. Journal of Open Source Software 2018, 3, 861.

(84) Jumper, J.; Evans, R.; Pritzel, A.; Green, T.; Figurnov, M.; Ronneberger, O.; Tunyasuvunakool, K.; Bates, R.; Žídek, A.; Potapenko, A.; others Highly accurate protein structure prediction with AlphaFold. nature 2021, 596, 583–589.

(85) Martin, E. W.; Holehouse, A. S.; Peran, I.; Farag, M.; Incicco, J. J.; Bremer, A.; Grace, C. R.; Soranno, A.; Pappu, R. V.; Mittag, T. Valence and Patterning of Aromatic Residues Determine the Phase Behavior of Prion-like Domains. Science 2020, 367, 694–699.

(86) Bremer, A.; Farag, M.; Borcherds, W. M.; Peran, I.; Martin, E. W.; Pappu, R. V.; Mittag, T. Deciphering How Naturally Occurring Sequence Features Impact the Phase Behaviours of Disordered Prion-like Domains. Nature Chemistry 2022, 14, 196–207.

(87) Alshareedah, I.; Borcherds, W. M.; Cohen, S. R.; Singh, A.; Posey, A. E.; Farag, M.; Bremer, A.; Strout, G. W.; Tomares, D. T.; Pappu, R. V.; Mittag, T.; Banerjee, P. R. Sequence-Specific Interactions Determine Viscoelasticity and Ageing Dynamics of Protein Condensates. Nature Physics 2024, 20, 1482–1491.

(88) Pedraza, E.; Tejedor, A. R.; Feito, A.; Gámez, F.; Collepardo-Guevara, R.; Sanz, E.; Espinosa, J. R. Predicting Saturation Concentrations of Phase-Separating Proteins via Thermodynamic Integration. Journal of Chemical Theory and Computation 2025, 21, 9919–9934.

(89) Armentia, L.; López, X.; De Sancho, D. Molecular Determinants of Arginine versus Lysine Cation–*π* Interactions in Biomolecular Condensates. Communications Chemistry 2026,

(90) Sancho, D. D.; López, X. Crossover in Aromatic Amino Acid Interaction Strength: Tyrosine vs. Phenylalanine in Biomolecular Condensates. eLife 2025, 14 .

(91) Rekhi, S.; Garcia, C. G.; Barai, M.; Rizuan, A.; Schuster, B. S.; Kiick, K. L.; Mittal, J. Expanding the Molecular Language of Protein Liquid–Liquid Phase Separation. Nature Chemistry 2024, 16, 1113–1124.

(92) Johnson, H. R.; Chalek, K.; Elathram, N.; Chau, A. T.; Domingo, A. R.; Aldana, J. E.; Nguyen, H.; de Loera, A.; Duarte, B. A.; Shapakidze, L.; Onofrei, D.; Debelouch- ina, G. T.; Lorenz, C. D.; Holland, G. P. Arg–Tyr Cation–*π* Interactions Drive Phase Separation and *β*-Sheet Assembly in Native Spider Dragline Silk. Proceedings of the National Academy of Sciences 2025, 122, e2523198122.

(93) Lim, J.; Kumar, A.; Low, K.; Verma, C. S.; Mu, Y.; Miserez, A.; Pervushin, K. Liquid– Liquid Phase Separation of Short Histidine- and Tyrosine-Rich Peptides: Sequence Specificity and Molecular Topology. The Journal of Physical Chemistry B 2021, 125, 6776–6790.

(94) Yu, Z.; Jacobson, M. P.; Josovitz, J.; Rapp, C. S.; Friesner, R. A. First-Shell Solvation of Ion Pairs: Correction of Systematic Errors in Implicit Solvent Models. The Journal of Physical Chemistry B 2004, 108, 6643–6654.

(95) Jia, M.; Yang, J.; Qin, Y.; Wang, D.; Pan, H.; Wang, L.; Xu, J.; Zhong, D. Determination of Protein Surface Hydration by Systematic Charge Mutations. The Journal of Physical Chemistry Letters 2015, 6, 5100–5105.

(96) Li, S.; Zhang, Y.; Chen, J. Backbone Interactions and Secondary Structures in Phase Separation of Disordered Proteins. Biochemical Society Transactions 2024, 52, 319– 329.

(97) Anbarasu, A.; Anand, S.; Babu, M. M.; Sethumadhavan, R. Investigations on C–H· · · *π* Interactions in RNA Binding Proteins. International Journal of Biological Macromolecules 2007, 41, 251–259.

(98) Calinsky, R.; Levy, Y. Aromatic Residues in Proteins: Re-Evaluating the Geometry and Energetics of *π*–*π*, Cation-*π*, and CH-*π* Interactions. The Journal of Physical Chemistry B 2024, 128, 8687–8700.

(99) Yeagle, P. L. The Membranes of Cells (Third Edition); Academic Press: Boston, 2016; pp 1–25.

(100) Lobo, S.; Najafi, S.; Shell, M. S.; Shea, J.-E. Hydrophobicity in Intrinsically Disordered Protein Force Fields: Implications for Conformational Ensembles and Protein–Protein Interactions. The Journal of Physical Chemistry B 2025, 129, 6817–6827.

(101) Najafi, S.; Lobo, S.; Shell, M. S.; Shea, J.-E. Context Dependency of Hydrophobicity in Intrinsically Disordered Proteins: Insights from a New Dewetting Free Energy-Based Hydrophobicity Scale. The Journal of Physical Chemistry B 2025, 129, 1904–1915.

(102) Miyazawa, S.; Jernigan, R. L. Estimation of Effective Interresidue Contact Energies from Protein Crystal Structures: Quasi-Chemical Approximation. Macromolecules 1985, 18, 534–552.

(103) Wimley, W. C.; White, S. H. Experimentally Determined Hydrophobicity Scale for Proteins at Membrane Interfaces. Nature Structural Biology 1996, 3, 842–848.

(104) Eisenberg, D.; Schwarz, E.; Komaromy, M. Analysis of Membrane and Surface Protein Sequences with the Hydrophobic Moment Plot. Journal of Molecular Biology 1984, 179, 125–142.

(105) Ashbaugh, H. S.; Hatch, H. W. Natively Unfolded Protein Stability as a Coil-to-Globule Transition in Charge/Hydropathy Space. Journal of the American Chemical Society 2008, 130, 9536–9542.

(106) Bachau, H.; Cormier, E.; Decleva, P.; Hansen, J. E.; Martín, F. Applications of Bsplines in Atomic and Molecular Physics. Reports on Progress in Physics 2001, 64, 1815–1943.

(107) Lindorff-Larsen, K.; Piana, S.; Dror, R. O.; Shaw, D. E. How fast-folding proteins fold. Science 2011, 334, 517–520.

(108) Robustelli, P.; Piana, S.; Shaw, D. E. Developing a molecular dynamics force field for both folded and disordered protein states. Proceedings of the National Academy of Sciences 2018, 115, E4758–E4766.

(109) Piana, S.; Robustelli, P.; Tan, D.; Chen, S.; Shaw, D. E. Development of a force field for the simulation of single-chain proteins and protein–protein complexes. Journal of chemical theory and computation 2020, 16, 2494–2507.

(110) Eastman, P.; Swails, J.; Chodera, J. D.; McGibbon, R. T.; Zhao, Y.; Beauchamp, K. A.; Wang, L.-P.; Simmonett, A. C.; Harrigan, M. P.; Stern, C. D.; others OpenMM 7: Rapid development of high performance algorithms for molecular dynamics. PLoS computational biology 2017, 13, e1005659.

(111) Regy, R. M.; Thompson, J.; Kim, Y. C.; Mittal, J. Improved coarse-grained model for studying sequence dependent phase separation of disordered proteins. Protein Science 2021, 30, 1371–1379.

(112) Ding, X.; Vilseck, J. Z.; Brooks III, C. L. Fast solver for large scale multistate Bennett acceptance ratio equations. Journal of chemical theory and computation 2019, 15, 799–802.

(113) Zhang, Z.; Liu, X.; Yan, K.; Tuckerman, M. E.; Liu, J. Unified Efficient Thermostat Scheme for the Canonical Ensemble with Holonomic or Isokinetic Constraints via Molecular Dynamics. The Journal of Physical Chemistry A 2019, 123, 6056–6079.

(114) Eastman, P.; Swails, J.; Chodera, J. D.; McGibbon, R. T.; Zhao, Y.; Beauchamp, K. A.; Wang, L.-P.; Simmonett, A. C.; Harrigan, M. P.; Stern, C. D.; Wiewiora, R. P.; Brooks, B. R.; Pande, V. S. OpenMM 7: Rapid Development of High Performance Algorithms for Molecular Dynamics. PLOS Computational Biology 2017, 13, e1005659.

(115) Martinsson, P.-G.; Rokhlin, V.; Tygert, M. A Randomized Algorithm for the De- composition of Matrices. Applied and Computational Harmonic Analysis 2011, 30, 47–68.

(116) Tipping, M. E.; Bishop, C. M. Mixtures of Probabilistic Principal Component Analysers.

(117) Bandyopadhyay, D.; Mehler, E. L. Quantitative Expression of Protein Heterogeneity: Response of Amino Acid Side Chains to Their Local Environment. Proteins: Structure, Function, and Bioinformatics 2008, 72, 646–659.

(118) Biou, V.; Gibrat, J.; Levin, J.; Robson, B.; Garnier, J. Secondary Structure Prediction: Combination of Three Different Methods. *”Protein Engineering*, Design and Selection*”* 1988, 2, 185–191.

(119) Black, S. D.; Mould, D. R. Development of Hydrophobicity Parameters to Analyze Proteins Which Bear Post- or Cotranslational Modifications. Analytical Biochemistry 1991, 193, 72–82.

(120) Cornette, J. L.; Cease, K. B.; Margalit, H.; Spouge, J. L.; Berzofsky, J. A.; DeLisi, C. Hydrophobicity Scales and Computational Techniques for Detecting Amphipathic Structures in Proteins. Journal of Molecular Biology 1987, 195, 659–685.

(121) Engelman, D. M.; Steitz, T. A.; Goldman, A. IDENTIFYING NONPOLAR TRANSBILAYER HELICES IN AMINO ACID SEQUENCES OF MEMBRANE PROTEINS. Annual Review of Biophysics 1986, 15, 321–353.

(122) Hamzi, H.; Rajabpour, A.; Roldán, É.; Hassanali, A. Learning the Hydrophobic, Hydrophilic, and Aromatic Character of Amino Acids from Thermal Relaxation and Interfacial Thermal Conductance. The Journal of Physical Chemistry B 2022, 126, 670–678.

(123) Hopp, T. P.; Woods, K. R. Prediction of Protein Antigenic Determinants from Amino Acid Sequences. Proceedings of the National Academy of Sciences 1981, 78, 3824– 3828.

(124) Jain, T.; Boland, T.; Lilov, A.; Burnina, I.; Brown, M.; Xu, Y.; Vásquez, M. Prediction of Delayed Retention of Antibodies in Hydrophobic Interaction Chromatography from Sequence Using Machine Learning. Bioinformatics 2017, 33, 3758–3766.

(125) Janin, J. Surface and inside Volumes in Globular Proteins. Nature 1979, 277, 491–492.

(126) Kyte, J.; Doolittle, R. F. A Simple Method for Displaying the Hydropathic Character of a Protein. Journal of Molecular Biology 1982, 157, 105–132.

(127) Meek, J. L. Prediction of Peptide Retention Times in High-Pressure Liquid Chromatography on the Basis of Amino Acid Composition. Proceedings of the National Academy of Sciences 1980, 77, 1632–1636.

(128) Ji, J.; Carpentier, B.; Chakraborty, A.; Nangia, S. An Affordable Topography-Based Protocol for Assigning a Residue’s Character on a Hydropathy (PARCH) Scale. Journal of Chemical Theory and Computation 2024, 20, 1656–1672.

(129) Gromiha, M. M.; Ponnuswamy, P. K. Hydrophobic Distribution and Spatial Arrangement of Amino Acid Residues in Membrane Proteins. International Journal of Peptide and Protein Research 1996, 48, 452–460.

(130) Rose, G. D.; Geselowitz, A. R.; Lesser, G. J.; Lee, R. H.; Zehfus, M. H. Hydrophobicity of Amino Acid Residues in Globular Proteins. Science 1985, 229, 834–838.

(131) Tanford, Charles. Contribution of Hydrophobic Interactions to the Stability of the Globular Conformation of Proteins. Journal of the American Chemical Society 1962, 84, 4240–4247.

