## Supporting Information for "MOFF2: A Transferable Coarse-Grained Protein Force Field for Predictive Condensate Simulations"

‡*Contributed equally to this work*

### Contents

**Explicit Energy Function** **S-3**

Bonded Potential . . . . . . . . . . . . . . . . . . . . . . . . . . . . . . . S-3

Electrostatic Potential . . . . . . . . . . . . . . . . . . . . . . . . . . . . . S-3

Pairwise Nonbonded Potential . . . . . . . . . . . . . . . . . . . . . . . . . S-4

Many-Body Potential . . . . . . . . . . . . . . . . . . . . . . . . . . . . . . S-5

Native Contact Potential for Folded Domains . . . . . . . . . . . . . . . . . S-6

**Force Field Optimization** **S-8**

Training Dataset . . . . . . . . . . . . . . . . . . . . . . . . . . . . . . . . S-8

Training Data from All-Atom Simulations . . . . . . . . . . . . . . . . . S-8

Training Data from Ensemble Reweighting for MDPs . . . . . . . . . . . . S-9

Noise Ensemble . . . . . . . . . . . . . . . . . . . . . . . . . . . . . . . . . S-11

Details of Noise Simulations . . . . . . . . . . . . . . . . . . . . . . . . S-11

Combining Umbrella Simulations into a Generalized Ensemble . . . . . . . S-14

Model Parameterization . . . . . . . . . . . . . . . . . . . . . . . . . . . . S-15

Potential Contrasting . . . . . . . . . . . . . . . . . . . . . . . . . . . . S-16

Fine-tuning with thermodynamic reweighting . . . . . . . . . . . . . . . . S-18

Ensemble reweighting for model evaluation . . . . . . . . . . . . . . . . . S-19

**Supplemental Tables** **S-21**

**Supplemental Figures** **S-38**

**References** **S-47**

### Explicit Energy Function

MOFF2 is a coarse-grained force field for protein simulations, adopting the one-bead-per-amino-acid (1BPA) representation. Its total energy is defined as

$$V_{\mathrm{MOFF2}}(\mathbf{r}) = V_{\mathrm{bond}} + V_{\mathrm{elec}} + V_{\mathrm{pair}} + V_{\mathrm{mb}}. \tag{S1}$$

Here, $V_{\mathrm{bond}}$, $V_{\mathrm{elec}}$, and $V_{\mathrm{pair}}$ correspond to the bonded, electrostatic, and pairwise nonbonded interactions, respectively, while $V_{\mathrm{mb}}$ represents a many-body potential that accounts for collective effects often neglected in 1BPA models.

#### Bonded Potential

The bonded potential is defined between consecutive amino acids along the sequence as

$$V_{\mathrm{bond}} = \sum_{i=1}^{N-1} \frac{k_{\mathrm{bond}}}{2} (r_{i,i+1} - r_0)^2, \tag{S2}$$

where $r_{i,i+1}$ is the distance between beads $i$ and $i+1$. The equilibrium bond length and force constant are set to $r_0 = 0.386$ nm and $k_{\mathrm{bond}} = 8000$ kJ mol$^{-1}$ nm$^{-2}$, respectively.

#### Electrostatic Potential

Electrostatic interactions are described using a Debye–Hückel potential:

$$V_{\mathrm{elec}} = \sum_{i<j} \frac{q_i q_j}{4\pi\epsilon_0\epsilon_{\mathrm{water}} r_{ij}} \exp(-r_{ij}/\lambda_D), \quad r_{ij} < 5\lambda_D, \tag{S3}$$

and $V_{\mathrm{elec}} = 0$ otherwise. Here, $q_i$ and $q_j$ are the charges of coarse-grained beads, $\epsilon_0$ is the vacuum permittivity, and $\epsilon_{\mathrm{water}} = 80.0$ is the dielectric constant of water. The Debye length $\lambda_D$ is given by

$$\lambda_D = \sqrt{\frac{k_B T \epsilon_0 \epsilon_{\mathrm{water}}}{2 N_A I e_c^2}}, \tag{S4}$$

where $k_B$ is the Boltzmann constant, $T$ the temperature, $N_A$ Avogadro's number, $I$ the ionic strength, and $e_c$ the elementary charge. Bonded pairs $(i, i+1)$ are excluded from electrostatic interactions.

#### Pairwise Nonbonded Potential

The pairwise nonbonded interaction accounts for both the direct amino acid contacts described by the Ashbaugh–Hatch (AH) potential[S1] and a first-solvation-shell (FSS) correction that introduces a desolvation barrier during contact formation

$$V_{\text{pair}} = \sum_{i<j} \left[ U_{\text{AH}}(r_{ij}) + U_{\text{FSS}}(r_{ij}) \right]. \tag{S5}$$

The AH term is

$$U_{\text{AH}}(r_{ij}) = \begin{cases} U_{\text{LJ}}(r_{ij}) + (1 - \lambda_{ij})\epsilon_{\text{LJ}}, & r_{ij} \leq 2^{1/6}\sigma_{ij}, \\ \lambda_{ij} U_{\text{LJ}}(r_{ij}), & 2^{1/6}\sigma_{ij} < r_{ij} < 4\sigma_{ij}, \\ 0, & r_{ij} \geq 4\sigma_{ij}, \end{cases} \tag{S6}$$

where the potential is shifted to zero at the cutoff in the implementation.

The Lennard–Jones potential is

$$U_{\text{LJ}}(r_{ij}) = 4\epsilon_{\text{LJ}} \left[ \left( \frac{\sigma_{ij}}{r_{ij}} \right)^{12} - \left( \frac{\sigma_{ij}}{r_{ij}} \right)^{6} \right]. \tag{S7}$$

Here, $r_{ij}$ is the interparticle distance, $\sigma_{ij}$ is the pair size parameter, and $\lambda_{ij}$ controls the AH interaction strength. The basal interaction strength is set to $\epsilon_{\text{LJ}} = 0.2$ kcal mol$^{-1}$, and the cutoff is $4\sigma_{ij}$. The $\sigma_{ij}$ values are identical to those used in the HPS model.[S2]

The desolvation potential is described using a Gaussian function

$$U_{\text{FSS}}(r_{ij}) = G_{ij} \exp\left[-\frac{1}{2}\left(\frac{r_{ij} - \left(2^{1/6}\sigma_{ij} + \Delta\mu\right)}{\delta}\right)^2\right] \Theta\left(r_{ij} - 2^{1/6}\sigma_{ij}\right) \Theta\left(4\sigma_{ij} - r_{ij}\right), \quad \text{(S8)}$$

where $G_{ij}$ is the amino-acid-pair-specific Gaussian corrections. $\Delta\mu = 0.25$ nm determines the displacement of the Gaussian center from the LJ minimum, and $\delta = 0.10$ nm is the Gaussian width. $\Theta$ denotes the Heaviside step function. Given 20 amino acid types, the pair-wise contact profile therefore contains 210 $\lambda_{ij}$ and 210 $G_{ij}$ parameters to be parameterized. Bonded pairs $(i, i+1)$ are excluded from nonbonded contacts.

#### Many-Body Potential

The density-dependent many-body term is expressed as

$$V_{\text{mb}} = \sum_{I=1}^{20} \sum_{i \in I} U_{\text{mb}}(\rho_i; c_I), \quad \text{(S9)}$$

where $I$ indexes the amino acid type, $\rho_i$ is the local density around particle $i$, and $c_I$ are type-specific parameters.

The local density is defined as

$$\rho_i = \sum_{j \neq i,\, r_{ij} \leq r_0 + 10/\eta} \frac{1}{2}\left[1 + \tanh(\eta(r_0 - r_{ij}))\right], \quad \text{(S10)}$$

with $\eta = 10.0$ nm$^{-1}$ and $r_0 = 0.7$ nm. Contributions from all neighboring particles within $r_0 + 10/\eta$ are included. Because bonded pairs are not excluded, for an interior residue, the two bonded neighbors are almost always close enough and typically contribute approximately 2 to $\rho_i$.

Because the functional form of $U_{\text{mb}}$ is not known a priori, it is represented by a cubic

B-spline basis:[S3]

$$U_{\mathrm{mb}}(\rho_i; c_I) = \begin{cases} \sum_{j=1}^{M+K-2} c_I^j B_{j,M}(\rho_i), & \rho_i < \rho_{\max}, \\ 0, & \rho_i \geq \rho_{\max}, \end{cases} \tag{S11}$$

where $B_{j,M}(\rho_i)$ are B-spline basis functions of order $M$, and $K$ controls the number of spline intervals. The cutoff density $\rho_{\max} = 15.0$ was chosen to encompass the maximum local densities observed in the training data. Considering 20 amino acid types, there are $20 \times (M + K - 2)$ coefficients $c_I^j$ to be optimized. In this work, a clamped B-spline basis with order $M = 4$ and $K = 10$ was used.

#### Native Contact Potential for Folded Domains

To preserve the tertiary structures of folded domains, an additional term, $V_{\mathrm{native}}$, is included in the total energy beyond the contributions in Eq. S1. The native contact energy is defined as

$$V_{\mathrm{native}} = V_{\mathrm{angle}} + V_{\mathrm{dihed}} + V_{\mathrm{nc}}. \tag{S12}$$

**Angle potential.** The angular term is expressed as

$$V_{\mathrm{angle}} = \sum_{i=1}^{N-2} U_{\mathrm{angle}}(\theta_i), \tag{S13}$$

with

$$U_{\mathrm{angle}}(\theta_i) = k_{\mathrm{angle}} M^2 \left[1 - \cos\left(\frac{\theta_i - \theta_i^0}{M}\right)\right], \tag{S14}$$

where $k_{\mathrm{angle}} = 120$ kJ mol$^{-1}$ rad$^{-2}$, $M = 5$, and $\theta_i^0$ is the reference bond angle extracted from the native structure.

**Dihedral potential.** The dihedral contribution is given by

$$V_{\mathrm{dihed}} = \sum_{i=1}^{N-3} U_{\mathrm{dihed}}(\phi_i), \tag{S15}$$

where

$$U_{\text{dihed}}(\phi_i) = \sum_{n=1,3} k_{\text{dihed},n} \left[1 + \cos\left(n(\phi_i - \phi_i^0 - \pi)\right)\right]. \tag{S16}$$

The coefficients are set to $k_{\text{dihed},1} = 3.0$ kJ mol$^{-1}$ and $k_{\text{dihed},3} = 1.5$ kJ mol$^{-1}$. The reference dihedral angle $\phi_i^0$ is determined from the native structure.

**Native contact potential.** Native tertiary interactions are modeled using a 10–12 potential,

$$V_{\text{nc}} = \sum_{\substack{i<j \\ i,j \in \text{ncList}}} \epsilon_{\text{nc}} \left[5\left(\frac{\mu_{ij}}{r_{ij}}\right)^{12} - 6\left(\frac{\mu_{ij}}{r_{ij}}\right)^{10}\right], \tag{S17}$$

where the default interaction strength is $\epsilon_{\text{nc}} = 6$ kJ mol$^{-1}$, and $\mu_{ij}$ is the C$\alpha$–C$\alpha$ distance in the reference native structure.

The native contact list (`ncList`) is constructed from the reference structure following two criteria: (1) residue pairs in contact according to the shadow algorithm,[S4] as implemented in `OpenABC`;[S5] and (2) residue pairs not directly connected by bond, angle, or dihedral interactions, i.e., $|i-j| > 3$ for residues on the same chain. For simulations used to construct noise ensembles or to evaluate model accuracy, we further restrict the contact list to residue pairs within the same continuous secondary structure element (either an $\alpha$-helix or a $\beta$-sheet), as identified using DSSP[S6] implemented in `MDTraj`.[S7]

**Exclusion of bonded pairs.** When the native contact potential is applied, atom pairs involved in bond, angle, or dihedral terms (i.e., pairs $i$ and $i+1/2/3$ on the same chain) are excluded from the evaluations of $V_{\text{pair}}$ and $V_{\text{elec}}$ to avoid double counting of short-range interactions.

### Force Field Optimization

#### Training Dataset

Training the coarse-grained force field by potential contrasting requires molecular configurations that cover the conformational and sequence regimes targeted by the model. To support transferability, we assembled a heterogeneous dataset comprising ordered proteins (OPs; Tables S1 and S2), intrinsically disordered proteins (IDPs; Table S4), and multi-domain proteins (MDPs; Table S3). The procedures used to generate these datasets are described below.

##### Training Data from All-Atom Simulations

Explicit-solvent atomistic molecular dynamics (MD) simulations can provide chemically accurate configurational ensembles and are well suited for generating training data for coarse-grained models. Accordingly, we collected configurations for both ordered and disordered proteins from long-timescale simulations.

**Ordered protein dataset.** We selected 19 ordered proteins with trajectories reported by D. E. Shaw Research.[S8–S10] To expand sequence diversity, we performed additional 1.2-$\mu$s-long simulations for 18 more proteins.

Initial structures were obtained either from the Protein Data Bank (PDB) or from AlphaFold2 predictions.[S11,S12] Each protein was placed in a dodecahedral water box with a minimum distance of 1.0 nm between the protein and the box boundary. Sodium and chloride ions were added to achieve a 150 mM salt concentration. Simulations were carried out in OpenMM[S13] using the a99SB-*disp* force field[S8] under NPT conditions (1 atm, 300 K). Hydrogen mass repartitioning was employed to enable a 4 fs integration timestep. The Langevin middle integrator[S14] was used with a friction coefficient of 1 ps$^{-1}$. Each simulation lasted 1.2 $\mu$s, and configurations were saved every 40 ps. Only the final 1.0 $\mu$s of each trajectory was used for training.

**Disordered protein dataset.** We included seven IDPs from Ref. S8 and 34 additional IDPs from Ref. S15. A subset of sequences from the latter dataset was excluded because they exhibited limited conformational variability, as indicated by a narrow range of radius of gyration ($R_g$) values, and therefore contributed little additional sampling diversity.

##### Training Data from Ensemble Reweighting for MDPs

To parameterize interactions between folded domains, disordered linkers, and domain surfaces, training data for MDPs were also included. Explicit-solvent atomistic simulations of MDPs are computationally prohibitive because of their large system sizes and slow conformational transitions, which limit equilibrium sampling. We therefore used experimentally reweighted coarse-grained ensembles as MDP training data.

We first generated coarse-grained simulations of the MDPs, as described in Details of Noise Simulations. Starting from these configurational ensembles, we applied a reweighting scheme to assign statistical weights to each configuration so that ensemble averages matched experimental observables. In this work, we targeted the experimental mean radius of gyration ($R_g$), although the method can, in principle, accommodate other thermodynamic observables.

Given a reference potential $u_0$ that defines the original ensemble $p_0 \propto \exp(-\beta u_0)$, the ensemble average of an observable $Y$ is

$$\langle Y \rangle_0 = \int p_0(x) Y(x)\, \mathrm{d}x \approx \frac{1}{N_0} \sum_{i=1}^{N_0} Y(x_i^{(0)}), \tag{S18}$$

where $x_i^{(0)}$ is the $i$-th configuration sampled from $p_0$. When $\langle Y \rangle_0$ deviates from the experimental value $\langle Y \rangle_{\mathrm{exp}}$, we reassign normalized weights $w_i^{\mathrm{ME}}$ to the existing configurations

$\{x_i^{(0)}\}$ such that the reweighted ensemble satisfies

$$1 = \sum_{i=1}^{N_0} w_i^{\mathrm{ME}}, \tag{S19}$$

$$\langle Y \rangle_{\mathrm{exp}} = \sum_{i=1}^{N_0} w_i^{\mathrm{ME}} Y(x_i^{(0)}). \tag{S20}$$

The reweighted ensemble is required to deviate minimally from the original one. This is achieved by maximizing the Shannon entropy,

$$S = -\sum_{i=1}^{N_0} w_i \log w_i, \tag{S21}$$

which is equivalent to minimizing the Kullback–Leibler (KL) divergence between $\{w_i\}$ and the uniform weights $w_i^{(0)} = 1/N_0$:

$$\mathrm{KL}(w_i || w_i^{(0)}) = \sum_{i=1}^{N_0} w_i \log\left(\frac{w_i}{w_i^{(0)}}\right) = \log N_0 - S. \tag{S22}$$

Maximizing $S$ therefore minimizes $\mathrm{KL}(w_i||w_i^{(0)})$, ensuring minimal perturbation to the original distribution.

In practice, we implemented this optimization using the `scipy` package[S16] with the "trust-constr" algorithm to minimize $-S$ under the constraints in Eqs. S19–S20. To avoid numerical instability, a small lower bound $w_i \geq 10^{-10}$ was applied. Analytical gradients, $\partial(-S)/\partial w_i = \log w_i + 1$, and the Hessian matrix, $\partial^2(-S)/(\partial w_i \partial w_j) = \delta_{ij}/w_i$, were supplied to improve convergence.

After determining the optimal weights $w_i^{\mathrm{ME}}$, configurations $\{x_i^{(0)}\}$ were resampled according to these weights to generate a new ensemble $\{x_i^{(1)}\}$ with $N_1$ configurations. The resulting reweighted ensembles, which reproduce experimental $R_g$ values, were used as training data for the MDP systems.

#### Noise Ensemble

In addition to the training data, potential contrasting requires a complementary set of *noise* configurations. For stable optimization, the noise ensemble must span a configurational space that overlaps with, but is not identical to, the data ensemble. As demonstrated in Ref. S17, this condition can be satisfied by constructing generalized ensembles that combine configurations from umbrella-sampling simulations. The following subsections describe the procedures used to generate noise ensembles for each protein class.

##### Details of Noise Simulations

All noise simulations were performed using a custom implementation of the HPS model following Ref. S2. The protein termini were left uncapped, resulting in a net charge of +1 and −1 for the N- and C-termini, respectively. Simulations were carried out in the NVT ensemble at the specified target temperatures and ionic strengths (see below). The Langevin middle integrator[S14] was employed with a friction coefficient of 1 $ps^{-1}$ and a timestep of 10 fs. Each simulation was equilibrated for 5 ns prior to the production run.

**Noise Simulations for IDPs.** For IDPs, conformational sampling was performed using the following energy function:

$$V_{\text{HPS}}(\mathbf{r}) = V_{\text{bond}} + V_{\text{pair}} + V_{\text{elec}}. \tag{S23}$$

The individual terms are defined in Eqs. S2–S3. Pairwise hydrophobicity parameters, $\lambda_{ij}$, were adopted directly from Ref. S2. The Debye length for the electrostatic term was computed from the temperature and ionic strength of the corresponding all-atom simulations (Table S4), and a cutoff of 4.0 nm was applied.

Enhanced sampling was achieved by applying an umbrella bias along the radius of gyration ($R_g$):

$$U^i_{\text{bias}} = \frac{\kappa_i}{2}\left[R_g(\mathbf{r}) - R^i_g\right]^2, \tag{S24}$$

where $\kappa_i$ and $R^i_g$ are the force constant and umbrella center of the $i$th window, respectively. The radius of gyration is computed as

$$R_g(\mathbf{r}) = \sqrt{\frac{\sum_{i=1}^{n} m_i |\vec{r}_i - \vec{r}_{\text{COM}}|^2}{\sum_{i=1}^{n} m_i}}, \tag{S25}$$

where $\vec{r}_i$ denotes the position of bead $i$, $n$ is the protein sequence length, and the center of mass is given by

$$\vec{r}_{\text{COM}} = \frac{\sum_{i=1}^{n} m_i \vec{r}_i}{\sum_{i=1}^{n} m_i}. \tag{S26}$$

Umbrella centers $R^i_g$ were distributed to cover a broad range of compact and extended conformations, ensuring overlap with the data ensemble while still sampling configurations outside the high-probability data region. Each simulation used the same salt concentration as the corresponding all-atom simulation and was initialized from the reference structure described in the *Training Dataset* section. The number of umbrella windows, force constants, and bias centers are listed in Table S15. Each simulation lasted $5 \times 10^7$ steps with a 10 fs timestep.

**Noise Simulations for Ordered Proteins.** For ordered proteins, we employed the same HPS model with the inclusion of a native potential, yielding

$$V_{\text{OP}}(\mathbf{r}) = V_{\text{HPS}} + V_{\text{native}}, \tag{S27}$$

where $V_{\text{native}}$ is defined in Eq. S12. The electrostatic cutoff was set to $5\lambda_D$ to account for the varying salt concentrations used in all-atom simulations that produce heterogeneous screening lengths.

Reference structures for constructing $V_{\text{native}}$ were selected from all-atom trajectories via hierarchical clustering. Using SciPy,[S16] we converted each all-atom trajectory to a C$\alpha$ representation and computed the pairwise RMSD matrix, with element $(i, j)$ representing the RMSD between frames $i$ and $j$. Average-linkage hierarchical clustering was applied to this matrix, followed by flat clustering into at most 50 clusters. The folded reference structure was chosen from the largest cluster as the frame with minimal mean RMSD relative to all other members. This structure was used to define reference values for bond angles, dihedral angles, and native contacts (Eqs. S14–S17).

As mentioned above, the native contact list (`ncList`) used in these simulations is restricted to residue pairs within the same continuous secondary structure element, either an $\alpha$-helix or a $\beta$-sheet. Importantly, this restricted contact set does not impose biases on tertiary structure formation. As a result, tertiary contacts arise solely from the nonbonded interactions described by the potential $V_{\text{pair}}$. This design enables MOFF2 to parameterize $V_{\text{pair}}$ in a manner that accurately stabilizes tertiary interactions and promotes the formation of collapsed conformations in folded proteins.

To enhance sampling, we introduced an umbrella bias along the RMSD from the reference structure:

$$U^i_{\text{bias}} = \frac{\kappa_i}{2}\left[\text{RMSD}(\mathbf{r}) - \text{RMSD}_i\right]^2, \tag{S28}$$

where $\kappa_i$ and $\text{RMSD}_i$ denote the bias strength and center of the $i$th window, respectively. Bias centers were distributed to span the RMSD range observed in all-atom trajectories, ensuring broad configurational coverage. Simulation temperatures and salt concentrations matched those of the atomistic simulations, with further details provided in the *Training Dataset* section. The number of umbrella windows, force constants, and bias centers are listed in Table S14. Each simulation lasted $5 \times 10^7$ steps with a 10 fs timestep.

**Noise Simulations for Multi-Domain Proteins.** For MDPs, disordered regions were modeled using $V_{\text{HPS}}(\mathbf{r})$ (Eq. S23), while folded domains followed $V_{\text{OP}}(\mathbf{r})$ (Eq. S27). Folded regions were identified based on AlphaFold pLDDT scores $\geq 70$. Unlike the case of single-domain proteins, native contacts here included all residue pairs identified by the shadow algorithm, regardless of secondary structure continuity, to capture global stabilization of folded domains. The native contact strength was set to 6 kJ mol$^{-1}$ to maintain structural integrity of each folded domain while allowing flexible interdomain interactions. This setup focuses the learning process on the interactions between ordered domain surfaces and disordered linkers rather than on local secondary structure stabilization.

Simulations were performed at the experimental temperatures and ionic strengths summarized in Table S16. Each trajectory was propagated for $5 \times 10^7$ steps with a 10 fs timestep.

##### Combining Umbrella Simulations into a Generalized Ensemble

For both ordered proteins and IDPs, configurations from multiple umbrella simulations were merged into a generalized ensemble,

$$G = \{x_{j=1,\ldots,N_i}^{(i=1,\ldots,M)}\},$$

with distribution

$$p_0(x) = \sum_{i=1}^{M} P(\lambda = i, x) = \sum_{i=1}^{M} \exp\left[-\beta(u_i(x) + v_i)\right], \tag{S29}$$

where $M$ is the number of umbrella windows and $N_i$ is the number of samples collected from the $i$-th window. The corresponding reduced potential is

$$u_0(x) = -k_B T \ln p_0(x) + \text{const.} = -\ln\left(\sum_{i=1}^{M} \exp[-\beta(u_i(x) + v_i)]\right) + \text{const.}, \tag{S30}$$

indicating that all noise samples in the generalized ensemble, $G$, can be regarded as drawn from the effective potential $u_0$.

The reweighting factors $v_i$ were determined using the multistate Bennett acceptance ratio (MBAR).[S18,S19] Assuming samples collected from $M$ thermodynamic states characterized by potentials $u_i$, the joint probability is

$$P(\lambda = i, x) = \exp[-\beta(u_i(x) + v_i)], \tag{S31}$$

where $\lambda$ denotes the state index. The probability that configuration $x$ originates from state $i$ is

$$P(\lambda = i|x) = \frac{\exp[-\beta(u_i(x) + v_i)]}{\sum_{j=1}^{M} \exp[-\beta(u_j(x) + v_j)]}. \tag{S32}$$

Since $P(x) = 1/N$ for the full ensemble, the marginal probability $P(\lambda = i) = N_i/N$ satisfies

$$N_i = \sum_{x \in G} \frac{\exp[-\beta(u_i(x) + v_i)]}{\sum_{j=1}^{M} \exp[-\beta(u_j(x) + v_j)]}. \tag{S33}$$

Equation S33 constitutes the MBAR self-consistency condition and can be solved iteratively by minimizing the MBAR loss function:

$$\mathcal{L}_{\mathrm{MBAR}}(v_1, \ldots, v_M) = \frac{1}{N} \sum_{x \in G} \ln \left[ \sum_{i=1}^{M} e^{-\beta(u_i(x) + v_i)} \right] + \sum_{i=1}^{M} \frac{N_i v_i}{N}. \tag{S34}$$

This procedure yields the optimal set of $v_i$ that balance the relative weights among all umbrella windows, forming a unified generalized noise ensemble for subsequent potential contrasting.

#### Model Parameterization

MOFF2 contains three classes of optimized parameters. The pairwise interaction parameters include 210 unique AH interaction coefficients $\lambda_{ij}$ (Eq. S6) and 210 unique Gaussian first-solvation-shell amplitudes $G_{ij}$ (Eq. S8), corresponding to all symmetric amino-acid pairs among 20 residue types. We collectively denote these pairwise parameters as

$$\boldsymbol{\theta}_{\mathrm{pair}} = \{\lambda_{ij}, G_{ij}\}.$$

The density-dependent many-body term contains 240 amino-acid-specific B-spline coefficients,

$$\boldsymbol{\theta}_{\rm mb} = \{c_i^j\},$$

corresponding to 20 amino-acid types and 12 learned spline coefficients per amino acid (Eq. S11).

As described in the main text, MOFF2 parameters were optimized in two stages. First, potential contrasting was used to learn a global energy function that differentiates configurations sampled from noise ensembles from configurations in reference atomistic ensembles. Second, thermodynamic reweighting was used to locally refine the parameters against experimental observables. Below, we summarize the parameterization details used in each stage.

##### Potential Contrasting

Potential contrasting was used to learn a global MOFF2 energy function from paired reference and noise ensembles. For each training protein, reference configurations were sampled from atomistic or high-resolution conformational ensembles, whereas noise configurations were generated from coarse-grained simulations designed to broadly overlap with the reference ensemble. The objective was to assign lower energies to reference configurations than to noise configurations under the learned MOFF2 potential, thereby optimizing the force field to distinguish physically relevant conformations from perturbed or less favorable configurations.

Because the training set contains three protein classes with different numbers of systems and different conformational characteristics, the total potential-contrasting loss was written as a weighted sum of class-specific losses:

$$\mathcal{L}_{\rm PC}(\boldsymbol{\theta}) = w_{\rm IDP}\mathcal{L}_{\rm IDP}(\boldsymbol{\theta}) + w_{\rm OP}\mathcal{L}_{\rm OP}(\boldsymbol{\theta}) + w_{\rm MDP}\mathcal{L}_{\rm MDP}(\boldsymbol{\theta}) + \mathcal{R}(\boldsymbol{\theta}),$$

where $\mathcal{L}_{\rm IDP}$, $\mathcal{L}_{\rm OP}$, and $\mathcal{L}_{\rm MDP}$ are the potential-contrasting losses for IDPs, OPs, and MDPs, respectively, and $w_{\mathrm{IDP}}$, $w_{\mathrm{OP}}$, and $w_{\mathrm{MDP}}$ are class weights used to balance the influence of the three protein classes during training.

In addition to the standard potential-contrasting loss function,[S17] we included regularization terms to control the magnitude of pairwise parameter updates and density-dependent many-body coefficients:

$$\mathcal{R}(\boldsymbol{\theta}) = \frac{\zeta_{\mathrm{pair}}}{2}\left[\frac{1}{210}\sum_{i=1}^{210}(\Delta\lambda_i)^2 + \frac{1}{210}\sum_{i=1}^{210}(G_i)^2\right] + \zeta_{\mathrm{mb}}\frac{1}{240}\sum_{i=1}^{240}(c_i)^2.$$

Here, $\Delta\lambda_i$ is the learned update to the initial AH pair coefficient, $G_i$ is the learned Gaussian first-solvation-shell correction, and $c_i$ denotes a density-dependent B-spline coefficient. The regularization term penalizes overly large pairwise and many-body parameters, reducing overfitting and improving stability when fitting heterogeneous OP, IDP, and MDP ensembles.

Potential contrasting was carried out using the open-source implementation available at `PCCG GitHub`. The optimization used the training reference ensembles and corresponding noise ensembles described above, including 37 OPs, 41 IDPs, and 13 MDPs (Tables S1–S4).

Because the performance of the potential-contrasting model depends on the choice of hyperparameters ($\{w_{\mathrm{IDP}}, w_{\mathrm{OP}}, w_{\mathrm{MDP}}, \zeta_{\mathrm{pair}}, \zeta_{\mathrm{mb}}\}$), we performed a systematic hyperparameter scan to identify the optimal parameterization. In addition to varying the hyperparameter values, we also evaluated alternative energy-function formulations. Specifically, we compared models containing only the pairwise contact potential ($V_{\mathrm{AH}}$), models combining the pairwise contact and density-dependent many-body potentials ($V_{\mathrm{AH}} + V_{\mathrm{mb}}$), and models employing the full MOFF2 energy function.

To assess model performance efficiently, we estimated the corresponding ($\langle R_g \rangle$) values for proteins in the training set using the ensemble-reweighting procedure described below (Ensemble reweighting for model evaluation), thereby avoiding computationally expensive molecular dynamics simulations for each candidate model. As shown in Figure S2, the full model containing $V_{\mathrm{AH}}$, $V_{\mathrm{FSS}}$, and $V_{\mathrm{mb}}$ achieved the best balance of rRMSE across OPs,

IDPs, and MDPs. The final selected hyperparameters were $\{w_{\text{IDP}} = 0.5, w_{\text{OP}} = 1.0, w_{\text{MDP}} = 1.0, \zeta_{\text{pair}} = 1.0, \text{and } \zeta_{\text{mb}} = 0.005\}$. We refer to the resulting force field obtained from potential contrasting as MOFF2-pc.

#### Fine-tuning with thermodynamic reweighting

To further improve MOFF2-pc transferability across sequence space without constructing new reference configurational ensembles, we used a thermodynamic reweighting procedure based on experimental $R_g$ values for selected proteins (Tables S7–S9) following Refs. S20,S21.

For each target protein, we first generated a reference ensemble $P_0$ using direct MD simulations with the MOFF2-pc model. We denote the corresponding potential energy as $U_0(\mathbf{x})$. For a new parameter set $\boldsymbol{\theta}$, the $R_g$ expectation under the perturbed energy function $U_1(\mathbf{x}|\boldsymbol{\theta})$ was estimated by reweighting configurations sampled from $P_0$:

$$\langle R_g(\boldsymbol{\theta})\rangle_{P_1} = \frac{\sum_{i=1}^{N} R_g(\mathbf{x}_i) w(\mathbf{x}_i|\boldsymbol{\theta})}{\sum_{i=1}^{N} w(\mathbf{x}_i|\boldsymbol{\theta})}, \tag{S35}$$

where $\{\mathbf{x}_i\}_{i=1}^{N} \sim P_0$, and the unnormalized reweighting factor is

$$w(\mathbf{x}|\boldsymbol{\theta}) = \exp[-\beta(U_1(\mathbf{x}|\boldsymbol{\theta}) - U_0(\mathbf{x}))]. \tag{S36}$$

Because Eq. S35 is differentiable with respect to $\boldsymbol{\theta}$, the parameters can be optimized directly to minimize the discrepancy between predicted and experimental $R_g$ values.

We included an effective-sample-size (ESS)[S20,S21] regularizer to maintain sufficient overlap between the reference ensemble $P_0$ and the reweighted ensemble $P_1$. For each protein $p$, the ESS was computed from the unnormalized reweighting weights as

$$\text{ESS}^p(\boldsymbol{\theta}) = \frac{\left[\sum_{i=1}^{N_p} w(\mathbf{x}_{p,i}|\boldsymbol{\theta})\right]^2}{\sum_{i=1}^{N_p} w(\mathbf{x}_{p,i}|\boldsymbol{\theta})^2}. \tag{S37}$$

The ESS penalty was then defined as

$$R_{\mathrm{ESS}}^{p}(\boldsymbol{\theta}) = \begin{cases} 0, & \mathrm{ESS}^{p}(\boldsymbol{\theta}) \geq \mathrm{ESS}_0, \\ \alpha \left[\mathrm{ESS}^{p}(\boldsymbol{\theta}) - \mathrm{ESS}_0\right]^2, & \mathrm{ESS}^{p}(\boldsymbol{\theta}) < \mathrm{ESS}_0. \end{cases} \tag{S38}$$

Here, the hyperparameter $\mathrm{ESS}_0 = 250$ defines the minimum acceptable effective sample size required for reliable reweighting, and $\alpha = 300$ controls the strength of the penalty when the reweighted ensemble falls below this threshold. Thus, the ESS regularizer prevents the optimization from selecting parameter updates for which the observable estimate is dominated by only a small number of configurations, keeping the fine-tuned model within the region of configurational overlap sampled by the MOFF2-pc reference ensemble.

For each protein $p$, the loss function was defined as

$$\ell_{\mathrm{OM}}^{p}(\boldsymbol{\theta}) = \left(\langle R_g^{p}(\boldsymbol{\theta})\rangle_{P_1} - R_{g,p}^{\mathrm{exp}}\right)^2 + R_{\mathrm{ESS}}^{p}(\boldsymbol{\theta}). \tag{S39}$$

The total fine-tuning loss was averaged over all proteins and included an additional $L_2$ penalty on changes to the density-dependent spline coefficients:

$$\mathcal{L}_{\mathrm{OM}}(\boldsymbol{\theta}) = \frac{1}{N_{\mathrm{prot}}} \sum_{p=1}^{N_{\mathrm{prot}}} \ell_{\mathrm{OM}}^{p}(\boldsymbol{\theta}) + \lambda_{\mathrm{density}} \left\|\Delta\boldsymbol{\theta}_{\mathrm{mb}}\right\|_2^2, \tag{S40}$$

where $N_{\mathrm{prot}}$ is the number of proteins included in reweighting (Tables S7–S9) and $\lambda_{\mathrm{density}} = 10^{-2}$. The $L_2$ penalty was applied only to updates of the density-spline coefficients, while no additional penalty was applied to the $V_{\mathrm{AH}}$ or $V_{\mathrm{FSS}}$ pairwise parameter updates. This choice was used to stabilize the fine-tuning process.

##### Ensemble reweighting for model evaluation

To evaluate models trained with different hyperparameters without performing new molecular dynamics simulations for every candidate model, we used thermodynamic reweighting based on reference ensembles. For each optimized parameter set, the ensemble-averaged radius of gyration, $\langle R_g \rangle$, of each training protein was estimated using free-energy perturbation theory.[S22]

Let $u_0(\mathbf{x})$ denote the reduced potential energy of the original ensemble (either generalized noise ensembles for potential contrasting or MOFF2-pc ensembles for fine-tuning), and let $u_1(\mathbf{x};\boldsymbol{\theta}^*)$ denote the reduced potential energy of a candidate optimized model with parameters $\boldsymbol{\theta}^*$. The ensemble average of $R_g$ under the candidate model is

$$\langle R_g \rangle_1 = \frac{\int R_g(\mathbf{x}) e^{-u_1(\mathbf{x};\boldsymbol{\theta}^*)}\, d\mathbf{x}}{\int e^{-u_1(\mathbf{x};\boldsymbol{\theta}^*)}\, d\mathbf{x}}. \tag{S41}$$

The expression can be rewritten using ensemble averages of $u_0(\mathbf{x})$ as follows:

$$\langle R_g \rangle_1 = \frac{\int R_g(\mathbf{x}) e^{-[u_1(\mathbf{x};\boldsymbol{\theta}^*) - u_0(\mathbf{x})]} e^{-u_0(\mathbf{x})}\, d\mathbf{x}}{\int e^{-[u_1(\mathbf{x};\boldsymbol{\theta}^*) - u_0(\mathbf{x})]} e^{-u_0(\mathbf{x})}\, d\mathbf{x}}. \tag{S42}$$

Thus, for configurations sampled from the reference ensemble, the candidate-model average can be estimated as

$$\langle R_g \rangle_1 \approx \frac{\left\langle R_g(\mathbf{x}) e^{-[u_1(\mathbf{x};\boldsymbol{\theta}^*) - u_0(\mathbf{x})]} \right\rangle_0}{\left\langle e^{-[u_1(\mathbf{x};\boldsymbol{\theta}^*) - u_0(\mathbf{x})]} \right\rangle_0}, \tag{S43}$$

where $\langle \cdot \rangle_0$ and $\langle \cdot \rangle_1$ denote ensemble averages over the reference and the candidate optimized-model ensemble, respectively. Here, $u_0$ and $u_1$ are reduced energies.

This reweighting procedure enabled rapid comparison of candidate models during hyperparameter selection. For each candidate parameter set, the reweighted $\langle R_g \rangle_1$ values were compared with the corresponding target values. To assess the reliability of the reweighting-based evaluation, we further compared reweighted $\langle R_g \rangle$ estimates with values obtained from direct MD simulations for representative systems. As shown in Figure S9, the reweighted values accurately reproduce those from MD simulations, supporting their use for hyperparameter screening.

### Supplemental Tables

Table S1: **Details of the OP training set collected from existing literature used for potential contrasting.** $N$ is the sequence length, $T$ is the simulation temperature, $I$ is the ionic strength, and $t$ is the trajectory length. TRE denotes temperature replica-exchange simulations. DES-Amber, a99SB-*disp*, and CHARMM22*/TIP3P trajectories were produced by D. E. Shaw Research and reported in Piana et al.[S9], Robustelli et al.[S8], and Lindorff-Larsen et al.[S10], respectively. The noise ensemble generation for OPs listed below are shown in Table S14.

| Protein | $N$ | $T$ (K) | $I$ (mM) | $t$ ($\mu$s) | Force field |
|---|---|---|---|---|---|
| $\alpha$3D | 73 | TRE | 3.11 | 1000 | DES-Amber |
| BBA | 28 | TRE | 33.04 | 1000 | DES-Amber |
| BBL | 47 | TRE | 199.58 | 1000 | DES-Amber |
| engrailed | 56 | TRE | 280.46 | 1000 | DES-Amber |
| gpw | 62 | TRE | 36.54 | 1000 | DES-Amber |
| $\lambda$-repressor | 80 | TRE | 51.04 | 1000 | DES-Amber |
| NTL9 | 39 | TRE | 121.46 | 1000 | DES-Amber |
| Protein B | 47 | TRE | 51.44 | 1000 | DES-Amber |
| BPTI | 58 | 300 | 37.02 | 20 | DES-Amber |
| calmodulin | 147 | 300 | 249.98 | 30 | DES-Amber |
| Ubiquitin | 76 | 300 | 150 | 10 | a99SB-*disp* |
| GB3 | 56 | 300 | 9.98 | 10 | a99SB-*disp* |
| HEWL | 129 | 300 | 26.63 | 10 | a99SB-*disp* |
| Chignolin | 10 | 340 | 25.9 | 106 | CHARMM22*/TIP3P |
| Trp-cage | 20 | 290 | 65 | 208 | CHARMM22*/TIP3P |
| Villin | 35 | 360 | 40 | 120 | CHARMM22*/TIP3P |
| WW domain | 35 | 360 | 7.1 | 1137 | CHARMM22*/TIP3P |
| Homeodomain | 52 | 360 | 45 | 327 | CHARMM22*/TIP3P |
| Protein G | 56 | 350 | 100 | 1154 | CHARMM22*/TIP3P |

Table S2: **Details of the OP training set generated in house used for potential contrasting.** Simulations were performed at $T = 300$ K and $I = 150$ mM NaCl using a99SB-*disp* in OpenMM.[S13] Each simulation used a 4 fs timestep and lasted $3 \times 10^8$ steps, corresponding to a 1.2 $\mu$s trajectory. Configurations were saved every $10^4$ steps, giving $3 \times 10^4$ snapshots per trajectory. The final 1.0 $\mu$s, corresponding to $2.5 \times 10^4$ snapshots, was used for potential contrasting training. Protein identities are shown as PDB or UniProt IDs. Structures identified by UniProt ID were obtained from the AlphaFold2 database[S12] and selected following Airas and Zhang[S23]. $N$ denotes the protein chain length. The noise ensemble generation for OPs listed below are shown in Table S14.

| Protein | $N$ |
|---|---|
| 1soy | 106 |
| 1wla | 153 |
| 2ea9 | 103 |
| 5tvz | 103 |
| P0C232 | 100 |
| P0CG98 | 100 |
| P00251 | 100 |
| P21149 | 100 |
| P21318 | 100 |
| P29669 | 100 |
| P31960 | 100 |
| P32729 | 100 |
| P61734 | 100 |
| P69995 | 100 |
| P75202 | 100 |
| P75459 | 100 |
| P80353 | 100 |
| P87285 | 100 |

Table S3: **Details of the MDP training set used for potential contrasting.** $N$ is the total number of residues, $T$ is the temperature, and $I$ is the ionic strength. The ionic strength of the buffer is computed with `http://phbuffers.org/BufferCalc/Buffer.html`. The experimental average ($\langle R_g \rangle$) values are taken from Ref. S33. The ordered domains (ODs) are specified using 1-based residue-indices and were assigned based on secondary structure annotations . "$\Delta$" indicates special case: (1) For THB-C2 (named ΔmC2 in Michie et al.[S24]), the sequence is PDB 5K6P without the first 4 residues. (2) For Ub2 and Ub3, 1UBQ is the structure of a single ubiquitin. (3) For Gal3, the SAXS ionic strength is not specified in Lin et al.[S26], thus we use protein expression ionic strength here as an approximation. (4) hnRNPA1* is hnRNPA1 (UniProt ID: P09651) without residues 251-302 and 311-316. Temperature and $\langle R_g \rangle$ are directly read from Cao et al.[S33] as the values are not reported in Martin et al.[S27]. (5) The full sequence of SH4UD-SH3-SH2 is reported in Gurumoorthy et al.[S29]. P12931 residue 2-79 is SH4UD, which is disordered. P12931 residue 86-250 is SH3-SH2. An additional sequence LPETG connects SH4UD and SH3-SH2. SAXS temperature and buffer condition are not specified in Gurumoorthy et al.[S29] so we applied the values in Cao et al.[S33]. (6) Wild-type TDP43 sequence comes from Q13148 and all tryptophan residues are mutated to alanine to get $\text{TDP43}_{\text{WtoA}}$. SAXS $T$ and buffer condition are not specified in Wright et al.[S30] so we use the values in Cao et al.[S33]. (7) D12, D23, and D34 are subchains of PDZK1 (UniProt ID: Q5T2W1). (8) For SMAD4, the SAXS buffer condition is not specified so the buffer condition is estimated based on purification buffer in Gomes et al.[S32]. Additional explanations for FPs-$\text{GS}_n$ ($n$ = 8 or 16) are summarized in *Noise Ensemble*. The noise ensemble generation for MDPs listed below are shown in Table S16.

| Protein | ID | $N$ | $T$ (K) | $I$ (mM) | $\langle R_g \rangle$ (nm) | ODs | Ref |
|---|---|---|---|---|---|---|---|
| THB-C2 | 5K6P$^{\Delta}$ | 133 | 295.15 | 146 | 1.99 | 2-38, 47-133 | S24 |
| Ub2 | 1UBQ$^{\Delta}$ | 152 | 293 | 332 | 2.034 | 1-70, 77-146 | S25 |
| Ub3 | 1UBQ$^{\Delta}$ | 228 | 293 | 332 | 2.699 | 1-70, 77-146, 153-222 | S25 |
| Gal3 | P17931 | 250 | 303 | 337$^{\Delta}$ | 2.892 | 113-250 | S26 |
| hnRNPA1* | P09651$^{\Delta}$ | 314 | 293.15$^{\Delta}$ | 150 | 3.12$^{\Delta}$ | 9-189 | S27 |
| FPs-$\text{GS}_8$ | A0A1S4NYF2$^{\Delta}$ | 491 | 293.15 | 149 | 3.45 | 2-232, 266-484 | S28 |
| FPs-$\text{GS}_{16}$ | A0A1S4NYF2$^{\Delta}$ | 507 | 293.15 | 149 | 3.546 | 2-232, 282-500 | S28 |
| SH4UD-SH3-SH2 | P12931$^{\Delta}$ | 248 | 293.15$^{\Delta}$ | 216$^{\Delta}$ | 3.28 | 85-248 | S29 |
| $\text{TDP43}_{\text{WtoA}}$ | Q13148$^{\Delta}$ | 414 | 293.15$^{\Delta}$ | 312$^{\Delta}$ | 4.11 | 3-79, 104-178, 191-261 | S30 |
| D12 | Q5T2W1$^{\Delta}$ | 213 | 283.15 | 156 | 2.95 | 1-110, 127-213 | S31 |
| D23 | Q5T2W1$^{\Delta}$ | 192 | 283.15 | 156 | 3.08 | 1-87, 107-192 | S31 |
| D34 | Q5T2W1$^{\Delta}$ | 222 | 283.15 | 156 | 3.34 | 1-86, 136-222 | S31 |
| SMAD4 | Q13485 | 552 | 283.15 | 188$^{\Delta}$ | 4.7 | 9-137, 287-296, 312-463, 491-543 | S32 |

Table S4: **Details of the IDP training set used for potential contrasting.** $N$ is the sequence length, $T$ is the simulation temperature, $I$ is the ionic strength, and $t$ is the trajectory duration. Simulation trajectories for the first 7 proteins were reported by Robustelli et al.[S8]. $IDP_{WZ}$ represents 34 IDPs selected from the dataset reported by Wang and Zhang[S15], with their sequence details listed in Table S5. The noise ensemble generation of IDPs listed below is shown in Table S15.

| Protein | $N$ | $T$ (K) | $I$ (mM) | $t$ ($\mu$s) | Force field |
|---|---|---|---|---|---|
| ACTR | 71 | 300 | 150 | 30 | a99SB-*disp* |
| A$\beta$40 | 40 | 300 | 50 | 30 | a99SB-*disp* |
| Ash1 | 83 | 300 | 150 | 30 | a99SB-*disp* |
| $N_{tail}$ | 132 | 300 | 100 | 30 | a99SB-*disp* |
| drkN SH3 | 59 | 300 | 50 | 30 | a99SB-*disp* |
| p15PAF | 110 | 300 | 50 | 30 | a99SB-*disp* |
| sic1 | 92 | 300 | 150 | 30 | a99SB-*disp* |
| $IDP_{WZ}$ (34) | 40 | 300 | 150 | 10.5 | a99SB-*disp* |

Table S5: **Protein sequences of the IDP$_{\mathbf{WZ}}$ dataset from Ref. S15 used for potential contrasting training.**

| No. | Sequence |
|---|---|
| 1 | ELDPE SEVQA VGTGT KTDSD STKAT SAETA GKTSS AILED |
| 2 | QSISS RVSYN LDICF LQQAI SSIND FKGLE NKYCV LGREA |
| 3 | ITHET RTTPS VNTIN KSNPQ GPSDN ATSIK KTLNR VPKWM |
| 4 | DWALE VENEE IEQTQ PTPLE GEDEE EGLDS EFKKK ITLQD |
| 5 | DCNFL DDSTV TNISA NHSIT SLVNE WAEYD LLSDT DLSSN |
| 6 | FLEEC REKEE FVNES TDDEE SKFDI ADYKD EMVSD EHLWN |
| 7 | EESLI ESEEA TREEV DDFVG SDDAV ALGDM RPQQD YGRIL |
| 8 | RFKNN NHTIS VFDHD LSTVT EPIQI MDTLE ATTAP GNDER |
| 9 | DYEAL KAQQD KGKKN GFGKG QVLDP SVLGQ GSMST RIDYA |
| 10 | IDQVS NDSND KKGAS DMPTN TENVL RNRSE GILRQ PEIER |
| 11 | TNSGS ACGSV LSVNS LSNKR SRADD YDCMD EDKEN QPHNP |
| 12 | VPHLV NQTKN ETEED ASETG TIDSH LDRKK KKEGA FKILL |
| 13 | VLYDE GYEDK KSLHF ADKKS IRTGY ASNTY ADDED DDFFD |
| 14 | NKFRD YKESD HYAGG AGGYN NASSM DTTHN DKSTY DSDKH |
| 15 | VKKHA SPDFI FVLSE RIKLP DIDSE DKPEL PRTNS YKFMM |
| 16 | PYTQT YTNPS HCQLC VPSAT SRSAS SASLS QQLQN SPTGC |
| 17 | LWLRE HSKEA PPGAP GAPSS PSELE CSANP QLNDK DPQYV |
| 18 | STESA RSSHD HLNND NRMFT AETSG ITDPA TQANM GNQIT |
| 19 | MNSTL GTSTI PRGLI HRPSA GRTTR PCDNC TCSPG LLSRQ |
| 20 | DAAAA AGLPT TSGSG GGNAG PGAGD EEESG KKTEK DSALM |
| 21 | LLVPE NSRPP VQALP KEYQV RPRTT YEDGP GTPEW KRARL |
| 22 | MFNMN LLSTP SSEEG SPQNR SSSMS SVEGK KDRDT FTNLQ |
| 23 | SNAVA TDTGI SRNTG SNTTK EQKFS AIDAT DSQND GSYGG |
| 24 | TMTSP SGSGR VKNIY NSSNG SPSPS GWDSP SFRNR YNFDD |
| 25 | NTPIA SGILQ PAAFD YFSRP ISTQD IISNN CGNTL SRGPL |
| 26 | AGGAN VSPSS GSTPH ILPSL PTSTS NASSG PPYGY PQPAH |
| 27 | MNIFS QVGGL SPNYD KNFIS NGDVH GRETD DGEFD DSVNM |
| 28 | MSETT NTLRR RTNEE WSATA GIAEQ HENQP SVSTD KKDQS |
| 29 | DDNPK DPKSY DKRDG SNVVD TSKPG DGNQG NDMDW LFRNA |
| 30 | ICSSE FKDNY YQKVE SPTRT PNDWK KNNLL SKNKN TENNK |
| 31 | DLNSK RYSNI PSSKP AGEAL SPVRS HNSGE YRRAD MMTGK |
| 32 | MVSIS ILKGK KKGTE RPIEV THHSY AGGRH EKTKR GTAGV |
| 33 | MSASE AGVTE QVKKL SVKDS SNDAV KPNKK ENKKS KQQSL |
| 34 | MKAFT SLLCG LGLST TLAKA ISLQR PLGLD KDVLL QAAEK |

Table S6: **Details of the IDP dataset used to test MOFF2-pc transferability.** Experimental conditions including temperature $T$ and ionic strength $I$, together with values of $\langle R_g \rangle$, are provided. Citations for the experimental studies are listed in the Ref. column. MOFF2-pc predictions for these testing IDPs were obtained by reweighting the corresponding noise ensembles using the procedure outlined in Ensemble reweighting for model evaluation. Details of the noise-ensemble generation are provided in Table S15.

| Protein | $T$ (K) | $I$ (mM) | $\langle R_g \rangle$ (nm) | Ref. |
|---|---|---|---|---|
| IBB | 300 | 168 | 3.2 | S34 |
| N49 | 300 | 168 | 1.59 | S34 |
| NUS | 300 | 168 | 2.49 | S34 |
| NUL | 300 | 168 | 3.0 | S34 |
| NLS | 300 | 168 | 2.4 | S34 |
| Hst5 | 293 | 150 | 1.38 | S35 |
| SH4UD | 293 | 216 | 2.82 | S36 |
| $(\text{Hst5})_2$ | 298 | 168 | 1.87 | S37 |
| p53 NTD | 277 | 99 | 2.39 | S38 |

Table S7: **Details of the IDP dataset used to evaluate MOFF2-pc transferability and for fine-tuning parameterization by thermodynamic reweighting.** The experimental values of $\langle R_g \rangle$ are provided, together with the condition, including temperature ($T$) and ionic strength ($I$), under which the experiments were performed. Citations for the experimental studies are provided in the Ref. column. To compare against experimental measurements, simulations were performed with the MOFF2-pc model in OpenMM[S13] under identical experimental conditions, i.e., the same $T$ and $I$ values. Each simulation used a 10 fs timestep and lasted $2 \times 10^8$ steps, corresponding to a 2 $\mu$s trajectory. Configurations were saved every $5 \times 10^3$ steps, giving $4 \times 10^4$ snapshots per trajectory. These configurations were also used for fine tuning parameters with thermodynamic reweighting.

| Protein | Type | $T$ (K) | $I$ (mM) | $\langle R_g \rangle$ (nm) | Ref. |
|---|---|---|---|---|---|
| A1-LCD$^{+NLS}$ | IDP | 298.00 | 150 | 2.583 | S39 |
| A1-LCD$^{-NLS}$ | IDP | 298.00 | 150 | 2.760 | S39 |
| A1-LCD$^{-6R+6K}$ | IDP | 298.00 | 150 | 2.787 | S39 |
| A1-LCD$^{-10R+10K}$ | IDP | 298.00 | 150 | 2.849 | S39 |
| A1-LCD$^{-8F+4Y}$ | IDP | 298.00 | 150 | 2.707 | S39 |
| A1-LCD$^{-3R+3K}$ | IDP | 298.00 | 150 | 2.634 | S39 |
| A1-LCD$^{-10R}$ | IDP | 298.00 | 150 | 2.671 | S39 |
| A1-LCD$^{-4D}$ | IDP | 298.00 | 150 | 2.642 | S39 |
| A1-LCD$^{+12D}$ | IDP | 298.00 | 150 | 2.801 | S39 |
| A1-LCD$^{+2R}$ | IDP | 298.00 | 150 | 2.623 | S39 |
| A1-LCD$^{+7K+12D}$ | IDP | 298.00 | 150 | 2.921 | S39 |
| A1-LCD$^{-6R}$ | IDP | 298.00 | 150 | 2.573 | S39 |
| A1-LCD$^{-9F+3Y}$ | IDP | 298.00 | 150 | 2.683 | S39 |
| A1-LCD$^{+8D}$ | IDP | 298.00 | 150 | 2.685 | S39 |
| A1-LCD$^{+12E}$ | IDP | 298.00 | 150 | 2.852 | S39 |
| A1-LCD$^{-12F+12Y}$ | IDP | 298.00 | 150 | 2.604 | S39 |
| A1-LCD$^{+7R}$ | IDP | 298.00 | 150 | 2.709 | S39 |
| A1-LCD$^{+4D}$ | IDP | 298.00 | 150 | 2.718 | S39 |
| A1-LCD$^{+7F-7Y}$ | IDP | 298.00 | 150 | 2.750 | S39 |
| A1-LCD$^{-9F+6Y}$ | IDP | 298.00 | 150 | 2.655 | S39 |

Table S8: **Details of the MDP dataset used to evaluate MOFF2-pc transferability and for fine-tuning parameterization by thermodynamic reweighting.** The experimental values of $\langle R_g \rangle$ are provided, together with the condition, including temperature ($T$) and ionic strength ($I$), under which the experiments were performed. Citations for the experimental studies are provided in the Ref. column. To compare against experimental measurements, simulations were performed with the MOFF2-pc model in OpenMM[S13] under identical experimental conditions, i.e., the same $T$ and $I$ values. Folded regions, reported using residue indices, were restrained to the native conformations during the simulations with the inclusion of the native contact energy $V_{\text{native}}$ (Eq. S12). Each simulation used a 10 fs timestep and lasted $2 \times 10^8$ steps, corresponding to a 2 $\mu$s trajectory. Configurations were saved every $5 \times 10^3$ steps, giving $4 \times 10^4$ snapshots per trajectory. These configurations were also used for fine tuning parameters with thermodynamic reweighting.

| Protein | Type | $T$ (K) | $I$ (mM) | Exp. $\langle R_g \rangle$ (nm) | Folded regions | Ref. |
|---|---|---|---|---|---|---|
| TIA1 | MDP | 293.15 | 100 | 2.75 | 6-82, 95-172, 190-275 | S38 |
| Ubq4 | MDP | 293.00 | 330 | 3.19 | 1-72, 77-148, 153-224, 229-300 | S38 |
| hSUMO-hnRNPA1* | MDP | 293.15 | 100 | 3.37 | 44-114, 132-209, 224-298 | S36 |
| H46 | MDP | 283.00 | 163 | 4.15 | 140-355 | S40 |
| PCPE | MDP | 293.15 | 506 | 4.04 | 12-125, 134-249, 293-412 | S37 |
| NiV-V | MDP | 293.15 | 232 | 6.97 | 406-457 | S35 |
| HeV-V | MDP | 293.15 | 232 | 6.86 | 404-456 | S40 |
| D14 | MDP | 283.15 | 156 | 3.90 | 31-121, 157-246, 265-354, 400-479 | S34 |
| S4FL | MDP | 283.15 | 169 | 4.70 | 15-138, 287-294, 323-466, 492-542 | S38 |
| ChiAM | MDP | 293.15 | 282 | 4.73 | 8-89, 92-172, 178-257, 266-356, 359-462, 471-567, 578-668 | S34 |
| GS0 | MDP | 293.15 | 150 | 3.20 | 1-226, 256-470 | S34 |
| GS24 | MDP | 293.15 | 150 | 3.57 | 1-226, 304-518 | S34 |
| GS32 | MDP | 293.15 | 150 | 3.75 | 1-226, 320-534 | S34 |
| GS48 | MDP | 293.15 | 150 | 4.11 | 1-226, 352-566 | S40 |

Table S9: **Details of the OP training set used for fine-tuning with thermodynamic reweighting.** The proteins shown below were selected from Tables S1 and S2. We estimated the reference value for $\langle R_g \rangle$ from the corresponding atomistic simulations. To compare with atomistic simulations, simulations were performed with the MOFF2-pc model in OpenMM[S13] under identical conditions, i.e., the same temperature (T) and ionic strength (I) values. Each simulation used a 10 fs timestep and lasted $2 \times 10^8$ steps, corresponding to a 2 $\mu$s trajectory. Configurations were saved every $5 \times 10^3$ steps, giving $4 \times 10^4$ snapshots per trajectory. These configurations were used for fine tuning parameters with thermodynamic reweighting.

| Protein | $T$ (K) | $I$ (mM) | Ref. $\langle R_g \rangle$ (nm) |
|---|---|---|---|
| Chignolin | 340.00 | 25.9 | 0.545 |
| Homeodomain | 360.00 | 45 | 1.114 |
| Protein-G | 350.00 | 100 | 1.101 |
| Trp-cage | 290.00 | 65 | 0.854 |
| Villin | 360.00 | 40 | 1.034 |
| WW-domain | 360.00 | 7.1 | 0.974 |
| $\alpha$3D | 300.00 | 3.11 | 1.274 |
| bba | 300.00 | 33.04 | 1.256 |
| bbl | 300.00 | 199.58 | 1.477 |
| engrailed | 300.00 | 280.46 | 1.096 |
| gpw | 300.00 | 36.54 | 1.224 |
| lambda-repressor | 300.00 | 51.04 | 1.220 |
| NTL9 | 300.00 | 121.46 | 1.368 |
| Protein-B | 300.00 | 51.44 | 0.988 |
| BPTI | 300.00 | 37.02 | 1.037 |
| calmodulin | 300.00 | 249.98 | 2.037 |
| GB3 | 300.00 | 9.98 | 1.026 |
| 1soy | 300.00 | 150 | 1.317 |
| 1wla | 300.00 | 150 | 1.504 |
| 2ea9 | 300.00 | 150 | 1.291 |
| 5tvz | 300.00 | 150 | 1.425 |

Table S10: **Details of the IDP testing set for MOFF2 transferability evaluation in Figure 3 of the main text.** The experimental values of $\langle R_g \rangle$ are provided, together with the condition, including temperature ($T$) and ionic strength ($I$), under which the experiments were performed. Citations for the experimental studies are provided in the Ref. column. To compare against experimental measurements, simulations were performed with the MOFF2 model in OpenMM.[S13] Each simulation used a 10 fs timestep and lasted $2 \times 10^8$ steps, corresponding to a 2 $\mu$s trajectory. Configurations were saved every $5 \times 10^3$ steps, giving $4 \times 10^4$ snapshots per trajectory. Only the last $2 \times 10^4$ snapshot configurations were used for analysis.

| Protein | $N$ | $T$ (K) | $I$ (mM) | $\langle R_g \rangle$ (nm) | Ref. |
|---|---|---|---|---|---|
| ANAC046 | 167 | 298.0 | 140.0 | 3.60 | S41 |
| BMAL1P624A | 98 | 283.35 | 154.0 | 2.77 | S42 |
| cDAXX | 246 | 293.0 | 130.0 | 4.75 | S43 |
| ChiZ164 | 67 | 293.0 | 65.0 | 2.42 | S44 |
| D91_FATZ1 | 209 | 293.0 | 180.0 | 3.86 | S45 |
| DomainV | 67 | 288.15 | 198.5 | 2.43 | S46 |
| DSS1 | 71 | 288.0 | 170.0 | 2.50 | S41 |
| ED3 | 373 | 293.15 | 153.0 | 6.51 | S47 |
| ED4 | 163 | 293.15 | 153.0 | 4.06 | S47 |
| GON7 | 114 | 283.0 | 211.0 | 3.18 | S48 |
| hKISS1 | 120 | 283.15 | 159.0 | 3.47 | S49 |
| HvASR1 | 143 | 293.15 | 150.0 | 3.51 | S50 |
| N_FATZ1 | 191 | 293.15 | 192.0 | 3.45 | S45 |
| NHE6cmdd | 116 | 288.0 | 170.0 | 3.20 | S41 |
| p27Cv14 | 107 | 293.0 | 95.0 | 2.94 | S51 |
| p27Cv15 | 107 | 293.0 | 95.0 | 2.92 | S51 |
| p27Cv31 | 107 | 293.0 | 95.0 | 2.81 | S51 |
| p27Cv44 | 107 | 293.0 | 95.0 | 2.49 | S51 |
| p27Cv56 | 107 | 293.0 | 95.0 | 2.33 | S51 |
| p27Cv78 | 107 | 293.0 | 95.0 | 2.21 | S51 |
| PARCL | 180 | 293.15 | 170.0 | 3.43 | S52 |
| PTMA | 111 | 288.0 | 160.0 | 3.70 | S41 |
| TIF2NRID | 150 | 283.15 | 175.0 | 3.74 | S53 |
| TtASR1 | 141 | 293.15 | 150.0 | 3.31 | S50 |
| VWF | 103 | 293.0 | 153.0 | 3.08 | S54 |

Table S11: **Details of the OP testing set for MOFF2 transferability evaluation in Figure 3 of the main text.** Proteins (indicated by their UniProt IDs) were randomly selected with balanced coverage across three chain-length ranges: 0–100, 100–300, and 300–600 residues, and ordered by chain length $N$. Each selected protein had at least 80% of residues with AlphaFold2 pLDDT scores $\geq 80$. Simulations were performed at $T = 298$ K and ionic strength $I = 150$ mM. The reference $\langle R_g \rangle$ was computed from the AlphaFold2-predicted initial structure. Each simulation used a 10 fs timestep and lasted $2 \times 10^8$ steps, corresponding to a 2 $\mu$s trajectory. Configurations were saved every $5 \times 10^3$ steps, giving $4 \times 10^4$ snapshots per trajectory; only the final $2 \times 10^4$ snapshots were used for analysis.

| No. | UniProt ID | $N$ | No. | UniProt ID | $N$ | No. | UniProt ID | $N$ |
|---|---|---|---|---|---|---|---|---|
| 1 | P62945 | 25 | 21 | Q15370 | 118 | 41 | O15121 | 323 |
| 2 | Q9UDW1 | 63 | 22 | O95168 | 129 | 42 | Q9UJ72 | 324 |
| 3 | O00244 | 68 | 23 | P05413 | 133 | 43 | P43235 | 329 |
| 4 | O15239 | 70 | 24 | P62266 | 143 | 44 | Q15165 | 354 |
| 5 | Q969W0 | 71 | 25 | Q9Y547 | 144 | 45 | O15335 | 359 |
| 6 | Q156A1 | 80 | 26 | Q9Y587 | 144 | 46 | P11177 | 359 |
| 7 | A8MT69 | 81 | 27 | Q15819 | 145 | 47 | Q8IUS5 | 362 |
| 8 | O95167 | 84 | 28 | A0A075B759 | 164 | 48 | P06132 | 367 |
| 9 | Q71UM5 | 84 | 29 | A6NEY8 | 169 | 49 | Q8N4Q0 | 377 |
| 10 | P07108 | 87 | 30 | Q96SL4 | 187 | 50 | Q6NVY1 | 386 |
| 11 | Q8N6N7 | 88 | 31 | A0A1W2PQJ5 | 194 | 51 | P17174 | 413 |
| 12 | P63167 | 89 | 32 | Q96S19 | 204 | 52 | Q16773 | 422 |
| 13 | Q96FJ2 | 89 | 33 | Q9NXJ5 | 209 | 53 | Q9HAC7 | 445 |
| 14 | P62072 | 90 | 34 | Q9Y2Q3 | 226 | 54 | Q14728 | 455 |
| 15 | O43504 | 91 | 35 | Q9UIJ7 | 227 | 55 | P16233 | 465 |
| 16 | A6NKH3 | 93 | 36 | P09417 | 244 | 56 | P0DTE8 | 511 |
| 17 | Q8N6V4 | 93 | 37 | P0CG30 | 244 | 57 | P0DTE4 | 527 |
| 18 | O95777 | 96 | 38 | P25786 | 263 | 58 | P06865 | 529 |
| 19 | S4R460 | 96 | 39 | Q17R31 | 274 | 59 | Q99518 | 535 |
| 20 | P14621 | 99 | 40 | P50225 | 295 | 60 | Q9UJ83 | 578 |

Table S12: **Details of the MDP testing set for MOFF2 transferability evaluation in Figure 3 of the main text.** Proteins (indicated by their UniProt IDs) were randomly selected with balanced coverage across three chain-length ranges: 0–100, 100–300, and 300–600 residues. Each selected protein had at least 50% of residues with AlphaFold2 pLDDT scores $\geq 80$ and at least 30% of residues with AlphaFold2 pLDDT scores $\leq 50$. OD ranges were assigned from the initial AlphaFold2 structures for residue pairs within the same continuous secondary structure element (either an $\alpha$-helix or a $\beta$-sheet), as identified using DSSP[S6] implemented in `MDTraj`.[S7] $N_{\mathrm{OD}}/N$ reports the fraction of residues assigned to continuous ordered regions with extra native contacts added for MDP simulations. Across the selected MDP test set, $N_{\mathrm{OD}}/N$ did not exceed 0.75, ensuring that all proteins retained a substantial disordered or flexible fraction. To compare against AlphaFold2-predicted reference structures, simulations were performed with the MOFF2 model in OpenMM[S13] at $T = 298$ K and ionic strength $I = 150$ mM. Each simulation used a 10 fs timestep and lasted $2 \times 10^8$ steps, corresponding to a 2 $\mu$s trajectory. Configurations were saved every $5 \times 10^3$ steps, giving $4 \times 10^4$ snapshots per trajectory. Only the last $2 \times 10^4$ snapshot configurations were used for analysis.

| No. | UniProt ID | OD region(s) | $N$ | $N_{\mathrm{OD}}/N$ |
|---|---|---|---|---|
| 1 | Q9NRI6 | 2-19 | 33 | 0.55 |
| 2 | A0A6I8PS25 | 8-30 | 40 | 0.57 |
| 3 | A0A669KAW2 | 3-24 | 42 | 0.52 |
| 4 | V9GZ13 | 9-31 | 50 | 0.46 |
| 5 | A0A286YFK9 | 14-39 | 51 | 0.51 |
| 6 | P53803 | 15-29, 31-58 | 58 | 0.74 |
| 7 | A0A1B0GTU2 | 14-41 | 59 | 0.47 |
| 8 | P48539 | 31-58 | 62 | 0.45 |
| 9 | H3BU77 | 27-53 | 68 | 0.40 |
| 10 | Q9H4G8 | 6-51, 53-60 | 78 | 0.69 |
| 11 | P0DP73 | 6-12, 28-40, 42-66 | 79 | 0.57 |
| 12 | Q5STR5 | 29-59 | 79 | 0.39 |
| 13 | Q13166 | 35-73 | 79 | 0.49 |
| 14 | Q96FX2 | 4-62 | 82 | 0.72 |
| 15 | A0A096LP55 | 27-90 | 91 | 0.70 |
| 16 | Q07654 | 18-27, 46-87 | 94 | 0.55 |
| 17 | Q5BLP8 | 18-27, 55-88 | 95 | 0.46 |
| 18 | L0R819 | 26-86 | 96 | 0.64 |
| 19 | H0UI37 | 29-95 | 97 | 0.69 |
| 20 | P0DP57 | 22-52, 61-73, 85-96 | 97 | 0.58 |
| 21 | Q02575 | 62-132 | 133 | 0.53 |
| 22 | Q06055 | 67-140 | 141 | 0.52 |
| 23 | Q9BXW4 | 18-43, 55-76, 85-89, 98-120 | 147 | 0.52 |
| 24 | Q12988 | 71-104, 108-109, 111-144 | 150 | 0.47 |
| 25 | P02511 | 68-109, 112-150, 159-164 | 175 | 0.50 |

| No. | UniProt ID | OD region(s) | $N$ | $N_{\mathrm{OD}}/N$ |
|---|---|---|---|---|
| 26 | Q9NZ72 | 42-56, 81-177 | 180 | 0.62 |
| 27 | A6NKQ9 | 56-90, 104-152 | 187 | 0.45 |
| 28 | Q14206 | 11-86, 152-161 | 197 | 0.44 |
| 29 | Q96FZ7 | 14-113, 127-143, 154-161 | 201 | 0.62 |
| 30 | Q9P0W0 | 30-52, 54-62, 66-67, 84-100, 108-131, 161-201 | 207 | 0.56 |
| 31 | P0DML3 | 30-62, 97-123, 135-154, 168-176, 178-215 | 217 | 0.59 |
| 32 | Q12962 | 116-176, 192-211, 216-218 | 218 | 0.39 |
| 33 | Q8TAA5 | 59-121, 147-148, 150-185, 191-220 | 225 | 0.58 |
| 34 | P40259 | 46-67, 73-80, 85-101, 103-125, 134-143, 155-180 | 229 | 0.46 |
| 35 | P23560 | 93-101, 139-149, 152-173, 175-187, 195-244 | 247 | 0.43 |
| 36 | P17931 | 114-250 | 250 | 0.55 |
| 37 | Q9GZT6 | 62-98, 105-106, 108-248 | 254 | 0.71 |
| 38 | Q9Y676 | 50-227, 229-234 | 258 | 0.71 |
| 39 | P25942 | 26-136, 138-188, 198-216 | 277 | 0.65 |
| 40 | Q9BPW8 | 70-173, 175-180, 182-212, 214-283 | 284 | 0.74 |
| 41 | Q12904 | 6-70, 72-73, 149-312 | 312 | 0.74 |
| 42 | Q9UNE2 | 18-19, 47-127, 134-165 | 315 | 0.37 |
| 43 | Q5JRS4 | 26-27, 29-31, 33-34, 36-38, 40-53, 57-77, 79-80, 83-84, 94-168, 181-182, 185-186, 189-233, 235-236, 242-243, 245-246, 248-250 | 329 | 0.55 |
| 44 | Q8N2F6 | 92-291, 303-340 | 343 | 0.69 |
| 45 | Q14106 | 1-115 | 344 | 0.33 |
| 46 | Q9GZM5 | 128-138, 152-169, 185-239, 242-294 | 350 | 0.39 |
| 47 | Q7L513 | 75-148, 154-200, 203-261 | 359 | 0.50 |
| 48 | O14904 | 56-60, 64-106, 109-152, 168-183, 190-217, 227-253, 288-344, 348-365 | 365 | 0.65 |
| 49 | Q5M8T2 | 9-38, 40-99, 104-123, 129-146, 155-181, 185-207, 216-269, 281-304 | 416 | 0.62 |
| 50 | P47972 | 70-72, 81-83, 164-203, 205-206, 225-395, 398-420 | 431 | 0.56 |
| 51 | Q9BTV6 | 4-39, 61-72, 78-123, 128-151, 162-330, 332-343, 431-450 | 452 | 0.71 |
| 52 | Q9Y5X2 | 62-63, 65-82, 93-133, 144-189, 194-268, 285-359, 369-438 | 465 | 0.70 |
| 53 | Q86X02 | 22-141, 189-258, 353-373, 436-437, 439-454 | 465 | 0.49 |
| 54 | Q13505 | 153-304, 311-365, 368-392, 421-436 | 466 | 0.53 |
| 55 | Q99928 | 48-56, 58-110, 114-204, 220-278, 281-305, 307-340, 444-464 | 467 | 0.63 |
| 56 | Q96HE7 | 50-86, 103-105, 108-110, 176-188, 199-200, 249-269, 275-294, 297-324, 329-330, 333-354, 385-415, 435-464 | 468 | 0.45 |
| 57 | O14757 | 8-11, 22-28, 30-41, 52-208, 210-268, 379-385, 387-406, 411-418, 423-430, 437-441, 450-466 | 476 | 0.64 |
| 58 | P98170 | 28-37, 43-74, 78-80, 82-90, 159-160, 163-176, 179-211, 214-220, 222-224, 258-293, 295-302, 306-311, 314-324, 327-334, 375-393, 406-417, 446-485, 487-494 | 497 | 0.53 |
| 59 | Q15750 | 21-38, 43-57, 62-90, 97-135, 147-369 | 504 | 0.64 |
| 60 | Q8NHP7 | 77-232, 234-280, 303-332 | 514 | 0.45 |

Table S13: **Comparison of simulated and experimental saturation concentrations for condensates formed by A1-LCD variants shown in Figure 4 of the main text.** Results obtained at $T = 293$ K were used to calculate deviations from experiment in units of M for comparison with the Mpipi-recharged model.[S56]

| Protein | Temperature (K) | MOFF2 $C_{\mathrm{sat}}$ ($\mu$M) | Exp. $C_{\mathrm{sat}}$ ($\mu$M) | Ref. |
|---|---|---|---|---|
| A1-LCD$^{+12D}$ | 293 | 90.8 | 529.3 | S39 |
| A1-LCD$^{+7K+12D}$ | 293 | 472.0 | 270.4 | S39 |
| A1-LCD$^{+7R+10D}$ | 293 | 20.9 | 21.0 | S39 |
| A1-LCD$^{+8D}$ | 293 | 100.2 | 138.7 | S39 |
| A1-LCD$^{+NLS}$ | 293 | 47.4 | 73.4 | S39 |
| A1-LCD$^{-10G+10S}$ | 293 | 312.5 | 268.1 | S39 |
| A1-LCD$^{-12F+12Y}$ | 293 | 29.8 | 61.0 | S39 |
| A1-LCD$^{-2K}$ | 293 | 7.6 | 14.1 | S39 |
| A1-LCD$^{-3R+3K}$ | 293 | 433.9 | 595.1 | S39 |
| A1-LCD$^{-8F+4Y}$ | 293 | 423.4 | 695.7 | S39 |
| A1-LCD$^{-9F+6Y}$ | 293 | 88.0 | 341.8 | S39 |
| A1-LCD$^{allW}$ | 293 | 55.5 | 0.23 | S55 |
| A1-LCD$^{FtoW}$ | 293 | 194.4 | 0.79 | S55 |
| A1-LCD$^{YtoW}$ | 293 | 84.9 | 14.6 | S55 |
| A1-LCD$^{W-}$ | 293 | 473.0 | 37.9 | S55 |
| A1-LCD$^{+12E}$ | 285 | 467.7 | 478.1 | S39 |
| A1-LCD$^{+2R}$ | 289 | 80.2 | 95.5 | S39 |
| A1-LCD$^{-NLS}$ | 297 | 154.6 | 146.1 | S39 |
| A1-LCD$^{+7F-7Y}$ | 298 | 468.1 | 452.8 | S39 |
| A1-LCD$^{+7R+12D}$ | 298 | 33.8 | 15.7 | S39 |
| A1-LCD$^{-6R}$ | 298 | 25.6 | 62.1 | S39 |

Table S14: **Details of the umbrella simulations used to generate OP noise ensembles for potential contrasting and evaluation of the ensemble-reweighting method.** The simulations used RMSD-based umbrella biases relative to the reference structures. The number of umbrella windows, biasing centers, and restraining constant $\kappa$ are listed below. Gray-highlighted proteins were later used to compare noise-ensemble reweighting against direct MD simulations with the MOFF2-pc model, as shown in Figure S9.

| Protein | No. windows | Umbrella centers (nm) | $\kappa$ (kJ/mol/nm$^2$) |
|---|---|---|---|
| α3d | 7 | [0.0, 0.5, 1.0, 1.5, 2.0, 2.5, 3.0] | 2000.0 |
| bba | 7 | [0.0, 0.5, 1.0, 1.5, 2.0, 2.5, 3.0] | 2000.0 |
| bbl | 7 | [0.0, 0.5, 1.0, 1.5, 2.0, 2.5, 3.0] | 2000.0 |
| engrailed | 7 | [0.0, 0.5, 1.0, 1.5, 2.0, 2.5, 3.0] | 2000.0 |
| gpw | 7 | [0.0, 0.5, 1.0, 1.5, 2.0, 2.5, 3.0] | 2000.0 |
| lambda-repressor | 7 | [0.0, 0.5, 1.0, 1.5, 2.0, 2.5, 3.0] | 2000.0 |
| NTL9 | 7 | [0.0, 0.5, 1.0, 1.5, 2.0, 2.5, 3.0] | 2000.0 |
| Protein-B | 7 | [0.0, 0.5, 1.0, 1.5, 2.0, 2.5, 3.0] | 2000.0 |
| BPTI | 7 | [0.0, 0.5, 1.0, 1.5, 2.0, 2.5, 3.0] | 2000.0 |
| calmodulin | 7 | [0.0, 0.5, 1.0, 1.5, 2.0, 2.5, 3.0] | 2000.0 |
| GB3 | 7 | [0.0, 0.5, 1.0, 1.5, 2.0, 2.5, 3.0] | 2000.0 |
| Hen-Egg-White-Lysozyme | 7 | [0.0, 0.5, 1.0, 1.5, 2.0, 2.5, 3.0] | 2000.0 |
| Chignolin | 7 | [0.0, 0.5, 1.0, 1.5, 2.0, 2.5, 3.0] | 2000.0 |
| Trp-cage | 7 | [0.0, 0.5, 1.0, 1.5, 2.0, 2.5, 3.0] | 2000.0 |
| Villin | 7 | [0.0, 0.5, 1.0, 1.5, 2.0, 2.5, 3.0] | 2000.0 |
| WW-domain | 7 | [0.0, 0.5, 1.0, 1.5, 2.0, 2.5, 3.0] | 2000.0 |
| Homeodomain | 7 | [0.0, 0.5, 1.0, 1.5, 2.0, 2.5, 3.0] | 2000.0 |
| Protein-G | 7 | [0.0, 0.5, 1.0, 1.5, 2.0, 2.5, 3.0] | 2000.0 |
| 1soy_clean | 7 | [0.00, 0.05, 0.10, 0.15, 0.20, 0.25, 0.30] | 2000.0 |
| 1wla_clean | 9 | [0.00, 0.05, 0.10, 0.15, 0.20, 0.25, 0.30, 0.35, 0.40] | 2000.0 |
| 2ea9_clean | 6 | [0.00, 0.05, 0.10, 0.15, 0.20, 0.25] | 2000.0 |
| 5tvz_clean | 5 | [0.00, 0.05, 0.10, 0.15, 0.20] | 2000.0 |
| Ubiquitin | 7 | [0.0, 0.5, 1.0, 1.5, 2.0, 2.5, 3.0] | 2000.0 |
| AF-P00251-F1-model_v4_w_H | 7 | [0.00, 0.05, 0.10, 0.15, 0.20, 0.25, 0.30] | 2000.0 |
| AF-P0C232-F1-model_v4_w_H | 7 | [0.00, 0.05, 0.10, 0.15, 0.20, 0.25, 0.30] | 2000.0 |
| AF-P0CG98-F1-model_v4_w_H | 11 | [0.0, 0.1, 0.2, 0.3, 0.4, 0.5, 0.6, 0.7, 0.8, 0.9, 1.0] | 500.0 |
| AF-P21149-F1-model_v4_w_H | 7 | [0.0, 0.1, 0.2, 0.3, 0.4, 0.5, 0.6] | 500.0 |
| AF-P21318-F1-model_v4_w_H | 8 | [0.00, 0.05, 0.10, 0.15, 0.20, 0.25, 0.30, 0.35] | 2000.0 |
| AF-P29669-F1-model_v4_w_H | 6 | [0.0, 0.1, 0.2, 0.3, 0.4, 0.5] | 500.0 |
| AF-P31960-F1-model_v4_w_H | 11 | [0.0, 0.1, 0.2, 0.3, 0.4, 0.5, 0.6, 0.7, 0.8, 0.9, 1.0] | 500.0 |
| AF-P32729-F1-model_v4_w_H | 7 | [0.00, 0.05, 0.10, 0.15, 0.20, 0.25, 0.30] | 2000.0 |
| AF-P61734-F1-model_v4_w_H | 8 | [0.00, 0.05, 0.10, 0.15, 0.20, 0.25, 0.30, 0.35] | 2000.0 |
| AF-P69995-F1-model_v4_w_H | 7 | [0.00, 0.05, 0.10, 0.15, 0.20, 0.25, 0.30] | 2000.0 |
| AF-P75202-F1-model_v4_w_H | 8 | [0.0, 0.1, 0.2, 0.3, 0.4, 0.5, 0.6, 0.7] | 500.0 |
| AF-P75459-F1-model_v4_w_H | 8 | [0.00, 0.05, 0.10, 0.15, 0.20, 0.25, 0.30, 0.35] | 2000.0 |
| AF-P80353-F1-model_v4_w_H | 9 | [0.00, 0.05, 0.10, 0.15, 0.20, 0.25, 0.30, 0.35, 0.40] | 2000.0 |
| AF-P87285-F1-model_v4_w_H | 7 | [0.0, 0.1, 0.2, 0.3, 0.4, 0.5, 0.6] | 500.0 |

Table S15: **Details of the umbrella simulations used to generate IDP noise ensembles for potential contrasting and evaluation of the ensemble-reweighting method.** The simulations used $R_g$-based umbrella biases. The number of umbrella windows, biasing centers, and restraining constant $\kappa$ are listed below. Gray-highlighted A1-LCD variants were later used to compare noise-ensemble reweighting against direct MD simulations with the MOFF2-pc model, as shown in Figure S9. The PC training and PC test labels differentiate proteins used for training model with potential contrasting in Figure 2A and testing model performance in Figure S3.

| Protein | Dataset role | No. windows | Umbrella centers (nm) | $\kappa$ (kJ/mol/nm$^2$) |
|---|---|---|---|---|
| A$\beta$40 | PC training | 7 | [0.0, 0.5, 1.0, 1.5, 2.0, 2.5, 3.0] | 20.0 |
| ACTR | PC training | 8 | [0.0, 0.5, 1.0, 1.5, 2.0, 2.5, 3.0, 3.5] | 20.0 |
| Ash1 | PC training | 11 | [0.0, 0.5, 1.0, 1.5, 2.0, 2.5, 3.0, 3.5, 4.0, 4.5, 5.0] | 20.0 |
| drkN-SH3 | PC training | 8 | [0.0, 0.5, 1.0, 1.5, 2.0, 2.5, 3.0, 3.5] | 20.0 |
| $N_{tail}$ | PC training | 11 | [0.0, 0.5, 1.0, 1.5, 2.0, 2.5, 3.0, 3.5, 4.0, 4.5, 5.0] | 20.0 |
| p15PAF | PC training | 11 | [0.0, 0.5, 1.0, 1.5, 2.0, 2.5, 3.0, 3.5, 4.0, 4.5, 5.0] | 20.0 |
| Sic1 | PC training | 11 | [0.0, 0.5, 1.0, 1.5, 2.0, 2.5, 3.0, 3.5, 4.0, 4.5, 5.0] | 20.0 |
| $IDP_{WZ}$ (34) | PC training | 11 | [0.0, 0.5, 1.0, 1.5, 2.0, 2.5, 3.0, 3.5, 4.0, 4.5, 5.0] | 20.0 |
| IBB | PC test | 10 | [0.0, 0.5, 1.0, 1.5, 2.0, 2.5, 3.0, 3.5, 4.0, 4.5] | 20.0 |
| NLS | PC test | 10 | [0.0, 0.5, 1.0, 1.5, 2.0, 2.5, 3.0, 3.5, 4.0, 4.5] | 20.0 |
| N49 | PC test | 10 | [0.0, 0.5, 1.0, 1.5, 2.0, 2.5, 3.0, 3.5, 4.0, 4.5] | 20.0 |
| NUS | PC test | 10 | [0.0, 0.5, 1.0, 1.5, 2.0, 2.5, 3.0, 3.5, 4.0, 4.5] | 20.0 |
| NUL | PC test | 10 | [0.0, 0.5, 1.0, 1.5, 2.0, 2.5, 3.0, 3.5, 4.0, 4.5] | 20.0 |
| SH4UD | PC test | 9 | [0.0, 0.5, 1.0, 1.5, 2.0, 2.5, 3.0, 3.5, 4.0] | 20.0 |
| Hst5 | PC test | 8 | [0.0, 0.5, 1.0, 1.5, 2.0, 2.5, 3.0, 3.5] | 20.0 |
| $(Hst5)_2$ | PC test | 8 | [0.0, 0.5, 1.0, 1.5, 2.0, 2.5, 3.0, 3.5] | 20.0 |
| p53 NTD | PC test | 10 | [0.0, 0.5, 1.0, 1.5, 2.0, 2.5, 3.0, 3.5, 4.0, 4.5] | 20.0 |
| A1-LCD$^{-10R+10K}$ | Reweighting vs. MD | 10 | [0.0, 0.5, 1.0, 1.5, 2.0, 2.5, 3.0, 3.5, 4.0, 4.5] | 20.0 |
| A1-LCD$^{-6R+6K}$ | Reweighting vs. MD | 10 | [0.0, 0.5, 1.0, 1.5, 2.0, 2.5, 3.0, 3.5, 4.0, 4.5] | 20.0 |
| A1-LCD$^{+12D}$ | Reweighting vs. MD | 10 | [0.0, 0.5, 1.0, 1.5, 2.0, 2.5, 3.0, 3.5, 4.0, 4.5] | 20.0 |
| A1-LCD$^{+12E}$ | Reweighting vs. MD | 10 | [0.0, 0.5, 1.0, 1.5, 2.0, 2.5, 3.0, 3.5, 4.0, 4.5] | 20.0 |
| A1-LCD$^{+7K+12D}$ | Reweighting vs. MD | 10 | [0.0, 0.5, 1.0, 1.5, 2.0, 2.5, 3.0, 3.5, 4.0, 4.5] | 20.0 |
| A1-LCD$^{-NLS}$ | Reweighting vs. MD | 10 | [0.0, 0.5, 1.0, 1.5, 2.0, 2.5, 3.0, 3.5, 4.0, 4.5] | 20.0 |
| A1-LCD$^{-12F+12Y}$ | Reweighting vs. MD | 10 | [0.0, 0.5, 1.0, 1.5, 2.0, 2.5, 3.0, 3.5, 4.0, 4.5] | 20.0 |
| A1-LCD$^{+7F-7Y}$ | Reweighting vs. MD | 10 | [0.0, 0.5, 1.0, 1.5, 2.0, 2.5, 3.0, 3.5, 4.0, 4.5] | 20.0 |
| A1-LCD$^{+7R}$ | Reweighting vs. MD | 10 | [0.0, 0.5, 1.0, 1.5, 2.0, 2.5, 3.0, 3.5, 4.0, 4.5] | 20.0 |

Table S16: **Details of the umbrella simulations used to generate MDP noise ensembles for potential contrasting.** The simulations used biases on the $R_g$, with the number of umbrella windows, the biasing centers, and the restraining constant $\kappa$ listed below.

| Protein | No. windows | Umbrella centers (nm) | $\kappa$ (kJ/mol/nm$^2$) |
|---|---|---|---|
| THB-C2 | 4 | [1.5, 2.0, 2.5, 3.0] | 20.0 |
| Ub2 | 4 | [1.5, 2.0, 2.5, 3.0] | 20.0 |
| Ub3 | 5 | [1.5, 2.0, 2.5, 3.0, 3.5] | 20.0 |
| Gal3 | 6 | [2.0, 2.5, 3.0, 3.5, 4.0, 4.5] | 20.0 |
| hnRNPA1* | 6 | [2.0, 2.5, 3.0, 3.5, 4.0, 4.5] | 20.0 |
| FPs-GS$_8$ | 7 | [2.0, 2.5, 3.0, 3.5, 4.0, 4.5, 5.0] | 20.0 |
| FPs-GS$_{16}$ | 7 | [2.0, 2.5, 3.0, 3.5, 4.0, 4.5, 5.0] | 20.0 |
| SH4UD-SH3-SH2 | 5 | [2.0, 2.5, 3.0, 3.5, 4.0] | 20.0 |
| TDP43$_{\mathrm{WtoA}}$ | 7 | [2.5, 3.0, 3.5, 4.0, 4.5, 5.0, 5.5] | 20.0 |
| D12 | 6 | [1.5, 2.0, 2.5, 3.0, 3.5, 4.0] | 20.0 |
| D23 | 6 | [1.5, 2.0, 2.5, 3.0, 3.5, 4.0] | 20.0 |
| D34 | 6 | [1.5, 2.0, 2.5, 3.0, 3.5, 4.0] | 20.0 |
| SMAD4 | 7 | [2.5, 3.0, 3.5, 4.0, 4.5, 5.0, 5.5] | 20.0 |

### Supplemental Figures

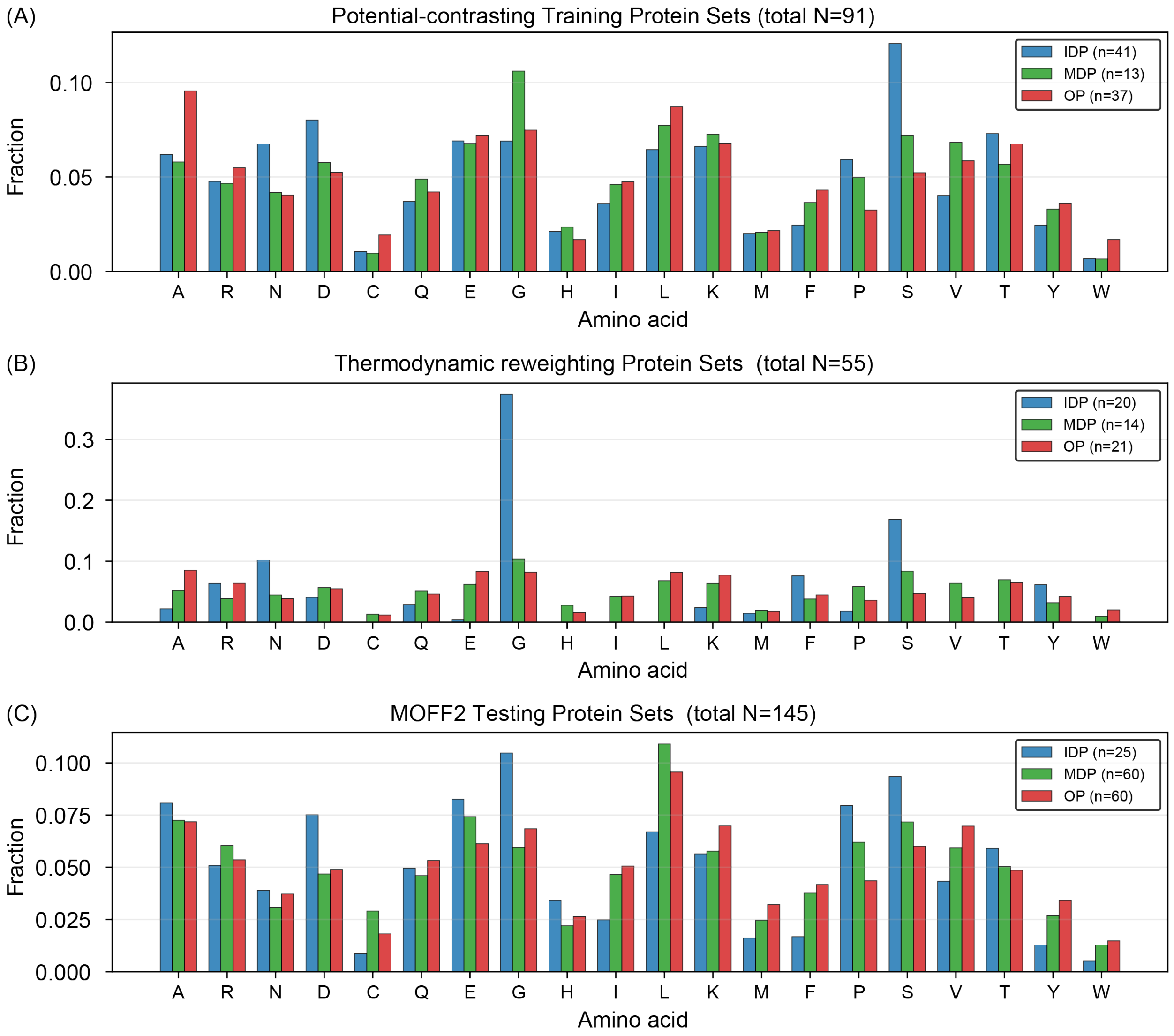


Figure S1: **Amino acid distribution for proteins included in potential contrasting (A), fine-tuning (B), and MOFF2 testing (C).** More details regarding the proteins can be found in Tables S1–S4 for panel A, (Tables S7–S9) for panel B, and (Tables S10–S12) for panel C.

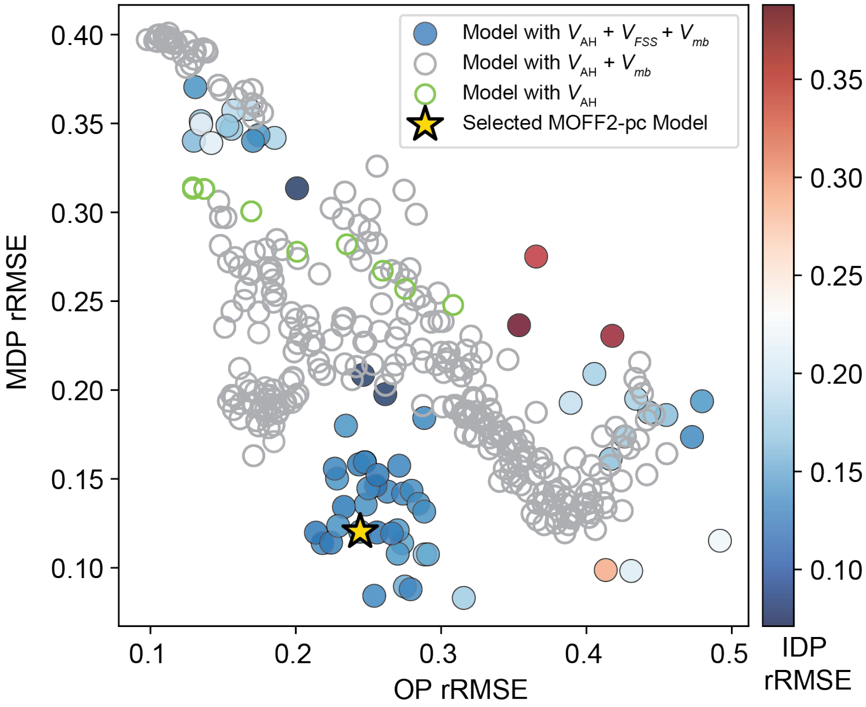


Figure S2: **Hyperparameter scanning for Potential Contrasting.** As detailed in Potential Contrasting, the hyperparameters were selected by evaluating the performance of the resulting optimized model on the training set proteins through a thermodynamic reweighting procedure without performing MD simulations. The model performance is quantified by the relative root mean squared errors (rRMSE) between simulation and reference values of $R_g$ for ordered, multi-domain, and intrinsically disordered proteins, as indicated in the $x$-axis, $y$-axis, and the colorbar. Each point represents a model with a set of interaction parameters optimized under a specific hyperparameter combination. Green open circles correspond to models trained with only the pairwise contact potential ($V_{\mathrm{AH}}$), gray open circles correspond to models trained with $V_{\mathrm{AH}}$ and the density-dependent many-body potential ($V_{\mathrm{mb}}$), and filled circles correspond to models with the full potential.

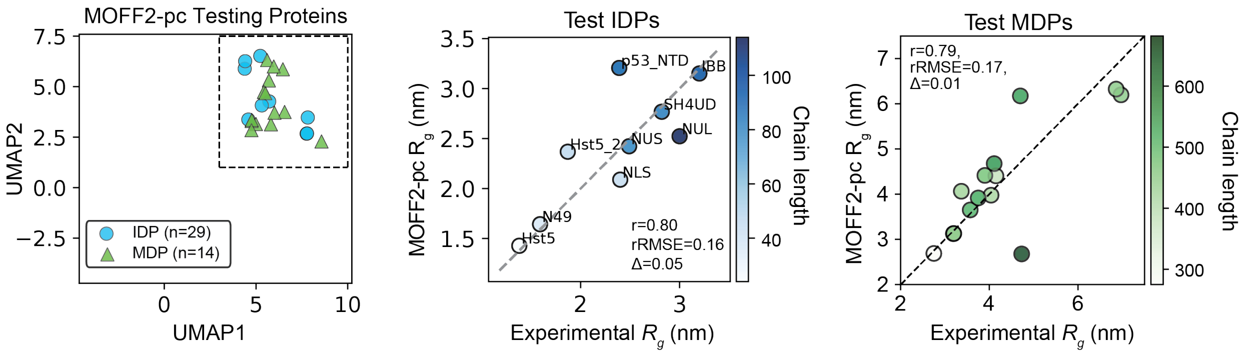


Figure S3: **Evaluating MOFF2-pc on proteins unseen during training.** The left panel shows the UMAP projection of the testing proteins, including 9 IDPs and 14 MDPs. Simulation and experimental details for the results shown in the right two panels, together with protein identities, are provided in Tables S6 and S8.

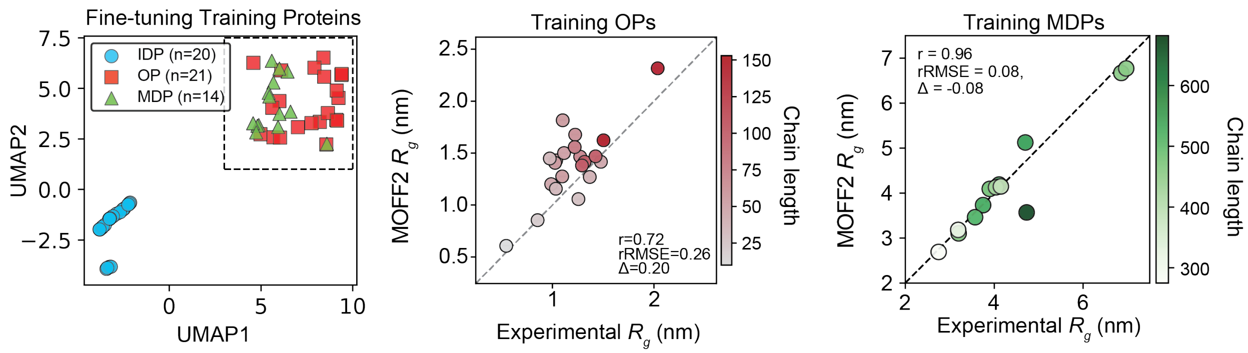


Figure S4: **Evaluating MOFF2 on proteins used for fine tuning with thermodynamic reweighting.** The left panel shows the UMAP projection of the proteins using for fine tuning. Simulation and experimental details for the results shown in the right two panels, together with protein identities, are provided in Tables S9 and S8.

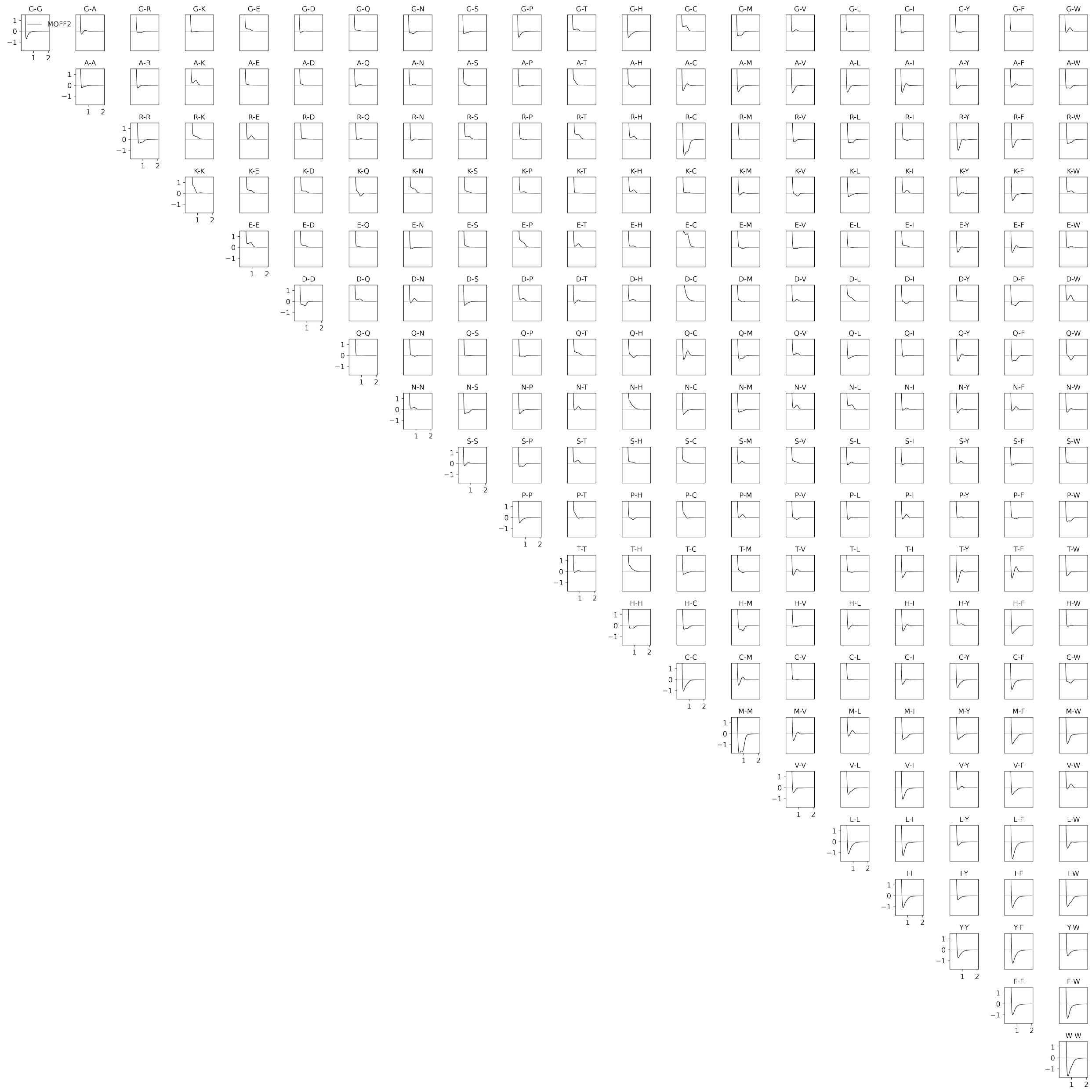


Figure S5: **MOFF2 residue-specific pair interactions ($V_{\mathbf{pair}}$) that combine the Ashbaugh-Hatch (AH) potential with a Gaussian first-solvation-shell correction as defined in Eq. S5.**

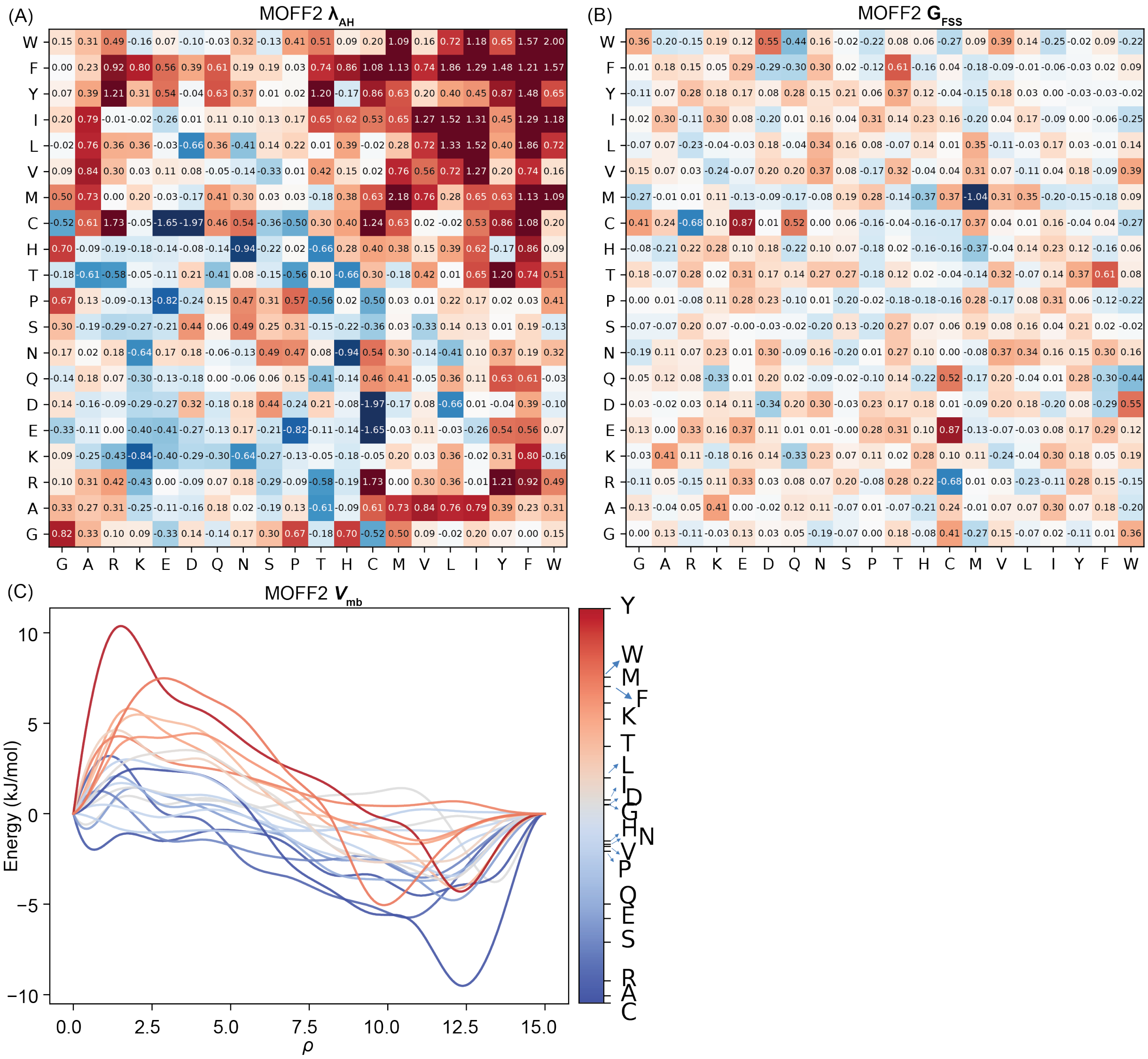


Figure S6: **MOFF2 energy profiles.** (A) Learned MOFF2 AH contact matrix, $\lambda_{AH}$, as defined in Eq. S6. (B) Learned MOFF2 first-solvation-shell Gaussian matrix, $G_{FSS}$, as defined in Eq. S8. (C) Learned MOFF2 density-dependent many-body potential, $V_{mb}$, as defined in Eq. S11.

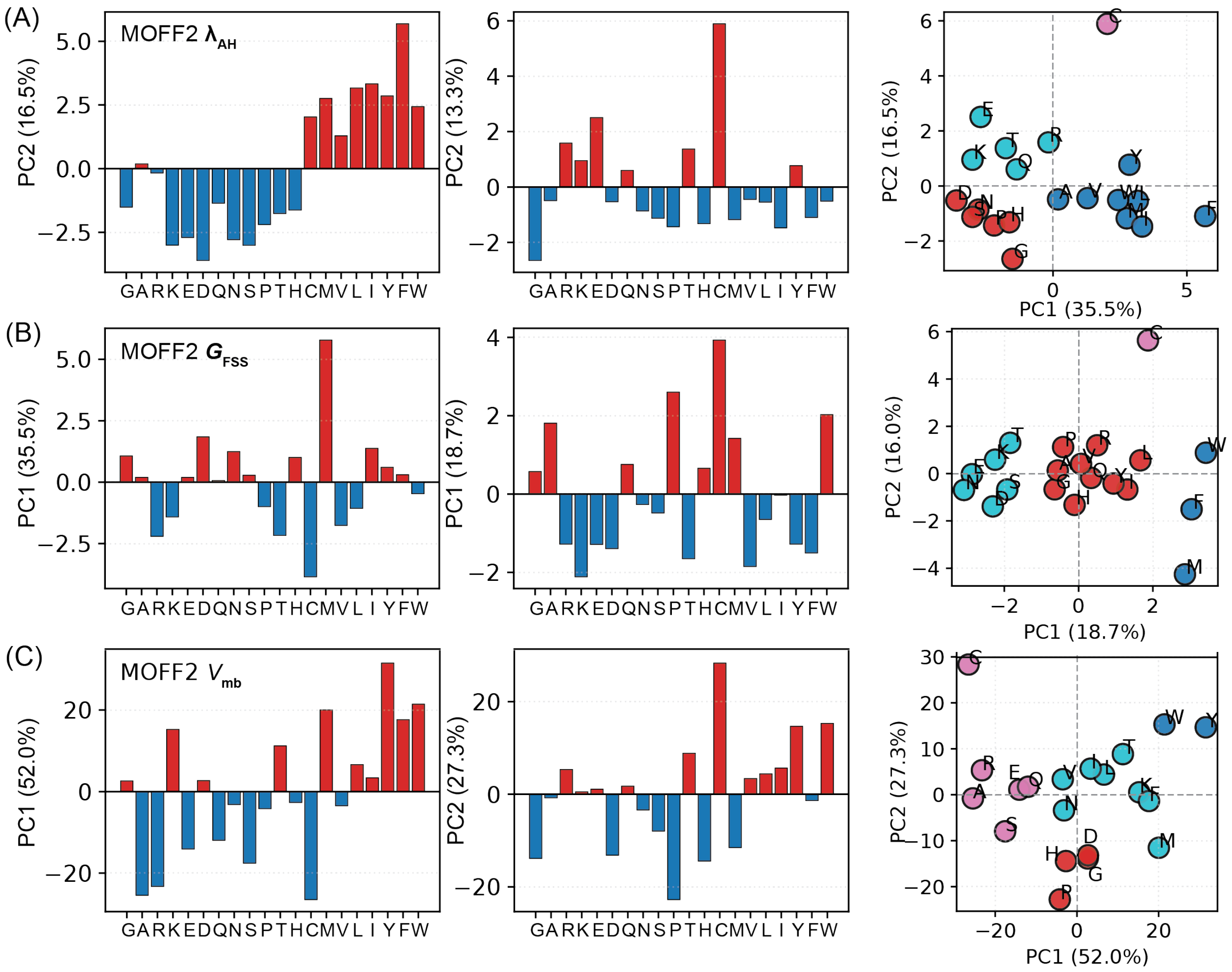


Figure S7: **Principal-component decomposition of residue-level parameter profiles.** (A)-(C) Projection of amino acids onto the first two principal components of the learned MOFF2 $\lambda_{\mathrm{AH}}$, $G_{\mathrm{FSS}}$, and $V_{\mathrm{mb}}$ profiles, respectively. Clustering based on the first two PCs is shown on the right, with colors indicating the cluster assignments obtained from $k$-means[S57] clustering in the PC1–PC2 space.

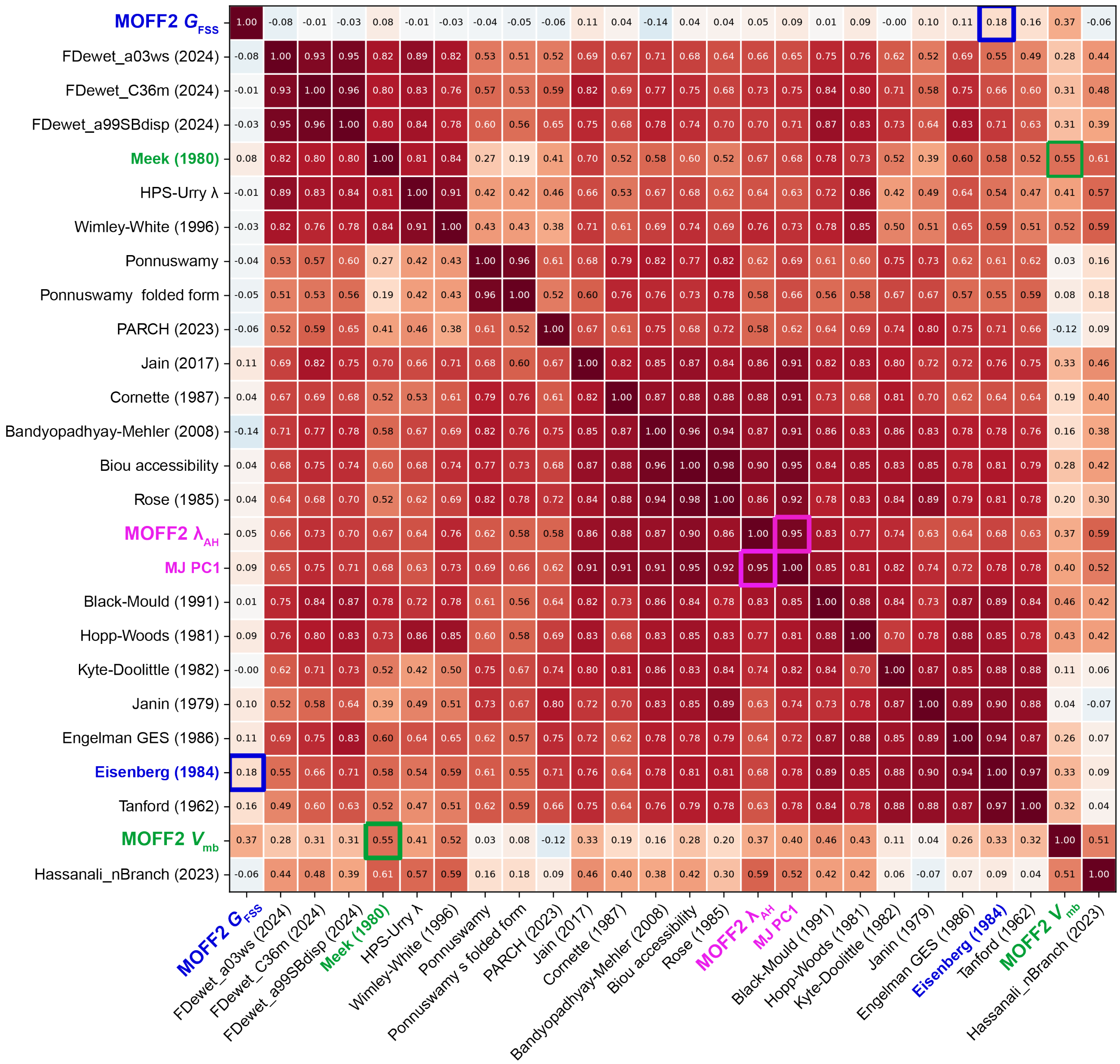


Figure S8: **Correlation between MOFF2 learned residue profiles and existing hydrophobicity scales.** The colorscale and numerical values indicate the Pearson correlation coefficients, computed using the first principal-component (PC1) residue profiles of MOFF2 learned parameters, as presented in Figure S7. The squares highlight the most correlated hydrophobicity scales.

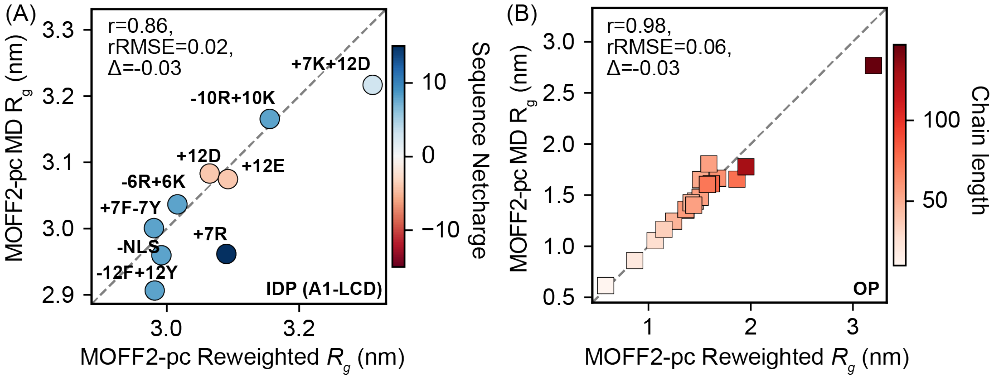


Figure S9: **Consistency between $\langle R_g \rangle$ values estimated from thermodynamic reweighting and those from direct MD simulations.** The proteins shown in parts A and B are listed in Table S15 and Table S14, respectively. The agreement between the two sets of values supports the use of generalized-ensemble reweighting for rapid hyperparameter screening.

#### References


(S1) Ashbaugh, H. S.; Hatch, H. W. Natively unfolded protein stability as a coil-to-globule transition in charge/hydropathy space. *Journal of the American Chemical Society* **2008**, *130*, 9536–9542.

(S2) Regy, R. M.; Thompson, J.; Kim, Y. C.; Mittal, J. Improved coarse-grained model for studying sequence dependent phase separation of disordered proteins. *Protein Science* **2021**, *30*, 1371–1379.

(S3) Hastie, T.; Tibshirani, R.; Friedman, J. H.; Friedman, J. H. *The elements of statistical learning: data mining, inference, and prediction*; Springer, 2009; Vol. 2.

(S4) Noel, J. K.; Whitford, P. C.; Onuchic, J. N. The shadow map: a general contact definition for capturing the dynamics of biomolecular folding and function. *The journal of physical chemistry B* **2012**, *116*, 8692–8702.

(S5) Liu, S.; Wang, C.; Latham, A. P.; Ding, X.; Zhang, B. OpenABC enables flexible, simplified, and efficient GPU accelerated simulations of biomolecular condensates. *PLoS Computational Biology* **2023**, *19*, e1011442.

(S6) Kabsch, W.; Sander, C. Dictionary of protein secondary structure: pattern recognition of hydrogen-bonded and geometrical features. *Biopolymers: Original Research on Biomolecules* **1983**, *22*, 2577–2637.

(S7) McGibbon, R. T.; Beauchamp, K. A.; Harrigan, M. P.; Klein, C.; Swails, J. M.; Hernández, C. X.; Schwantes, C. R.; Wang, L.-P.; Lane, T. J.; Pande, V. S. MDTraj: a modern open library for the analysis of molecular dynamics trajectories. *Biophysical journal* **2015**, *109*, 1528–1532.

(S8) Robustelli, P.; Piana, S.; Shaw, D. E. Developing a molecular dynamics force field for both folded and disordered protein states. *Proceedings of the National Academy of Sciences* **2018**, *115*, E4758–E4766.

(S9) Piana, S.; Robustelli, P.; Tan, D.; Chen, S.; Shaw, D. E. Development of a force field for the simulation of single-chain proteins and protein–protein complexes. *Journal of chemical theory and computation* **2020**, *16*, 2494–2507.

(S10) Lindorff-Larsen, K.; Piana, S.; Dror, R. O.; Shaw, D. E. How fast-folding proteins fold. *Science* **2011**, *334*, 517–520.

(S11) Jumper, J.; Evans, R.; Pritzel, A.; Green, T.; Figurnov, M.; Ronneberger, O.; Tunyasuvunakool, K.; Bates, R.; Žídek, A.; Potapenko, A.; others Highly accurate protein structure prediction with AlphaFold. *nature* **2021**, *596*, 583–589.

(S12) Varadi, M.; Anyango, S.; Deshpande, M.; Nair, S.; Natassia, C.; Yordanova, G.; Yuan, D.; Stroe, O.; Wood, G.; Laydon, A.; others AlphaFold Protein Structure Database: massively expanding the structural coverage of protein-sequence space with high-accuracy models. *Nucleic acids research* **2022**, *50*, D439–D444.

(S13) Eastman, P.; Swails, J.; Chodera, J. D.; McGibbon, R. T.; Zhao, Y.; Beauchamp, K. A.; Wang, L.-P.; Simmonett, A. C.; Harrigan, M. P.; Stern, C. D.; others OpenMM 7: Rapid development of high performance algorithms for molecular dynamics. *PLoS computational biology* **2017**, *13*, e1005659.

(S14) Zhang, Z.; Liu, X.; Yan, K.; Tuckerman, M. E.; Liu, J. Unified efficient thermostat scheme for the canonical ensemble with holonomic or isokinetic constraints via molecular dynamics. *The Journal of Physical Chemistry A* **2019**, *123*, 6056–6079.

(S15) Wang, C.; Zhang, B. Sequence-Dependent Conformational Landscapes of Intrinsically Disordered Proteins Reveal Asymmetric Chain Compaction. *Journal of Chemical Theory and Computation* **2025**, *21*, 11282–11292.

(S16) Virtanen, P.; Gommers, R.; Oliphant, T. E.; Haberland, M.; Reddy, T.; Cournapeau, D.; Burovski, E.; Peterson, P.; Weckesser, W.; Bright, J.; others SciPy 1.0: fundamental algorithms for scientific computing in Python. *Nature methods* **2020**, *17*, 261–272.

(S17) Ding, X. Optimizing force fields with experimental data using ensemble reweighting and potential contrasting. *The Journal of Physical Chemistry B* **2024**, *128*, 6760–6769.

(S18) Matsunaga, Y.; Kamiya, M.; Oshima, H.; Jung, J.; Ito, S.; Sugita, Y. Use of Multistate Bennett Acceptance Ratio Method for Free-Energy Calculations from Enhanced Sampling and Free-Energy Perturbation. *Biophysical Reviews* **2022**, *14*, 1503–1512.

(S19) Ding, X.; Vilseck, J. Z.; Brooks, C. L. I. Fast Solver for Large Scale Multistate Bennett Acceptance Ratio Equations. *Journal of Chemical Theory and Computation* **2019**, *15*, 799–802.

(S20) Thaler, S.; Zavadlav, J. Learning Neural Network Potentials from Experimental Data via Differentiable Trajectory Reweighting. *Nature Communications* **2021**, *12*, 6884.

(S21) Riveros, I.; Zhang, B. NEAT-DNA: A Chemically Accurate, Sequence-Dependent Coarse-Grained Model for Large-Scale DNA Simulations. *Journal of Chemical Theory and Computation* **2026**, *22*, 3709–3719.

(S22) Zwanzig, R. W. High-temperature equation of state by a perturbation method. I. Nonpolar gases. *The Journal of Chemical Physics* **1954**, *22*, 1420–1426.

(S23) Airas, J.; Zhang, B. Scaling Graph Neural Networks to Large Proteins. *arXiv preprint arXiv:2410.03921* **2024**,

(S24) Michie, K. A.; Kwan, A. H.; Tung, C.-S.; Guss, J. M.; Trewhella, J. A highly conserved yet flexible linker is part of a polymorphic protein-binding domain in myosin-binding protein C. *Structure* **2016**, *24*, 2000–2007.

(S25) Jussupow, A.; Messias, A. C.; Stehle, R.; Geerlof, A.; Solbak, S. M.; Paissoni, C.; Bach, A.; Sattler, M.; Camilloni, C. The dynamics of linear polyubiquitin. *Science advances* **2020**, *6*, eabc3786.

(S26) Lin, Y.-H.; Qiu, D.-C.; Chang, W.-H.; Yeh, Y.-Q.; Jeng, U.-S.; Liu, F.-T.; Huang, J.-r. The intrinsically disordered N-terminal domain of galectin-3 dynamically mediates multisite self-association of the protein through fuzzy interactions. *Journal of Biological Chemistry* **2017**, *292*, 17845–17856.

(S27) Martin, E. W.; Thomasen, F. E.; Milkovic, N. M.; Cuneo, M. J.; Grace, C. R.; Nourse, A.; Lindorff-Larsen, K.; Mittag, T. Interplay of folded domains and the disordered low-complexity domain in mediating hnRNPA1 phase separation. *Nucleic acids research* **2021**, *49*, 2931–2945.

(S28) Moses, D.; Guadalupe, K.; Yu, F.; Flores, E.; Perez, A. R.; McAnelly, R.; Shamoon, N. M.; Kaur, G.; Cuevas-Zepeda, E.; Merg, A. D.; others Structural biases in disordered proteins are prevalent in the cell. *Nature Structural & Molecular Biology* **2024**, *31*, 283–292.

(S29) Gurumoorthy, V.; Shrestha, U. R.; Zhang, Q.; Pingali, S. V.; Boder, E. T.; Urban, V. S.; Smith, J. C.; Petridis, L.; O'Neill, H. Disordered Domain Shifts the Conformational Ensemble of the Folded Regulatory Domain of the Multidomain Oncoprotein c-Src. *Biomacromolecules* **2023**, *24*, 714–723.

(S30) Wright, G. S.; Watanabe, T. F.; Amporndanai, K.; Plotkin, S. S.; Cashman, N. R.; Antonyuk, S. V.; Hasnain, S. S. Purification and structural characterization of aggregation-prone human TDP-43 involved in neurodegenerative diseases. *Iscience* **2020**, *23*.

(S31) Hajizadeh, N. R.; Pieprzyk, J.; Skopintsev, P.; Flayhan, A.; Svergun, D. I.; Löw, C. Probing the architecture of a multi-PDZ domain protein: Structure of PDZK1 in solution. *Structure* **2018**, *26*, 1522–1533.

(S32) Gomes, T.; Martin-Malpartida, P.; Ruiz, L.; Aragón, E.; Cordeiro, T. N.; Macias, M. J. Conformational landscape of multidomain SMAD proteins. *Computational and Structural Biotechnology Journal* **2021**, *19*, 5210–5224.

(S33) Cao, F.; von Bülow, S.; Tesei, G.; Lindorff-Larsen, K. A coarse-grained model for disordered and multi-domain proteins. *Protein Science* **2024**, *33*, e5172.

(S34) Fuertes, G.; Banterle, N.; Ruff, K. M.; Chowdhury, A.; Mercadante, D.; Koehler, C.; Kachala, M.; Estrada Girona, G.; Milles, S.; Mishra, A.; others Decoupling of size and shape fluctuations in heteropolymeric sequences reconciles discrepancies in SAXS vs. FRET measurements. *Proceedings of the National Academy of Sciences* **2017**, *114*, E6342–E6351.

(S35) Jephthah, S.; Staby, L.; Kragelund, B.; Skepo, M. Temperature dependence of intrinsically disordered proteins in simulations: What are we missing? *Journal of chemical theory and computation* **2019**, *15*, 2672–2683.

(S36) Arbesú, M.; Maffei, M.; Cordeiro, T. N.; Teixeira, J. M.; Pérez, Y.; Bernadó, P.; Roche, S.; Pons, M. The unique domain forms a fuzzy intramolecular complex in Src family kinases. *Structure* **2017**, *25*, 630–640.

(S37) Fagerberg, E.; Månsson, L. K.; Lenton, S.; Skepö, M. The effects of chain length on the structural properties of intrinsically disordered proteins in concentrated solutions. *The Journal of Physical Chemistry B* **2020**, *124*, 11843–11853.

(S38) Zhao, J.; Blayney, A.; Liu, X.; Gandy, L.; Jin, W.; Yan, L.; Ha, J.-H.; Canning, A. J.; Connelly, M.; Yang, C.; others EGCG binds intrinsically disordered N-terminal domain of p53 and disrupts p53-MDM2 interaction. *Nature communications* **2021**, *12*, 986.

(S39) Bremer, A.; Farag, M.; Borcherds, W. M.; Peran, I.; Martin, E. W.; Pappu, R. V.; Mittag, T. Deciphering How Naturally Occurring Sequence Features Impact the Phase Behaviours of Disordered Prion-like Domains. *Nature Chemistry* **2022**, *14*, 196–207.

(S40) Mylonas, E.; Hascher, A.; Bernado, P.; Blackledge, M.; Mandelkow, E.; Svergun, D. I. Domain conformation of tau protein studied by solution small-angle X-ray scattering. *Biochemistry* **2008**, *47*, 10345–10353.

(S41) Pesce, F.; Newcombe, E. A.; Seiffert, P.; Tranchant, E. E.; Olsen, J. G.; Grace, C. R.; Kragelund, B. B.; Lindorff-Larsen, K. Assessment of Models for Calculating the Hydrodynamic Radius of Intrinsically Disordered Proteins. *Biophysical Journal* **2023**, *122*, 310–321.

(S42) Garg, A.; Orru, R.; Ye, W.; Distler, U.; Chojnacki, J. E.; Köhn, M.; Tenzer, S.; Sönnichsen, C.; Wolf, E. Structural and Mechanistic Insights into the Interaction of the Circadian Transcription Factor BMAL1 with the KIX Domain of the CREB-binding Protein. *Journal of Biological Chemistry* **2019**, *294*, 16604–16619.

(S43) Schmit, J. D.; Bouchard, J. J.; Martin, E. W.; Mittag, T. Protein Network Structure Enables Switching between Liquid and Gel States. *Journal of the American Chemical Society* **2020**, *142*, 874–883.

(S44) Hicks, A.; Escobar, C. A.; Cross, T. A.; Zhou, H.-X. Sequence-Dependent Correlated Segments in the Intrinsically Disordered Region of ChiZ. *Biomolecules* **2020**, *10*, 946.

(S45) Sponga, A.; Arolas, J. L.; Schwarz, T. C.; Jeffries, C. M.; Rodriguez Chamorro, A.; Kostan, J.; Ghisleni, A.; Drepper, F.; Polyansky, A.; De Almeida Ribeiro, E.; Pedron, M.; Zawadzka-Kazimierczuk, A.; Mlynek, G.; Peterbauer, T.; Doto, P.;

Schreiner, C.; Hollerl, E.; Mateos, B.; Geist, L.; Faulkner, G.; Kozminski, W.; Svergun, D. I.; Warscheid, B.; Zagrovic, B.; Gautel, M.; Konrat, R.; Djinović-Carugo, K. Order from Disorder in the Sarcomere: FATZ Forms a Fuzzy but Tight Complex and Phase-Separated Condensates with $\alpha$-Actinin. *Science Advances* **2021**, *7*, eabg7653.

(S46) Chan-Yao-Chong, M.; Deville, C.; Pinet, L.; van Heijenoort, C.; Durand, D.; Ha-Duong, T. Structural Characterization of N-WASP Domain V Using MD Simulations with NMR and SAXS Data. *Biophysical Journal* **2019**, *116*, 1216–1227.

(S47) Gondelaud, F.; Bouakil, M.; Le Fèvre, A.; Miele, A. E.; Chirot, F.; Duclos, B.; Liwo, A.; Ricard-Blum, S. Extended Disorder at the Cell Surface: The Conformational Landscape of the Ectodomains of Syndecans. *Matrix Biology Plus* **2021**, *12*, 100081.

(S48) Arrondel, C.; Missoury, S.; Snoek, R.; Patat, J.; Menara, G.; Collinet, B.; Liger, D.; Durand, D.; Gribouval, O.; Boyer, O.; Buscara, L.; Martin, G.; Machuca, E.; Nevo, F.; Lescop, E.; Braun, D. A.; Boschat, A.-C.; Sanquer, S.; Guerrera, I. C.; Revy, P.; Parisot, M.; Masson, C.; Boddaert, N.; Charbit, M.; Decramer, S.; Novo, R.; Macher, M.-A.; Ranchin, B.; Bacchetta, J.; Laurent, A.; Collardeau-Frachon, S.; van Eerde, A. M.; Hildebrandt, F.; Magen, D.; Antignac, C.; van Tilbeurgh, H.; Mollet, G. Defects in t6A tRNA Modification Due to GON7 and YRDC Mutations Lead to Galloway-Mowat Syndrome. *Nature Communications* **2019**, *10*, 3967.

(S49) de Opakua, A. I.; Merino, N.; Villate, M.; Cordeiro, T. N.; Ormaza, G.; Sánchez-Carbayo, M.; Diercks, T.; Bernadó, P.; Blanco, F. J. The Metastasis Suppressor KISS1 Is an Intrinsically Disordered Protein Slightly More Extended than a Random Coil. *PLOS ONE* **2017**, *12*, e0172507.

(S50) Hamdi, K.; Salladini, E.; O'Brien, D. P.; Brier, S.; Chenal, A.; Yacoubi, I.; Longhi, S.

Structural Disorder and Induced Folding within Two Cereal, ABA Stress and Ripening (ASR) Proteins. *Scientific Reports* **2017**, *7*, 15544.

(S51) Das, R. K.; Huang, Y.; Phillips, A. H.; Kriwacki, R. W.; Pappu, R. V. Cryptic Sequence Features within the Disordered Protein p27Kip1 Regulate Cell Cycle Signaling. *Proceedings of the National Academy of Sciences* **2016**, *113*, 5616–5621.

(S52) Ostendorp, A.; Ostendorp, S.; Zhou, Y.; Chaudron, Z.; Wolffram, L.; Rombi, K.; von Pein, L.; Falke, S.; Jeffries, C. M.; Svergun, D. I.; Betzel, C.; Morris, R. J.; Kragler, F.; Kehr, J. Intrinsically Disordered Plant Protein PARCL Colocalizes with RNA in Phase-Separated Condensates Whose Formation Can Be Regulated by Mutating the PLD. *Journal of Biological Chemistry* **2022**, *298*, 102631.

(S53) Senicourt, L.; le Maire, A.; Allemand, F.; Carvalho, J. E.; Guee, L.; Germain, P.; Schubert, M.; Bernadó, P.; Bourguet, W.; Sibille, N. Structural Insights into the Interaction of the Intrinsically Disordered Co-activator TIF2 with Retinoic Acid Receptor Heterodimer (RXR/RAR). *Journal of Molecular Biology* **2021**, *433*, 166899.

(S54) del Amo-Maestro, L.; Sagar, A.; Pompach, P.; Goulas, T.; Scavenius, C.; Ferrero, D. S.; Castrillo-Briceño, M.; Taulés, M.; Enghild, J. J.; Bernadó, P.; Gomis-Rüth, F. X. An Integrative Structural Biology Analysis of Von Willebrand Factor Binding and Processing by ADAMTS-13 in Solution. *Journal of Molecular Biology* **2021**, *433*, 166954.

(S55) Alshareedah, I.; Borcherds, W. M.; Cohen, S. R.; Singh, A.; Posey, A. E.; Farag, M.; Bremer, A.; Strout, G. W.; Tomares, D. T.; Pappu, R. V.; Mittag, T.; Banerjee, P. R. Sequence-Specific Interactions Determine Viscoelasticity and Ageing Dynamics of Protein Condensates. *Nature Physics* **2024**, *20*, 1482–1491.

(S56) Pedraza, E.; Tejedor, A. R.; Feito, A.; Gámez, F.; Collepardo-Guevara, R.; Sanz, E.; Espinosa, J. R. Predicting Saturation Concentrations of Phase-Separating Proteins via

Thermodynamic Integration. *Journal of Chemical Theory and Computation* **2025**, *21*, 9919–9934.

(S57) Jin, X.; Han, J. *Encyclopedia of Machine Learning*; Springer, Boston, MA, 2011; pp 563–564.
